# Dysregulation of epithelial differentiation architecture underlies field cancerization in head and neck cancer

**DOI:** 10.64898/2026.10.12.757511

**Authors:** Claire Mills, Yin Huang, Chan Mi Lee, Connor Finkbeiner, Rachel K. Lex, Dominik J. Otto, Ruijin Yang, Hoku West-Foyle, Daniel Jones, Annalyssa Long, Sam Minot, Emma Nguyen, Pushpa Itagi, Samantha Schuster, Anna Elz, Lena Schroeder, Neal Futran, Emily Marchiano, Rocco Ferrandino, Rosa Ana Risques, Gavin Ha, Anshu Bandhlish, Brittany Barber, Manu Setty, Slobodan Beronja

**Affiliations:** Human Biology Division, Fred Hutchinson Cancer Center, Seattle WA; Department of Oral Health Sciences, University of Washington, Seattle WA; Basic Sciences Division, Fred Hutchinson Cancer Center, Seattle WA; Herbold Computational Biology Program, Public Health Sciences Division, Fred Hutchinson Cancer Center, Seattle WA; Molecular and Cellular Biology Program, University of Washington, Seattle WA; Genome Sciences Department, University of Washington, Seattle WA; Cellular Imaging Shared Resource, Fred Hutchinson Cancer Center, Seattle WA; Vaccine and Infectious Disease Division, Fred Hutchinson Cancer Center, Seattle WA; Immunotherapy Integrated Research Center, Fred Hutchinson Cancer Center, Seattle WA; Data Core, Fred Hutchinson Cancer Center, Seattle WA; Department of Otolaryngology-Head and Neck Surgery, University of Washington School of Medicine, Seattle; Department of Laboratory Medicine and Pathology, University of Washington, Seattle, WA

## Abstract

Head and neck squamous cell carcinoma (HNSCC) has exceptionally high recurrence rates, exceeding 50%, contributing to mortality and morbidity. This recurrence rate is proposed to be due to field cancerization, a phenomenon where tumor-adjacent “field” tissue that appears histologically normal harbors molecular changes that prime it for subsequent tumor growth. Current models attribute field cancerization to shared driver mutations between tumors and adjacent at-risk tissue, but this assumption has never been directly tested. To comprehensively characterize this phenomenon, we profiled patient-matched tumor, normal-adjacent field (NAF), contralateral normal, and blood samples from HPV-negative HNSCC patients using whole genome sequencing, targeted driver gene sequencing, single-nucleus multiome profiling, spatial transcriptomics and functional proliferation assays. While histologically normal NAF tissues did harbor expanded mutant clones, these mutations rarely overlapped with paired tumors and were not shared across patients, suggesting molecular features beyond mutations are altered in NAF. Single-cell multiomics revealed that NAF epithelium loses its normal organizational hierarchy: basal stem cell markers persist in differentiated cells, while terminal differentiation programs fail to activate, with reconfiguration of both gene regulatory networks and cell-cell interactions. Proliferating cells appeared in tissue layers normally restricted to post-mitotic cells, a pattern typically seen only in overt dysplasia. Within this disrupted hierarchy, a distinct epithelial state arose in NAF that expressed both normal epithelial and tumor-associated markers, which we localized to the parabasal layers using immunofluorescence and spatial transcriptomics. Together, our findings demonstrate that field cancerization operates through epithelial hierarchy disorganization that disrupts differentiation, generates cells with tumor-like features, and may create a permissive environment for the expansion of cancer-associated mutations, all within tissue that appears histologically normal.

## Introduction

Field cancerization, a phenomenon first described over seventy years ago, presents a fundamental challenge in oncology: tissues that surround malignant tumors appear histologically normal yet harbor molecular alterations that predispose them to malignant transformation^1–3^. These pre-malignant changes may represent the earliest events in tumorigenesis, occurring before any histological abnormalities become apparent. Additionally, as this tissue appears normal, it is often not excised during surgical tumor removal and may contribute to treatment failure.

HPV-negative head and neck squamous cell carcinoma (HNSCC) is among the most aggressive epithelial cancers, with patients frequently presenting with late-stage disease and experiencing local recurrence rates that exceed 50% within two years of treatment^4–10^. Field cancerization has long been implicated in HNSCC, where previous studies have identified genetic changes, including rare *TP53* mutations and chromosomal alterations, in tumor-free tissue margins^9–13^. The prevailing model interprets these findings as evidence that a stepwise accumulation of mutations drives normal tissue toward malignancy^3,10,11^. However, whether these mutations represent clonal spread from adjacent tumors or independent expansion within a permissive tissue environment remains unresolved. To distinguish between these possibilities, we profiled patient-matched tumor, tumor-proximal normal-adjacent field (NAF), contralateral normal (CN), and blood samples from HPV-negative HNSCC patients.

Using whole genome sequencing and targeted cancer-associated gene sequencing, we found that NAF tissues harbored expanded mutant clones at higher variant allele frequencies than contralateral normal tissues, yet these mutations rarely overlapped with those driving the adjacent tumors and were not consistent across patients. WGS similarly revealed neither elevated mutation burden nor chromosomal alterations in NAF relative to CN. These results argue against a shared-mutation model of field cancerization and instead suggest that the tumor-adjacent field is permissive to clonal expansion, consistent with a breakdown in the tissue’s capacity to constrain mutant clones.

To identify what changes in cellular state or tissue organization might underlie this permissive environment, we performed paired single-nucleus RNA+ATAC-seq on 22 matched tumor, NAF, and CN tissues from eight patients, providing single-cell resolution of both transcriptomic and chromatin accessibility profiles. We also developed a computational approach integrating both modalities to characterize cell-cell signaling changes in the microenvironment.

Our analysis revealed three key features of field cancerization in histologically normal NAF tissues. First, the epithelial differentiation hierarchy was disorganized: basal progenitor cell-associated markers persisted in differentiated suprabasal layers, and cells retained proliferative capacity in compartments normally restricted to post-mitotic cells. Second, signaling networks in the NAF microenvironment were reprogramed, with differentiation-promoting signals shifting toward proliferative and epithelial-to-mesenchymal transition pathways. Third, we identified an “emergent” epithelial cell state enriched in NAF that shares transcriptional programs with tumor cells and may represent an intermediate between normal epithelium and malignancy. We validated that these transcriptional changes were reflected in altered protein expression patterns in an independent patient cohort and further confirmed these findings using spatial transcriptomics. Together, these findings demonstrate that field cancerization operates through regulatory reprogramming that disrupts epithelial differentiation, gives rise to cells with tumor-like molecular features, and may create conditions permissive to the expansion of cancer-associated mutant clones, all within tissue that appears histologically normal.

## Results

### Patient-matched tissue sampling enables comprehensive investigation of field cancerization

To comprehensively characterize field cancerization, we designed a prospective study collecting fresh matched tissues from HPV-negative, treatment-naïve HNSCC patients undergoing surgical tumor excision (**Fig. 1A, Suppl. Tables S1, S2**). In addition to tumor (T), we collected normal-adjacent field (NAF) tissues located 0.1–1 cm from the tumor edge. This NAF tissue serves as our experimental sample for studying field cancerization, as it is likely to harbor early molecular alterations predisposing it to malignant transformation^9–13^. Critically, we also collected contralateral normal (CN) tissues from the opposite side of the oral cavity (~2–10 cm from tumor). This distant tissue provides a within-patient control that accounts for patient-specific factors including age and environmental exposures while serving as our best approximation of truly normal oral epithelium from each patient. Matched peripheral blood was also collected from each patient and used as a germline reference to distinguish somatic mutations from inherited variants. Each tissue was reviewed by a head and neck pathologist to confirm diagnoses and ensure NAF tissues were histologically normal. This comprehensive design allowed us to rigorously test the existing model of field cancerization while also enabling detection of non-genetic alterations that might define the cancerized field.

**Figure 1.**
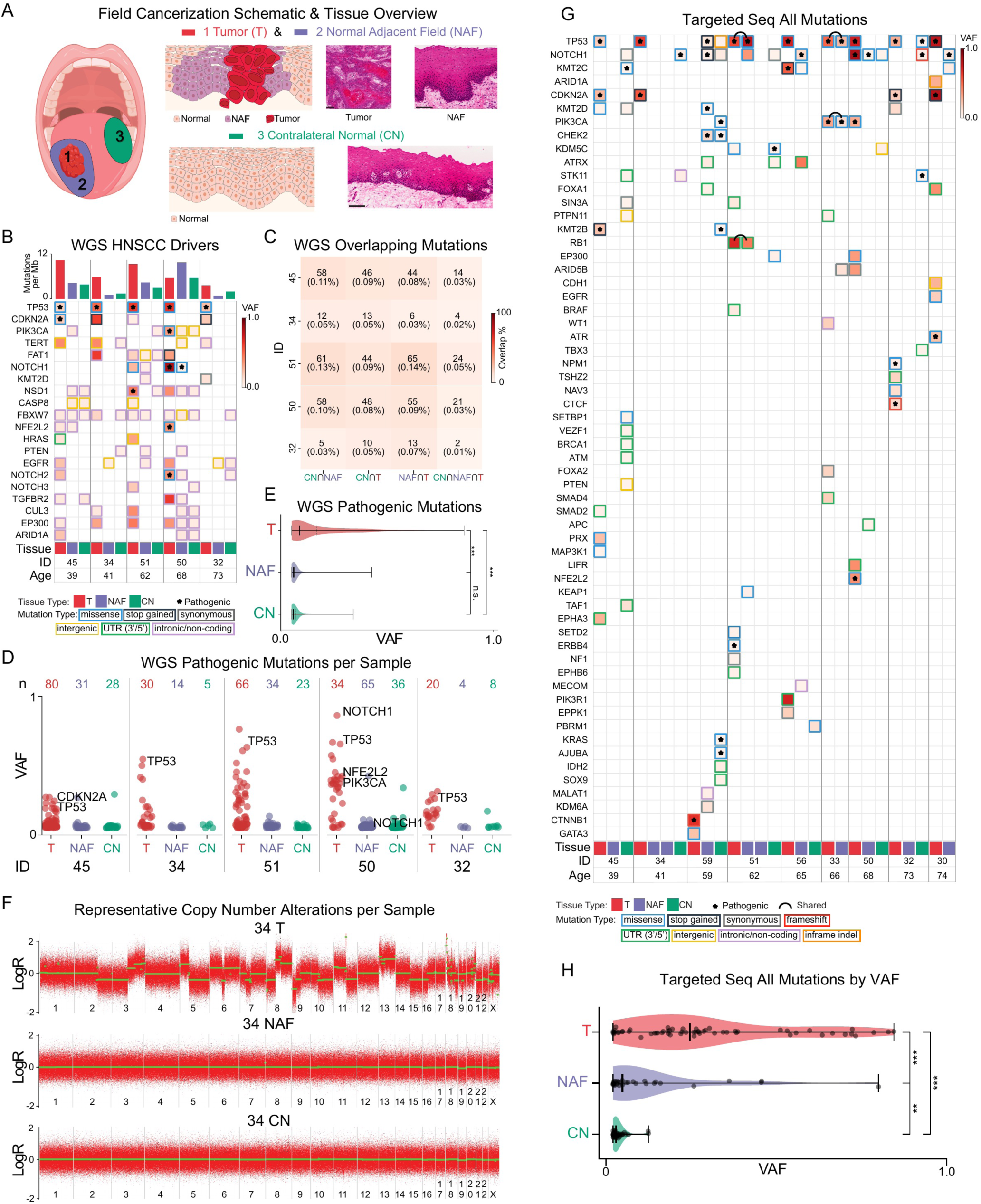
HNSCC-adjacent normal epithelium harbors expanded mutant clones. (**A**) Schematic of tissue collection strategy. Matched tissues were collected from HPV-negative HNSCC patients: tumor (T, red), normal-adjacent field (NAF, purple, 100 μm–1 cm from tumor margin), and contralateral normal (CN, green, approximately 2–10 cm from tumor). Representative H&E staining shows histology of each tissue type. Scale bars indicate 100 µm. (**B**) Whole genome sequencing of laser capture microdissected epithelial cells from five patients (n = 20 samples, including matched blood for each patient). Top: mutation burden (mutations per Mb). Bottom: variant allele frequency (VAF) of mutations in canonical HNSCC driver genes. Mutation types indicated by outline color; known pathogenic variants marked with an asterisk. (**C**) Overlap of somatic mutations between tissue types within each patient. Numbers indicate shared mutations with percentage of total patient mutations in parentheses. (**D**) VAF of AlphaMissense predicted pathogenic missense variants across tissue types for each patient. Numbers above each column indicate variant count per tissue. Select cancer-associated genes are labeled. (**E**) Aggregate VAF distribution of predicted pathogenic variants by tissue type. Thick solid lines indicate median; thin dashed lines indicate mean. Statistical significance determined by two-sided Mann-Whitney U test followed by Bonferroni correction; ***p < 0.001, n.s. not significant (T vs NAF, p < 3e-21; T vs CN, p < 6e-18; NAF vs CN, p = 1.0). (**F**) Representative copy number profiles for patient 34 showing log_2_ ratio across chromosomes. NAF and CN tissues maintain diploid genomes while tumor displays chromosomal alterations. (**G**) Targeted sequencing of cancer-associated genes across nine patients (n = 36 samples). VAF of detected mutations shown by color intensity. Mutation type indicated by outline color; known pathogenic variants marked with stars; mutations shared between tissues within a patient indicated by connecting arcs. (**H**) VAF distribution from targeted sequencing by tissue type. Statistical significance determined by two-sided Mann-Whitney U test followed by Bonferroni correction; ***p < 0.001, **p < 0.01 (T vs CN, p < 3e⁻11; T vs NAF, p = 7e-5; NAF vs CN, p = 8e-3).

### HNSCC-adjacent normal epithelium harbors expanded mutant clones

To test whether field cancerization in HNSCC reflects shared mutations between tumors and adjacent tissue or independent clonal expansion within the field, we performed comprehensive genomic profiling of patient-matched tumor (T), normal-adjacent field (NAF), contralateral normal (CN), and blood samples. We first investigated the mutational landscape in these tissues using whole genome sequencing (WGS) of these tissues from 5 patients (n = 20 samples). We isolated epithelial cells (mean ~25,000 cells per sample) from tissues by laser capture microdissection to enhance detection of rare variants (**Suppl. Fig. 1A**). Samples were sequenced to ~72X mean depth for tissues (~40X for blood). We identified mutations across the genome in all tissue samples, with generally higher burden in tumors (**Fig. 1B, top)**. Mutational signature analysis revealed that normal tissues across all patients were dominated by clock-like signatures (SBS1, SBS5) (**Suppl. Fig. 1B**). Tumors showed additional contributions from APOBEC-associated mutagenesis (SBS2, SBS13), consistent with established HNSCC mutational processes^14^. DNA repair deficiency signatures (SBS3, SBS30) were notably enriched in a subset of tumors.

While we identified HNSCC-associated mutations^15^ across tumors in our cohort, including those in *TP53*, *CDKN2A*, *PIK3CA*, and *NOTCH1*, mutations detected in NAF and CN were predominantly non-coding or synonymous, with mutation burden being largely consistent between NAF and CN (**Fig. 1B, Suppl. Fig. 1C, Suppl. Table S3**). Examination of variant allele frequencies (VAF) further supported the absence of a mutational gradient between NAF and tumor (**Suppl. Fig. 1D**). VAF distributions showed that NAF and CN mutations occurred at low frequencies, whereas tumors displayed both low and high VAF mutations distributed bimodally, consistent with a subset of mutations undergoing clonal expansion. The vast majority of mutations were sample-specific, with overlapping mutations between NAF and tumor being rare (**Fig. 1C**). To assess whether NAF tissues harbor an enrichment of damaging mutations beyond known HNSCC-associated cancer drivers, we utilized AlphaMissense^16^ to identify predicted pathogenic mutations. We found that NAF and CN showed similar low frequency distributions of pathogenic variants that are predicted to affect protein structure or function and were rarely in cancer-associated genes (**Fig. 1D, E**).

We next assessed whether NAF tissues harbored chromosomal copy number alterations. Copy number analysis demonstrated that tumors harbored extensive chromosomal alterations, including amplifications, deletions, and loss of heterozygosity, which is consistent with previous reports for HNSCC^15^, including frequent gains of 3q, 11q, 7p, 8q, and 5p and losses of 9p, 3p, and 8p (**Fig. 1F, Suppl. Fig. 2, Suppl. Table S4**). In contrast, both NAF and CN maintained stable, diploid genomes.

Taken together, WGS data did not reveal consistent genomic differences between NAF and CN: both tissue types harbored scattered mutations at low allele frequencies, lacked chromosomal instability, and were dominated by the same endogenous mutational signatures. This pattern is consistent with somatic mosaicism, in which endogenous mutational processes generate clonal patches throughout normal epithelium regardless of proximity to a tumor^17,18^.

The sequencing depth of WGS may limit detection of rare, expanded clones. To address this, we performed deeper targeted sequencing of 127 cancer-associated genes (**Suppl. Table S5**) in an expanded cohort of 9 patients (n = 36 tissues; mean ~12,400 epithelial cells per sample) at ~2,300X raw depth with unique molecular identifier (UMI)-based error correction, yielding ~900X deduplicated coverage and enabling variant detection at 2% VAF. Consistent with WGS, tumors harbored mutations in canonical HNSCC drivers including *TP53*, *NOTCH1*, and *PIK3CA*, while NAF and CN displayed sporadic mutations across various cancer-associated genes (**Fig. 1G, Suppl. Fig. 3A**). Targeted sequencing, however, revealed a key difference that WGS lacked the sensitivity to detect: NAF tissues harbored cancer-associated mutations at significantly higher allele frequencies than CN (**Fig. 1H**; Bonferroni corrected p < 9e-3), indicating clonal expansion specifically in the tumor-adjacent field. To evaluate whether these expanded NAF clones derive from adjacent tumors, we identified the subset of reads that support tumor-identified mutations in matched NAF, CN, and blood samples. The majority of tumor mutations had no corresponding reads in NAF or CN with a few exceptions (**Suppl. Fig. 3B, Fig. 1G**, arcs). This confirms that positive clonal expansion in NAF is largely driven by mutations acquired independently of the adjacent tumor.

These findings suggest that the tumor-adjacent field may harbor somatically acquired mutations that, unlike in the truly normal epithelium, undergo clonal expansion, potentially reflecting non-genetic features of the tumor-adjacent tissues. These genetic changes may create permissive conditions for positive selection of cancer-associated mutations.

### Single-cell multiomics characterizes field cancerization in HNSCC

To identify the cellular and tissue-level changes that may underlie field cancerization and clonal expansion of cancer-associated genes, we employed single-nucleus multiomic sequencing (snRNA+ATAC-seq) to simultaneously interrogate gene expression and chromatin accessibility at single-cell resolution (**Fig. 2A**). Given the well-documented challenges in capturing viable epithelial cells from HNSCC^19^, we optimized tissue handling, dissociation conditions, and nuclear isolation to achieve high epithelial recovery (~40% of total cells captured compared to <10% in previous studies^20–28^), yielding over 60,000 high-quality epithelial nuclei across our dataset (see **Methods, Suppl. Fig. 4A**).

**Figure 2.**
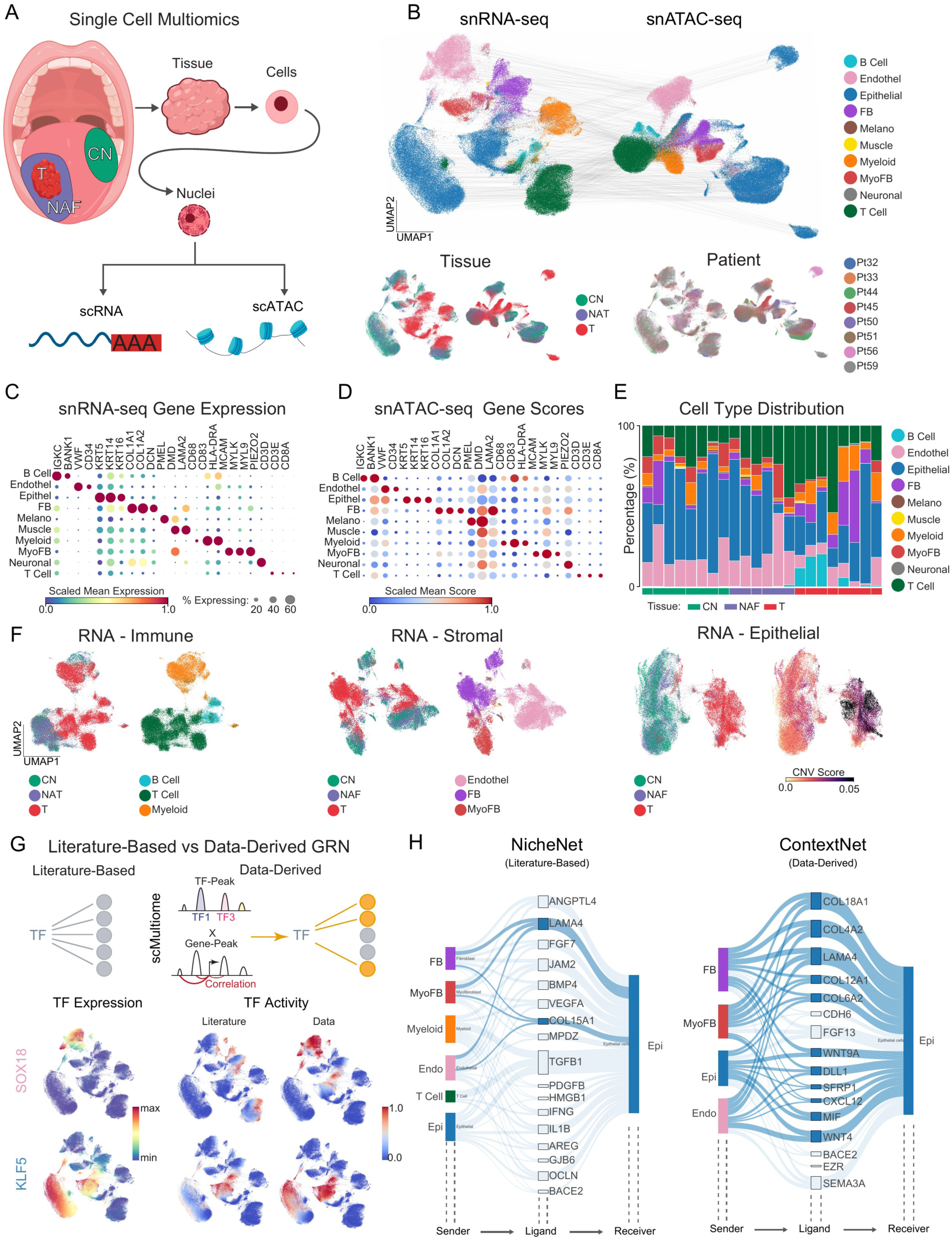
Single-cell multiomics atlas of HNSCC field cancerization. (**A**) Schematic of single-nucleus multiomics workflow. Matched Tumor (T), Normal-Adjacent Field (NAF), and Contralateral Normal (CN) tissues were dissociated into cells. Next, nuclei were isolated for paired snRNA-seq and snATAC-seq. (**B**) UMAP visualization of snRNA-seq and snATAC-seq data colored by cell type, tissue type, and patient. (**C**) Dot plot showing gene expression of canonical marker genes across cell types in snRNA-seq data. (**D**) Dot plot showing gene accessibility scores for marker genes across cell types in snATAC-seq data. (**E**) Cell type composition per sample ordered by tissue type. (**F**) RNA UMAP visualization of immune, stromal, and epithelial compartments colored by tissue and cell subtype, with epithelial cells colored by CNV score computed using inferCNV to distinguish tumor from normal cells, since tumor and normal epithelium can co-occur within the same specimen. (**G**) Top: Schematic showing the identification of data-derived transcription factor targets. Bottom: Comparison of transcription factor activity inferred from the data-derived versus literature-based gene regulatory networks for *SOX18* (endothelial) and *KLF5* (epithelial). Left: gene expression; Middle: literature-based activity; Right: data-derived activity. Activity is computed as mean z-scored expression of the top 150 predicted targets. (**H**) Comparison of NicheNet (left) and ContextNet (right) results, which use literature-based and data-derived regulatory networks respectively, with epithelial cells as receivers and stromal and epithelial cells as senders. For each method, the 15 highest-ranked predicted interactions are shown together with other known interactions identified among the top 30 predictions. Known interactions are highlighted in dark blue.

We profiled 22 fresh tissue samples from 8 treatment-naïve HPV-negative HNSCC patients, capturing 145,769 high-quality nuclei (**Fig. 2B**): 67,520 from tumor, 39,423 from NAF, and 38,826 from CN tissues. Cell type annotation identified epithelial, immune, and stromal populations (**Fig. 2B-F, Suppl. Table S6, 7**). Cell types were generally well-represented across tissue sites and patients (**Fig. 2B, E, Suppl. Fig. 4B**), and tissues contained a similar cell type distribution across samples with only a subset of tumor samples showing elevated B cell infiltration (**Fig. 2E, F, left**), as previously reported in HNSCC^28^. Consistent with our WGS findings, inferCNV^29,30^ analysis confirmed that NAF and CN epithelial cells maintained diploid genomes, while tumor cells harbored elevated copy number alterations (**Fig. 2F, right**). This allowed us to accurately distinguish tumor from non-tumor epithelial cells, even in samples where both were present in close proximity (**Suppl. Fig. 4C, D**).

Characterizing field cancerization also requires understanding how cells communicate within the tissue microenvironment. NicheNet^31^ has emerged as an effective framework for inferring cell-cell communication from single-cell data, integrating ligand-receptor interactions with intracellular signaling and gene regulatory networks that propagate signals from ligands to downstream target genes. However, its reliance on literature-derived regulatory relationships can limit its ability to capture cell-type or tissue-specific gene regulation. Our paired RNA and ATAC measurements presented an opportunity to address this. We constructed a data-derived gene regulatory network linking transcription factors to their target genes based on coordinated changes in accessibility and expression (**Fig. 2G, top**). Our analysis confirmed that this approach generated cell-type-specific transcription factor activity patterns, such as *SOX18* enrichment in endothelial cells and *KLF5* in epithelial cells, whereas the literature-based network yielded diffuse, non-specific patterns (**Fig. 2G, bottom, Suppl. Fig. 5A**). We incorporated this data-derived network into the NicheNet framework to develop ContextNet, a computational tool for data-driven inference of cell-cell communication (**Suppl. Fig. 5B, C**). ContextNet demonstrated greater sensitivity in recovering known activation-associated signaling in independent benchmarks (**Suppl. Fig. 6;** see **Methods**) and outperformed NicheNet in identification of the known basement membrane- and Wnt-mediated signaling interactions between epithelial and stromal cells (**Fig. 2H**).

Together, this single-cell multimodal dataset and computational framework provide the resolution needed to characterize both cell-autonomous and microenvironmental features of field cancerization in HNSCC.

### Single-cell multiomic profiling defines normal oral epithelial hierarchy

We next characterized the molecular landscape and tissue dynamics of human oral epithelium using contralateral normal (CN) samples to establish a baseline of epithelial organization against which to compare normal-adjacent field (NAF) and tumor tissues. A defining feature of oral epithelium is its stratified squamous morphology, where proliferative potential is restricted to stem/progenitor cell compartments proximal to the basement membrane, while postmitotic suprabasal cells provide barrier function and are regularly shed. Two competing models have been proposed to explain how stratified epithelia maintain homeostasis. The classical hierarchical model (**Fig. 3A, left**) posits that slow-cycling stem cells in the basal layer give rise to rapidly dividing transit-amplifying cells that undergo several rounds of division before terminal differentiation^32,33^. Label-retaining cell assays and molecular profiling of murine tissues supported this model by identifying slow-cycling cells distinguishable from their more proliferative progeny^34,35^. More recently, lineage tracing studies challenged this view by demonstrating that tissue homeostasis can be maintained by progenitor populations making stochastic cell fate decisions (**Fig. 3A, right**), with balanced probabilities of basal cell self-renewal and differentiation^36–38^. Studies of the human oral mucosa suggest that the basal layer of oral epithelium is relatively quiescent, with the majority of Ki67-positive proliferating cells located in the parabasal layers^39–42^. This pattern of a quiescent basal compartment overlaid by proliferative parabasal cells is also observed in human esophagus, ectocervix, and vagina, suggesting a conserved hierarchical organizational principle in certain human squamous epithelia^41,43^. However, comprehensive molecular characterization of this proposed hierarchy in human oral mucosa is yet to be done. To better understand potential molecular alterations occurring in NAF tissues, we first sought to define the organization of normal oral epithelial differentiation using our single-cell multiomics data.

**Figure 3.**
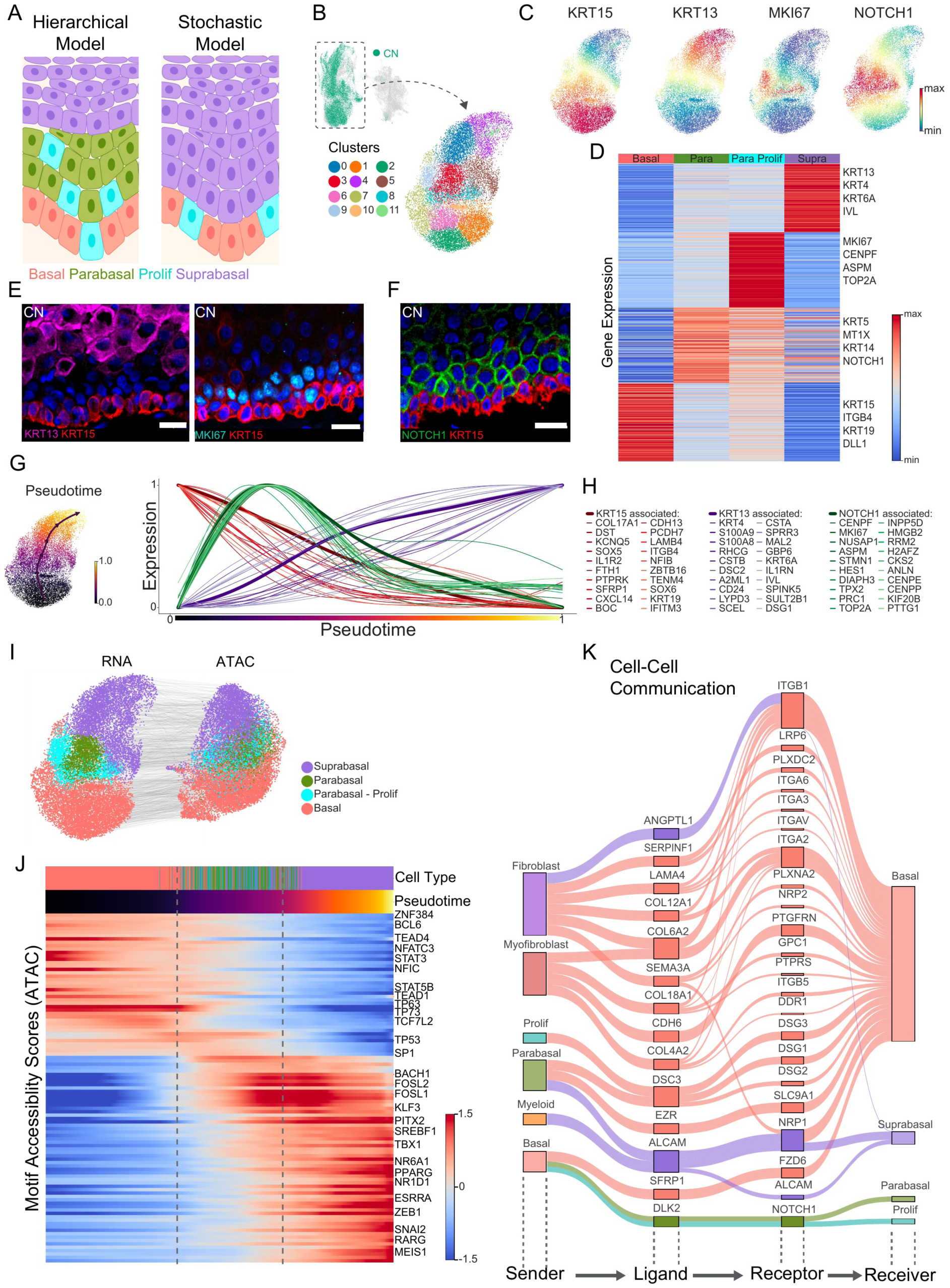
Hierarchical organization of normal human oral epithelium. (**A**) Schematic illustrating the hierarchical (left) and stochastic (right) models of epithelial stem cell differentiation. (**B**) UMAP visualization of contralateral normal epithelial cells colored by Leiden clusters. (**C**) UMAP visualization of marker gene expression: *KRT15* (basal), *KRT13* (suprabasal), *MKI67* (proliferating), and *NOTCH1* (parabasal). (**D**) Heatmap of differentially expressed genes across epithelial cell states. (**E**) Immunofluorescence staining of contralateral normal tissue: KRT13 (magenta)/KRT15 (red) or MKI67 (teal)/KRT15 (red) co-staining. Nuclei are stained with DAPI (blue). Scale bars indicate 20 µm. (**F**) NOTCH1 (green)/KRT15 (red) co-staining in contralateral normal tissue. Nuclei are stained with DAPI (blue). Scale bar indicates 20 µm. (**G**) Pseudotime analysis. Left: UMAP colored by pseudotime. Right: gene expression dynamics across pseudotime for *KRT15*-associated (red), *KRT13*-associated (purple), and *NOTCH1*-associated (green) gene programs. (**H**) Genes positively correlated with *KRT15* (red), *KRT13* (purple), and *NOTCH1* (green) along pseudotime. (**I**) UMAP visualization of epithelial cells from RNA (left) and ATAC (right) modalities colored by cell state. (**J**) ChromVAR transcription factor motif accessibility scores across pseudotime. (**K**) Cell-cell communication analysis using ContextNet. Sankey diagram generated from ccflowR depicting ligand-receptor signaling flow from sender cell types (left) through ligands and receptors to receiver epithelial compartments (right). Edge width reflects a composite score integrating ligand expression in sender cells and predicted regulatory activity on receiver cells; edge color indicates receiver cell type. Top 22.5% of the predicted interactions from **Suppl.** Fig. 9A are shown.

To define the organization of normal oral epithelial differentiation, we analyzed the subset of epithelial cells from contralateral normal tissues (**Fig. 3B, Suppl. Fig. 7A**). KRT15 and KRT13 are established markers of basal and suprabasal cells in human oral epithelium, respectively ^42,44^, and their expression patterns clearly delineated these two populations in our data (**Fig. 3C, D**). Interestingly, *KRT15*-expressing basal cells did not transition directly to *KRT13*-expressing suprabasal cells and instead, we observed an intermediate *MKI67*-high population between these two states (**Fig. 3C, Suppl. Fig. 7B**). The presence of this intermediate population raised the question of whether it represents a stable cell state or a transient phase through which cells rapidly pass. To address this, we applied Mellon^45^, a computational method that estimates cell density across the phenotypic space and identifies cells that transition quickly through a state as occupying low-density regions. The *MKI67*-high population exhibited low cell density, consistent with a transient, rapidly dividing population rather than a stable compartment (**Suppl. Fig. 7C**). To confirm the spatial organization of these cell states, we performed immunofluorescent staining on contralateral normal tissue sections. KRT15 staining was restricted to the basal layer adjacent to the basement membrane, KRT13 marked the suprabasal compartment, and MKI67-positive cells localized almost exclusively to the intermediate layers (**Fig. 3E**).

Prior studies identified proliferation markers such as MKI67 as the defining feature of parabasal cells, leaving it unclear whether non-proliferating cells also reside in this compartment or how they might be identified^41^. To address this question, we performed differential gene expression analysis on our epithelial population. This revealed a broader population of cells between basal and suprabasal cells that is transcriptionally distinct, defined by a set of parabasal cell state markers, including *NOTCH1*, *KRT5*, and *MT1X* (**Fig. 3D, Suppl. Fig. 7D, Suppl. Table S8**) and a subset of cells which are actively proliferating. Among these markers, *NOTCH1* was particularly notable given its established role in promoting the transition from proliferation to differentiation in stratified epithelia^46^. Immunofluorescent staining confirmed NOTCH1 protein expression specifically in the parabasal layers, positioned between KRT15-positive basal cells and KRT13-positive suprabasal cells (**Fig. 3F**). Importantly, NOTCH1 expression was seen throughout the parabasal region, confirming it as a general parabasal marker rather than a proliferation-associated gene and the first specific molecular identifier of the parabasal cell state.

Having established the parabasal compartment as a transcriptional intermediate, we next examined the dynamics of gene expression programs along the continuum of epithelial differentiation. We applied Palantir^47^ trajectory analysis, which orders cells along a pseudotime trajectory based on the transcriptional similarity to a defined starting population, in this case basal cells. We identified genes whose expression changed coordinately with *KRT15*, *KRT13*, and *NOTCH1* along this trajectory (**Fig. 3G, H**), and observed that the gene expression dynamics defining the oral epithelial hierarchy proceeded through continuous rather than stepwise transitions. Basal stem cell genes, including *KRT15*, *COL17A1* and *ITGB4*, were highest at the start of the trajectory and were downregulated progressively with differentiation. Differentiation genes including *KRT13*, *DSC2*, and *IVL* were upregulated in the terminal suprabasal state. Proliferation genes such as *MKI67* and *TOP2A* peaked in the parabasal compartment, consistent with the transit-amplifying nature of these cells. These coordinated transcriptional dynamics confirm that proliferation is spatially and temporally enriched in the parabasal compartment, forming a continuous bridge between the relatively quiescent basal stem cell pool and postmitotic differentiated cells.

Next, we sought to understand the regulatory landscape and microenvironmental signals controlling epithelial cell differentiation in the oral mucosa by leveraging the paired RNA-ATAC nature of our data. Cells clustered similarly in both RNA and ATAC profiles, confirming that chromatin state broadly tracks with transcriptional identity along the differentiation trajectory (**Fig. 3I**). To identify key transcription factors (TF) driving differentiation, we used chromVAR^48^ to quantify TF motif accessibility across cells and examined how accessibility of these motifs changed over pseudotime (**Fig. 3J, Suppl. Fig. 8**). To connect these intrinsic programs to extracellular signals, we applied ContextNet, a computational framework we developed that uses data-derived gene regulatory networks to link ligand-receptor interactions to downstream transcriptional programs (**Fig. 3K, Suppl. Fig. 9A**).

In the basal compartment, peaks with motifs for several known epithelial TF programs were highly accessible (**Fig. 3J)**. These included TEAD1/4, the DNA-binding partners of YAP/TAZ in the Hippo signaling pathway, which are known for their basal cell-specific transcriptional activity in promoting stem cell self-renewal in stratified squamous epithelia^49–52^. These TFs were predicted to regulate the expression of genes in the *KRT15*-associated program (**Fig. 3H**) including *KRT15*, *CDH13*, and *TENM4* (**Suppl. Fig. 8**). In addition, TP63 and TP73, known master regulators of stratified epithelial identity and stem cell renewal^53–55^, exhibited high motif accessibility in basal cells (**Fig. 3J**) and were predicted to regulate the expression of *KRT15*-associated genes *DST*, *CXCL14*, and *CDH13* (**Suppl. Fig. 8**). Using ContextNet, we identified basal cells as the primary receivers of stromal signals: fibroblasts and myofibroblasts provided extracellular matrix cues (LAMA4, COL18A1, SERPINF1), with these signals converging on integrin receptors (ITGB1, ITGA2) and LRP6, consistent with maintenance of basement membrane attachment and stem cell niche signaling (**Fig. 3K**). Ligand-target analysis predicted that these diverse stromal inputs also converge on a shared set of *KRT15*-associated genes (**Fig. 3H**), with *ZBTB16*, *FTH1*, and *TP73* emerging as among the strongest predicted downstream targets across multiple ligands (**Suppl. Fig. 9B**). The same genes were also among the targets of basal TFs identified by chromVAR, indicating that extrinsic stromal cues feed into transcription factor programs that intrinsically define basal identity.

At the opposite end of the trajectory, the suprabasal compartment was defined by coordinated activation of several well-characterized differentiation programs. The first program was an AP-1 module driven by FOSL1 and FOSL2 (**Fig. 3J**) with regulatory links to cornified envelope and barrier genes^56–58^ including *IVL*, *SPRR3* and *SCEL* (**Suppl. Fig. 8**). The second was lipid-sensing transcription factors SREBF1 and SREBF2 with predicted targets in lipid metabolism and barrier formation such as *SULT2B1*, as well as broad regulation of *KRT13*-associated differentiation genes (**Fig. 3H, Suppl. Fig. 8**)^56–58^. Together, these programs coordinate the structural and metabolic requirements of terminal differentiation and barrier formation. As suprabasal cells differentiate and delaminate from the basement membrane, they lose direct contact with the underlying stroma, and their signaling environment shifts accordingly. ALCAM-NRP1 signaling emerged as the strongest ligand-receptor interaction in the suprabasal compartment. This signaling axis, previously implicated in barrier function, showed the strongest predicted regulatory potential for a broad subset of the *KRT13*-associated gene program, including *IL1RN*, *IVL*, *RHCG*, *SPRR3*, *CSTB*, *FOSL1* (**Fig. 3K, Suppl. Fig. 9B**)^59^.

While the transcription factor programs governing basal and suprabasal identity are well characterized, the regulatory landscape of the parabasal compartment is not well defined. Unlike the basal and suprabasal populations which were defined by distinct TF regulatory programs, parabasal cells exhibited reduced basal TF motif accessibility alongside the emergence of accessible chromatin at *E2F6*, *YBX1*, and *NFYB* loci (**Suppl. Fig. 8**). *NOTCH1* expression itself peaked in the parabasal compartment (**Fig. 3D, F**), and *HES1*, a canonical Notch target, was among the *NOTCH1*-correlated genes identified by trajectory analysis (**Fig. 3G, H**). Consistent with the known roles of TP53 and TP63 in *NOTCH1* transcriptional regulation^60,61^, both showed predicted regulatory links to *NOTCH1* in this compartment (**Suppl. Fig. 8**), suggesting that the parabasal state represents the site where p53/p63-dependent Notch activation may initiate the transition from proliferation to differentiation commitment. Consistent with this, the primary microenvironmental inputs to the parabasal and proliferating compartments were the DLK2 signal from basal cells, predicted to signal through NOTCH1 with *HES1* as the top downstream target (**Fig. 3K, Suppl. Fig. 9B**).

Collectively, these analyses provide a comprehensive molecular map of human oral epithelial differentiation, defining the transcriptional programs, chromatin dynamics, and intercellular signaling that arrange the hierarchical progression from largely quiescent basal stem cells through the proliferative parabasal compartment to terminally differentiated suprabasal cells.

### Epithelial hierarchy is disrupted in field cancerization

Having defined the molecular architecture of normal oral epithelial differentiation in CN tissues (**Fig. 3**), we next asked whether this hierarchy is preserved in the normal-adjacent field (NAF). Disruption of epithelial organization is a hallmark of squamous cell carcinoma, and the degree of morphological disarray is a primary criterion used by pathologists to grade disease progression^62^. Our genomic analyses demonstrated that NAF harbors expanded mutant clones independent of those driving the adjacent tumors (**Fig. 1G, H**), raising the question of whether non-genetic changes in tissue organization may contribute to clonal expansion in the field-cancerized epithelium.

To characterize the epithelial hierarchy in NAF, we combined epithelial cells from NAF and CN tissues in our data and transferred the cell-type assignments derived from CN samples (**Fig. 4A, B**). NAF and CN cells largely overlapped in cell-state space across patient samples, pseudotime, and cell type (**Fig. 4A-C, Suppl. Fig. 10A**), indicating that the overall epithelial hierarchy was broadly maintained. However, when we examined the tightly coordinated gene expression programs identified in CN, specifically the *KRT15*, *KRT13*, and *NOTCH1*-associated gene modules that define the basal-to-suprabasal differentiation trajectory (**Fig. 3G, H**), NAF cells displayed greater transcriptional noise, indicating a potential disruption of the epithelial hierarchy observed in contralateral normal tissue (**Fig. 4D**).

**Figure 4.**
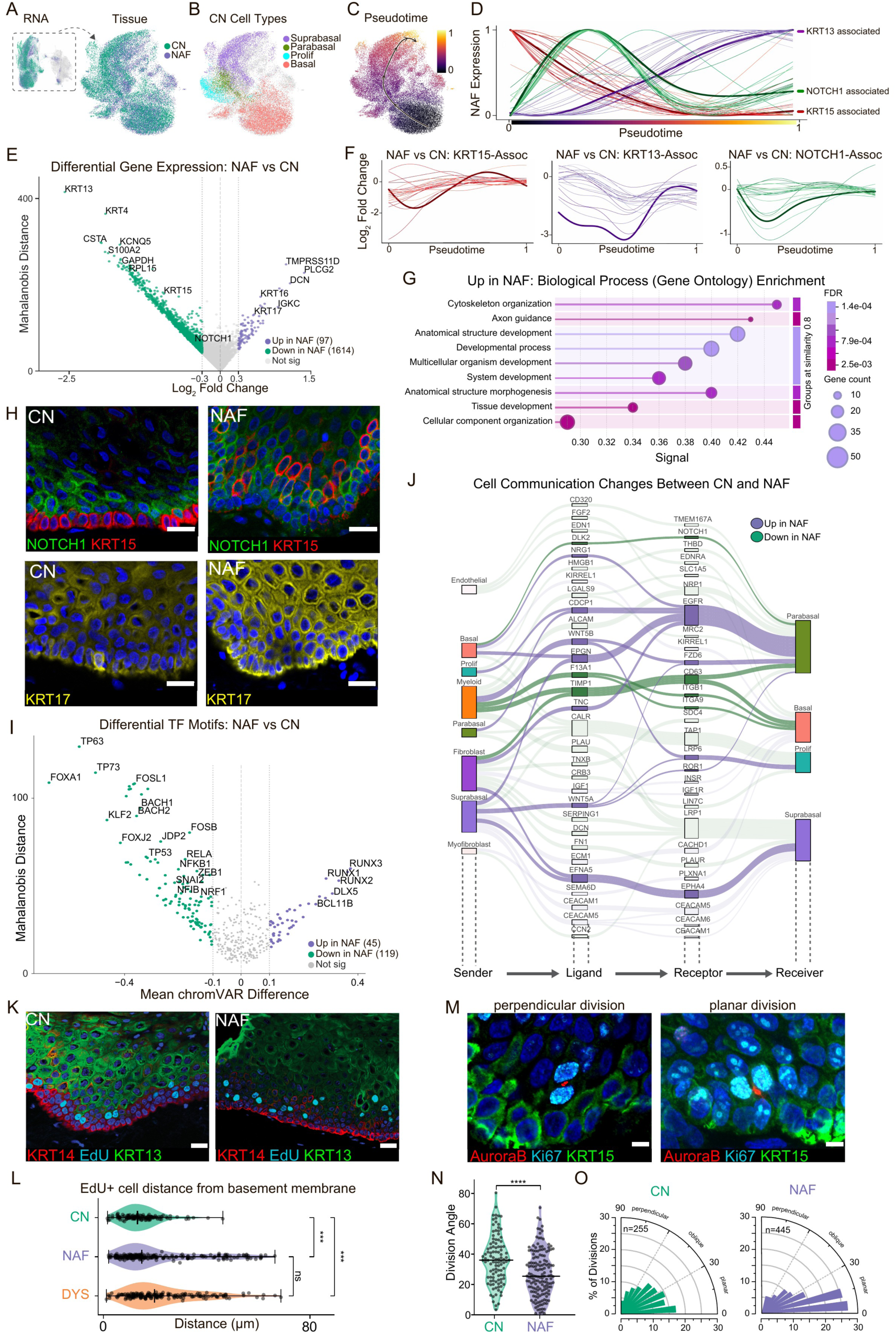
Epithelial hierarchy is disrupted in field cancerization. RNA UMAP visualizations of co-embedded CN (contralateral normal) and NAF (normal-adjacent field) epithelial cells colored by (**A**) tissue type, (**B**) CN cell type including Suprabasal, Parabasal, Proliferative (Prolif), Basal, and (**C**) pseudotime calculated using basal as a starting cell state. (**D**) Gene expression dynamics for NAF epithelial cells for *KRT15*-associated (red), *NOTCH1*-associated (green), and *KRT13*-associated (purple) gene programs over pseudotime (for gene list see Fig. 3H). (**E**) Kompot differential expression analysis comparing NAF to CN epithelial cells. Volcano plot shows log_2_ fold change versus Mahalanobis distance; genes significantly upregulated in NAF (purple, n = 97) and downregulated in NAF (green, n = 1,614) are highlighted. (**F**) Log_2_ fold change of individual genes in NAF relative to CN across pseudotime for *KRT15*-associated (left - red), *KRT13*-associated (middle - purple), and *NOTCH1*-associated (right - green) gene modules. Each line represents the log_2_ fold change of a single gene from Fig. 3H in NAF relative to CN across pseudotime, where a value of zero indicates no difference between tissues, negative values indicate lower expression in NAF, and positive values indicate higher expression in NAF. (**G**) STRINGdb^72^ Gene Ontology (Biological Process) enrichment analysis of genes significantly upregulated in NAF. Dot size indicates gene count; color indicates FDR. (**H**) Immunofluorescence staining of CN and NAF tissues for NOTCH1 (green) and KRT15 (red) (top panels) and KRT17 (yellow, bottom panels). Nuclei are stained with DAPI (blue). Scale bars indicate 20 µm. (**I**) Kompot differential analysis of ATAC chromVAR transcription factor motif accessibility scores comparing NAF to CN. Scatter plot shows mean chromVAR score difference versus Mahalanobis distance; TFs with increased accessibility in NAF (purple, n = 45) and decreased accessibility in NAF (green, n = 119) are highlighted. (**J**) ContextNet cell-cell communication analysis showing differential ligand-receptor signaling between NAF and CN tissues. Sankey diagram depicts signaling flow from sender cell types (left) through prioritized ligands and receptors to receiver epithelial compartments (right). Interactions upregulated in NAF are shown in purple; interactions downregulated in NAF are shown in green. The top ranked routes are shown. (**K**) Immunofluorescence staining of CN (top) and NAF (bottom) tissue slice cultures for KRT14 (red), EdU (cyan), KRT13 (green), and DAPI (blue). Scale bars indicate 20 µm. (**L**) Distance of EdU^+^ nuclei from the basement membrane (defined by basal KRT14 signal) in contralateral normal (CN, n = 124), normal-adjacent field (NAF, n = 225), and dysplastic (DYS, n = 177) tissues. Violin plots show data distribution with median (horizontal line); individual data points are overlaid. Statistical significance was determined by Welch’s ANOVA followed by Games-Howell post hoc tests. ***p < 1e-3, ns = not significant. (**M**) Representative images of perpendicular and planar cell divisions using immunofluorescence staining of Aurora B kinase (red) to measure mitotic spindle orientation and division angle, co-stained with Ki67 (cyan), KRT15 (green) and DAPI (blue). (**N**) Division angles for individual cells in CN and NAF patient tissues (n= 3 patients per tissue type). Cell divisions quantified per tissue: n= 255 (CN), n= 445 NAF. Mann-Whitney U = 68928, p = 2.2×10^−6^. Scale bars indicate 5 µm. (**O**) Cell division angle rose plots for CN and NAF with 10° bins.

To test this, we applied Kompot^63^ differential expression analysis to identify expression changes between NAF and CN epithelial cells. By sharing information across neighboring states and accounting for their statistical dependencies, Kompot detects subtle transcriptional changes even when the underlying cell populations are highly similar. Kompot revealed widespread downregulation of key epithelial differentiation genes in NAF relative to CN, including *KRT15, KRT13*, and *NOTCH1*, the canonical markers of oral epithelial differentiation (**Fig. 4E, Suppl. Table S9**). To next determine where along the differentiation continuum these changes occurred, we examined the expression changes in *KRT15*, *KRT13*, and *NOTCH1*-associated gene modules (**Fig. 3H**) across pseudotime (**Fig. 4F, Suppl. Fig. 10B-D**). *KRT15*-associated basal genes were downregulated early in differentiation but paradoxically upregulated later along the epithelial trajectory, suggesting aberrant retention of basal gene expression in differentiating NAF cells. *KRT13*-associated suprabasal genes were consistently downregulated across the full differentiation axis, indicating a broad reduction of the suprabasal differentiation program in NAF. *NOTCH1*-associated genes were reduced during the proliferative window but modestly increased at terminal differentiation. Together, these patterns suggest that the coordinated transition from basal to suprabasal identity is disrupted in NAF, with basal programs persisting inappropriately and suprabasal programs failing to fully activate.

Additionally, we observed that the gene expression changes between NAF and CN were asymmetric, with the number of downregulated genes (>1600) substantially greater than the upregulated genes (<100) (**Fig. 4E**). Gene ontology enrichment analysis of genes downregulated in NAF revealed suppression of core cellular machinery, with the most enriched terms being ribosomal biogenesis, cytoplasmic translation, and ubiquitin-protein regulation (**Suppl. Fig. 10E**), suggesting a broad reduction in translational capacity in the field-cancerized epithelium. This was coupled with upregulation of cancer-associated genes in NAF, including *KRT17, TMPRSS11D*, *PLCG2*, and *DCN*, which have been implicated in cellular stress and tumor growth and invasion^64,65^. Gene ontology enrichment of the upregulated genes highlighted developmental processes, cytoskeleton organization, and anatomical structure morphogenesis (**Fig. 4G**), supporting our findings that tissue morphogenesis-related basal gene programs are maintained and the expected transitions in normal differentiation appear disrupted.

We validated that these transcriptional changes were reflected in altered protein expression using immunofluorescence staining (**Fig. 4H**, **Fig. 3F**). In CN tissues, KRT15 protein was restricted to the basal layer and NOTCH1 was expressed in parabasal cells, consistent with the normal differentiation hierarchy. However, in NAF tissues, KRT15 expression, while overall at a lower level, was expanded beyond the basal compartment into suprabasal cells, while NOTCH1 expression was generally diminished (**Fig. 4H**). Supporting our differential gene expression results, KRT17 was markedly more abundant in NAF compared to CN (**Fig. 4H**). Together, these findings across transcriptomic and protein-level analyses confirm that field cancerization significantly affects the programs governing epithelial differentiation and tissue organization.

Having identified widespread transcriptional disruption of the epithelial hierarchy in NAF, we next leveraged the paired RNA-ATAC nature of our data to determine whether these changes are reflected in the underlying chromatin regulatory landscape. We applied Kompot to compare chromVAR TF motif accessibility scores between NAF and CN epithelial cells across the shared ATAC cell-state landscape (**Fig. 4I, Suppl. Table S10**). Strikingly, the loss of chromatin accessibility at TF loci in NAF corresponded closely to the regulatory programs that define each compartment of the normal epithelial hierarchy described in **Fig. 3** (**Fig. 4I, Suppl. Fig. 10F**). Motif accessibility for basal master regulators TP63 and TP73 showed the largest and most significant reductions, alongside FOXA1. Motif accessibility for AP-1 family members (FOS and JUN factors) that drive the suprabasal differentiation program was consistently reduced, as was accessibility for the regulators of lipid metabolism and barrier formation in the suprabasal compartment. The parabasal-associated factors YBX1, NFYB, and TP53, which we identified as marking the transition from proliferation to differentiation commitment, were also significantly reduced. These results indicate that field cancerization does not selectively disrupt a single layer of the epithelium but rather alters the regulatory programs governing the hierarchical differentiation trajectory.

In contrast, in NAF, there was increased accessibility at loci for the RUNX family members, previously implicated in epithelial-to-mesenchymal transition and invasion in head and neck squamous cell carcinoma^66–68^ (**Fig. 4I**). Further TF loci with increased accessibility included *DLX5*, *BCL11B*, and at WNT pathway effectors *LEF1* and *TCF7L2* **(Fig. 4I, Suppl. Table S10**). While some of these factors play roles in normal epithelial regulation, their coordinated increase in chromatin accessibility in NAF, alongside the RUNX family, suggests the activation of aberrant regulatory programs associated with developmental and oncogenic pathways. We also observed increased accessibility at motifs for repressors such as REST and PRDM1 in NAF cells. Since these factors normally maintain a repressed chromatin state at their target sites, increased openness at these motifs suggests a loss of repressor occupancy, and consequently, a weakening of their role in maintaining cell identity (**Suppl. Table S10**).

To determine whether the transcriptional and chromatin disruption of the NAF epithelial hierarchy is accompanied by altered microenvironment signals, we applied ContextNet using the MultiNicheNet^69^ framework to compare predicted ligand-receptor communication and downstream gene expression changes between NAF and CN tissues (**Fig. 4J, Suppl. Fig. 11**). Two patterns emerged. First, signaling axes that maintain the normal hierarchy were reduced in NAF. In the basal compartment, TIMP1 signaling from fibroblasts was reduced, with predicted loss of *KRT15*-associated targets including *KRT19*, consistent with erosion of normal basal identity (**Fig. 4J, Suppl. Fig. 11A**). In the parabasal compartment, the DLK2-NOTCH1 axis from basal cells was downregulated, consistent with the reduced NOTCH1 expression and protein staining (**Fig. 4J, Suppl. Fig. 11A**). Second, establishment of EFNA5-EPHA4 signaling, part of the Ephrin pathway known for its developmental roles in establishing compartment boundaries^70^, from fibroblasts and suprabasal cells was directed toward the suprabasal compartment, with KRT17 among predicted downstream targets (**Fig. 4J, Suppl. Fig. 11B**). Suprabasal cells themselves became more prominent senders to other epithelial compartments through WNT signaling (WNT5A, WNT5B, LRP6, FZD6), suggesting that the altered cell identities generate novel communication circuits (**Fig. 4J, Suppl. Fig. 11B**). Similarly, increased EGFR-directed signaling to parabasal cells converged from multiple cell types, including TNC from fibroblasts, CDCP1 from proliferating epithelial cells, EPGN from suprabasal cells (**Fig. 4J, Suppl. Fig. 11B**). Together, these results indicate that disruption of the epithelial hierarchy in NAF is not only reflected in cell-intrinsic regulatory changes but is reinforced by a reorganized signaling environment, where the loss of homeostatic cues and the emergence of aberrant communication circuits may jointly sustain the disorganized tissue state.

The convergence of EGFR-directed signaling on parabasal cells in NAF raised the question of whether these altered signaling patterns translate to functional changes in proliferative behavior. To test this, we performed EdU pulse-labeling experiments on patient tissue slice cultures to mark cells actively entering S-phase (see **Methods**). Unlike Ki67, which marks cycling cells, EdU incorporation during a defined 2-hour pulse specifically labels cells actively synthesizing DNA, providing a more precise readout of proliferative activity and its spatial distribution within the tissue. Using this approach, we found that, in CN tissues, EdU incorporation was restricted primarily to the parabasal layer, with some basal proliferation, consistent with the normal proliferative architecture of oral epithelium (**Fig. 3E**, **Fig. 4K, left**). In contrast, NAF tissues contained proliferating cells displaced into the suprabasal compartment, a region normally restricted to terminally differentiated, post-mitotic cells (**Fig. 4K, right**). Quantification of EdU+ cell distance from the basement membrane confirmed that the proliferative zone in NAF is significantly expanded relative to CN (p < 1e-3, Welch’s ANOVA with Games-Howell post hoc test), with a distribution not significantly different from overtly dysplastic tissues (**Fig. 4L**). We also assessed the orientation of cell division by measuring mitotic spindle angles between newly generated daughter cells^71^. Compared to CN, dividing cells in NAF tissues exhibited an increase in planar cell divisions, indicative of higher rates of symmetric cell renewal (**Fig. 4M-O**). These results demonstrate that non-genetic alterations in tissue organization and cell behavior, including loss of the spatial constraints on proliferation that normally govern tissue homeostasis, are already present in histologically normal field-cancerized epithelium.

### Identification and characterization of an emergent NAF-specific cell population

Given the widespread disruption of epithelial transcriptional programs and chromatin landscapes in NAF (**Fig. 4**), we next asked whether these alterations lead to changes in composition of epithelial cell types or generate distinct cell populations not present in contralateral normal tissue. Kompot^63^ differential abundance analysis identified an epithelial population significantly enriched in NAF relative to CN cells (**Fig. 5A, B, Suppl. Fig. 12A, B**). These cells formed a transcriptionally distinct cluster that did not correspond to any of the previously characterized normal epithelial subtypes (basal, parabasal, proliferating, or suprabasal), suggesting the appearance of a novel ‘emergent state’ within the cancerized field (**Fig. 5C**). This population was consistently detected across multiple patients (**Suppl. Fig. 12C, D**), indicating it represents a recurrent feature of field cancerization rather than patient-specific variation.

**Figure 5.**
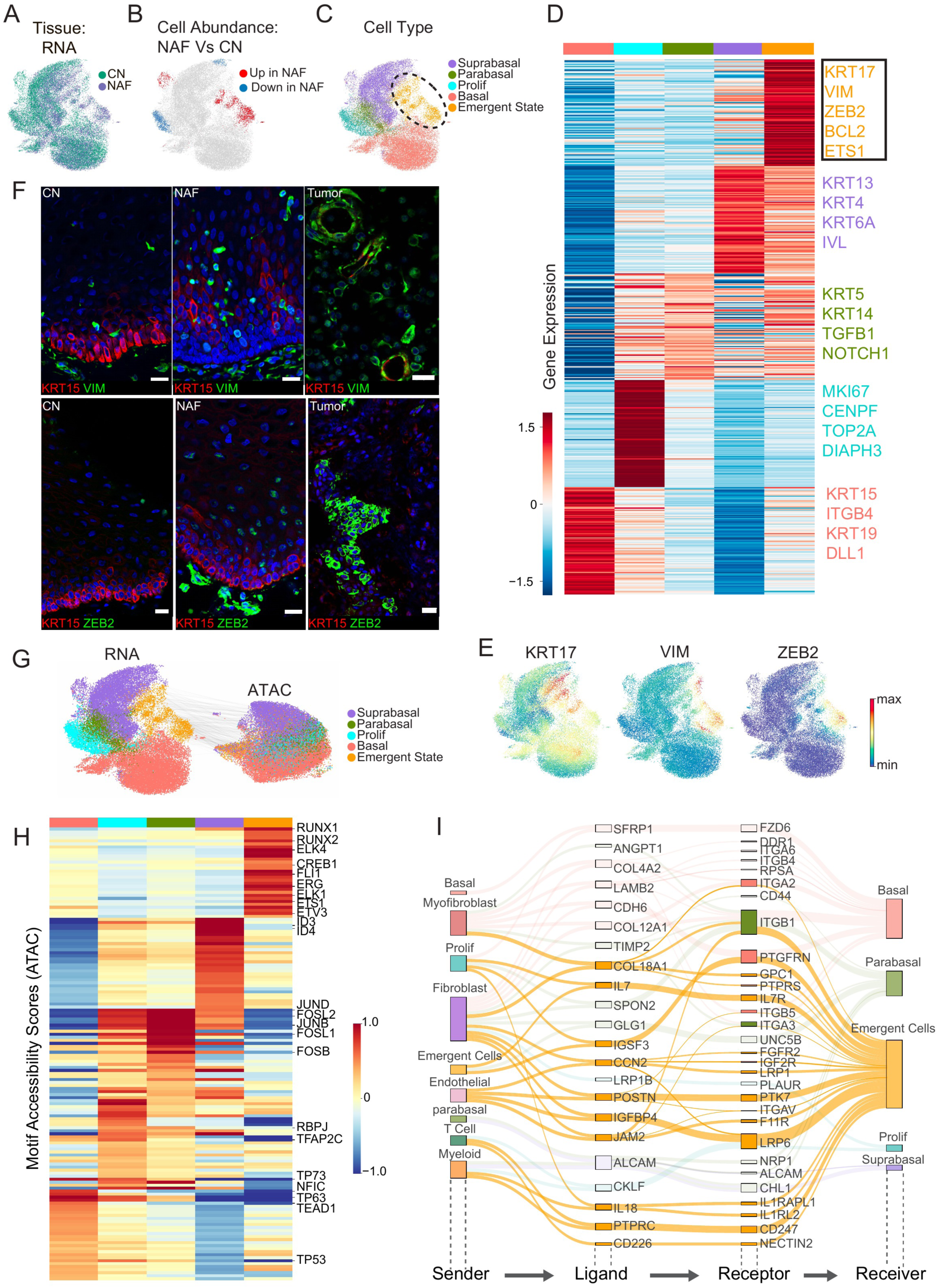
Identification of an emergent NAF-specific epithelial population with tumor features. (**A**) RNA UMAP of co-embedded CN (contralateral normal) and NAF (normal-adjacent field) epithelial cells colored by tissue type. (**B**) UMAP colored by Kompot differential abundance analysis comparing NAF to CN epithelial cells; red indicates enrichment in NAF, blue depletion in NAF. (**C**) UMAP colored by cell type annotation, including the Emergent State population (dashed circle). (**D**) Heatmap of differentially expressed genes across epithelial cell states. (**E**) UMAP visualization of *KRT17*, *VIM*, and *ZEB2* expression in CN and NAF epithelial cells. (**F**) Immunofluorescence co-staining of KRT15 (red) with VIM (green, top row) and KRT15 (red) with ZEB2 (green, bottom row) in CN, NAF, and tumor tissues. Nuclei stained with DAPI (blue). Scale bars indicate 20 µm. (**G**) UMAPs of epithelial cells from RNA and ATAC modalities colored by cell type, including the emergent state population. (**H**) Heatmap of chromVAR transcription factor motif accessibility scores across epithelial cell states, with selected gained (top) and lost (bottom) motifs labeled. (**I**) ContextNet cell-cell communication analysis showing ligand-receptor signaling flow directed toward emergent state compartments (yellow) with other connections towards epithelial cell types in the background.

To characterize these emergent state cells, we first performed differential gene expression analysis comparing them to all other epithelial cells. This revealed that these cells express a distinct molecular program (**Fig. 5D, Suppl. Table S11**). Among the most prominent markers, these cells exhibited elevated expression of *KRT17*, a stress-responsive keratin^73^; *VIM* (vimentin), a marker of epithelial-mesenchymal plasticity^74^; *ZEB2*, a transcriptional repressor of epithelial identity^75^; *BCL2*, an anti-apoptotic regulator^76^; and *ETS1*, a transcription factor involved in cell fate specification and invasion^77^ (**Fig. 5D, E**).

To visualize these cells *in situ*, we performed immunofluorescence staining for VIM and ZEB2 alongside the basal marker KRT15 across CN, NAF, and tumor tissues (**Fig. 5F**). In CN epithelium, VIM and ZEB2 expression was largely absent, with VIM signal mostly restricted to stromal cells beneath the epithelial compartment. In NAF, VIM-positive and ZEB2-positive epithelial cells were clearly identifiable throughout the epithelium, with enrichment in the parabasal layers. These cells were interspersed among KRT15-positive basal cells but extended beyond the basal compartment, consistent with the transcriptional identity defined by our single-cell analysis. We also compared staining against tumor samples where we saw VIM and ZEB2 widely expressed in the tumor epithelial cells.

Having defined the transcriptional identity of the emergent population, we next sought to characterize the regulatory landscape that distinguishes these cells from normal epithelial states. Paired RNA and ATAC profiles clustered concordantly across all epithelial populations, including the emergent state cells, confirming that the transcriptional changes in this population are accompanied by corresponding shifts in chromatin accessibility (**Fig. 5G**). To identify the transcription factors driving this altered chromatin state, we compared chromVAR motif accessibility scores in emergent state cells against basal, dividing, parabasal, and suprabasal populations (**Fig. 5H**).

Emergent state cells showed a striking loss of chromatin accessibility at transcription factor motifs that define the normal epithelial hierarchy. Importantly, we observed reduced motif accessibility at *TP63* and *TP73*, the master regulators of stratified epithelial identity, followed by *NFIC* and *TFAP2C* (**Fig. 5H, Suppl. Table S12**). Motif accessibility for basal-associated programs including at *TEAD1*/*4* loci were also reduced, as were motifs for AP-1 factors that drive suprabasal differentiation. The coordinated loss of regulatory programs spanning all epithelial compartments indicates that emergent state cells have departed from the normal differentiation hierarchy rather than being arrested at a particular stage within it. In place of these programs, emergent state cells gained accessibility at motifs for transcription factor families implicated in cell fate transitions, invasion, and stress response. The most prominent increase was observed at loci for the ETS family, with *ETS1* also transcriptionally upregulated (**Fig. 5D, H, Suppl. Table S12**). ETS factors regulate matrix metalloproteinase expression, cell migration, and angiogenesis, and their overexpression in stratified epithelia has been shown to block terminal differentiation and promote invasive behavior^78,79^. Motif accessibility of the RUNX family and their obligate binding partner CBFB were also coordinately gained, consistent with the RUNX enrichment we observed more broadly in NAF epithelium (**Fig. 4I**, **Fig. 5H, Suppl. Table S12**). A third module of gained motif accessibility was CREB/ATF family members suggesting activation of stress-responsive and cAMP-dependent signaling in this population^80^.

To determine whether the altered regulatory state of emergent cells is accompanied by changes in microenvironmental communication, we applied ContextNet to infer ligand-receptor interactions directed toward the emergent population (**Fig. 5I, Suppl. Fig. 13A, B**). Emergent state cells retained a subset of the stromal interactions that characterize the normal basal niche, including COL18A1 from fibroblasts and myofibroblasts. However, whereas these ligands were previously linked to the KRT15-associated basal program, their predicted downstream targets in emergent state cells shifted to include KRT17 (**Suppl. Fig. 13C**), suggesting that the basal signaling architecture is being co-opted to support the altered transcriptional state. Additionally, emergent state cells received new stromal inputs not predicted in any normal epithelial compartment, including POSTN from fibroblasts and endothelial cells to PTK7 receptors, a pairing that converges on Wnt/β-catenin signaling and has been independently associated with stemness and invasiveness in epithelial tumors^81,82^. Myeloid cells contributed PTPRC signaling to CD247, resulting in downstream activation of RUNX1 among the predicted downstream effectors (**Suppl. Fig. 13C**).

### NAF emergent cells co-localize with parabasal cells and are characterized by tumor-like features

To next examine how the emergent state relates to malignant cells, we co-embedded all epithelial cells from CN, NAF, and tumor samples (**Fig. 6A**). As expected, tumor cells were transcriptionally distinct from normal epithelial populations, consistent with the somatic copy number alterations that distinguish them (**Fig. 2F, Suppl. Fig. 4C, D**). We computed a tumor similarity score for each cell based on its transcriptional distance from normal contralateral cells (**Fig. 6B**, see **Methods**). Emergent state cells were intermediate between normal epithelial populations and tumor, indicating that they represent a state along the transcriptional continuum toward malignancy rather than an unrelated altered state (**Fig. 6B, C**). Copy number analysis confirmed that emergent state cells lack the somatic alterations present in the adjacent tumors (**Fig. 2F, Suppl. Fig. 4C, D**), establishing that their transcriptional convergence with tumor reflects molecular reprogramming within genomically normal epithelium rather than contamination by malignant cells. Comparison of differentially expressed genes between emergent and tumor cells revealed substantial overlap, with tumor cells expressing many of the same markers that define the emergent state (**Suppl. Fig. 14A**), further supporting the transcriptional continuity between these populations. Many of the genes upregulated in emergent state cells and in tumor cells, including *KRT17* and *VIM*, showed a progressive gradient of expression from normal through emergent state to tumor (**Suppl. Fig. 14B**). Immunofluorescence of tumor tissue showed widespread KRT17 expression alongside general loss of KRT15 and NOTCH1 (**Fig. 5F, Suppl. Fig. 14C**), consistent with full progression along the trajectory initiated in the emergent population.

**Figure 6.**
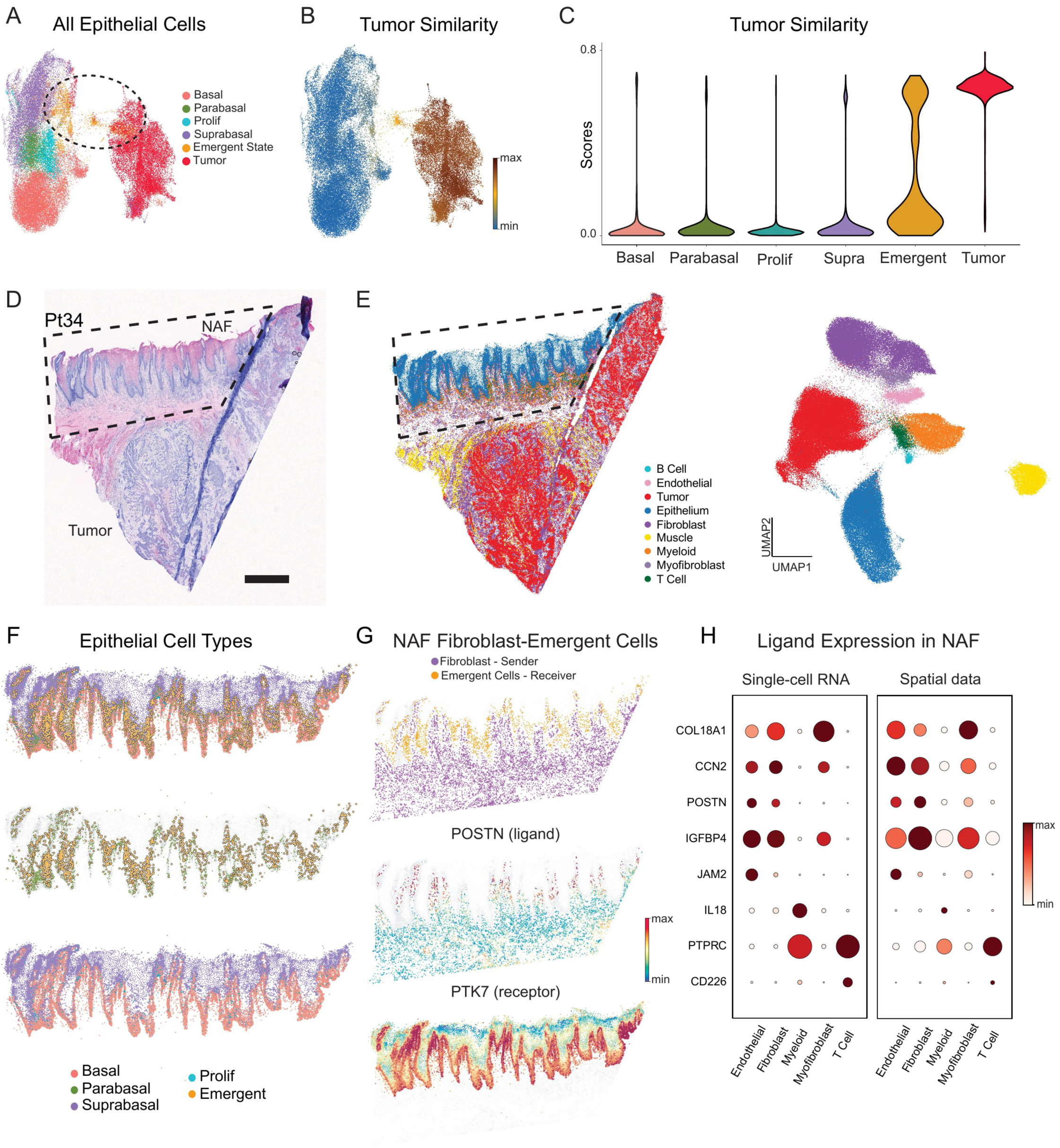
NAF emergent cells are characterized by tumor-like features and are co-localized with parabasal cells. (**A**) UMAP of all epithelial cells from CN, NAF, and tumor samples colored by cell type. Dashed circle highlights emergent state cells. (**B**) UMAP from (**A**) colored by tumor similarity; warmer colors indicate greater transcriptional proximity to tumor. (**C**) Quantification of the tumor similarity for different epithelial cell types. (**D**) H&E staining of a HNSCC tissue section with NAF and tumor regions. NAF region is highlighted in the dotted box. (**E**) Left: Single-cell resolution spatial transcriptomics profiling of the tissue section in (D) colored by cell types. Right: UMAP of the spatial cells colored by cell types. (**F**) NAF region of the (E), showing all epithelial cells (top), parabasal and emergent cells (middle), and basal and suprabasal cells (bottom). (**G**) Top: Epithelial cells and the surrounding stroma in the NAF region highlighting fibroblasts in purple and emergent cells in orange. Middle: Expression of the POSTN ligand in fibroblasts. Bottom: Expression of the PTKR receptor in epithelial cells. (**H**) Dot plots showing the expression of predicted ligands across different sender populations from single-cell RNA data (left) and spatial data (right) in the NAF region. Dot size indicates the percentage of cells expressing the ligand, and color indicates mean scaled expression.

To next determine the spatial organization of emergent state cells within intact tissue architecture we performed single-cell resolution, whole-transcriptome spatial profiling on patient-matched tumor and NAF tissue sections (see **Methods**). We profiled tissues from patients both within (**Suppl. Fig. 15A, C**) and outside (**Fig. 6D, Suppl. Fig. 15B**) of our single-cell cohort. NAF tissue showed reactive atypia without dysplasia in both samples as verified by a pathologist. Spatial transcriptomics data recovered all major cell types identified by snRNA-seq, confirming that our profiling captured the full cellular diversity of the HNSCC microenvironment (**Fig. 6E, Suppl. Fig. 15A-C**). We next separated the NAF and tumor sites from the spatial data based on histology (**Fig. 6D, E**) and transferred cell-type labels from single-cell data to spatially resolve all epithelial cell states (**Suppl. Fig. 16A-D**). Basal, parabasal, and suprabasal cells and transitions between them were identified as expected, with basal cells proximal to the stroma, suprabasal to the outside, and parabasal in between (**Fig. 6F, Suppl. Fig. 16E, F**). Emergent state cells were spatially localized within and throughout the parabasal epithelial compartment of NAF tissue (**Fig. 6F, Suppl. Fig. 16G, H**, see **Methods**). This spatial positioning indicates that the emergent cell state arises at the interface between basal stem cells and the proliferative parabasal region where the normal differentiation hierarchy begins to fail. We similarly analyzed CN tissues from the same patients and observed expected expression of normal epithelial markers and low expression of emergent genes (**Suppl. Fig. 16I, J)**, suggesting that the NAF-specific nature of the emergent state is not an artifact of tissue processing or data analysis.

Emergent cell state identity is in part promoted by aberrant cell-cell communication **(Fig. 5I**). We therefore utilized our spatial data to determine whether the predicted cell interactions are spatially co-localized in intact tissue. Given their potential role in epithelial tumors, we first tested whether the predicted interaction from fibroblasts to emergent cells mediated by POSTN ligand and PTK7 receptors respectively are spatially co-localized (**Fig. 6G**). We observed highest POSTN expression in fibroblasts closest to the epithelium, and high PTK7 expression in basal and parabasal layers, fulfilling the spatial requirement for the predicted interaction. Further, the expression patterns of ligands mediating other predicted interactions in emergent cells were also consistent between single-cell and spatial data (**Fig. 6H, Suppl. Fig. 16K**). POSTN-PTK7 signaling persisted in tumor, with co-localization of POSTN-expressing fibroblasts and PTK7 expressing tumor cells (**Suppl. Fig. 16L, M**). While these interactions in NAF are largely restricted to the emergent population, the ligands and receptors are widely expressed across the malignant epithelium in the tumor cells, suggesting that a signaling relationship initially compartmentalized within a defined niche becomes broadly co-opted during tumor progression.

Together, these spatial data provide a critical link between the molecular states identified by single-cell multiomics and their organization within intact tissue. Emergent cells occupy a defined position within the epithelial architecture, residing near the parabasal region where the normal differentiation hierarchy first breaks down, and are embedded within a niche whose signaling relationships are localized by spatial proximity. The conservation of these signaling relationships in tumor tissue directly links the transcriptional identity and communication network surrounding emergent cells in the cancerized field to the microenvironmental changes seen in malignancy and further suggests that the emergent state may be an intermediate between normal tissue homeostasis and cancer.

## Discussion

The traditional view of field cancerization is informed by the concept of multistep carcinogenesis, in which the gradual accumulation of cancer-driving lesions eventually overwhelms the tumor suppressive processes in a tissue and drives neoplastic growth. Within this framework, field cancerization represents an early step: a tumor-adjacent, morphologically normal region that harbors numerous cancer-associated mutations, poised for secondary tumor formation or recurrence^3,10,11^. The reliance on the multistep concept made sense, for what else could explain the mutational abundance in these tumor surrounding fields? However, an alternative source of mutations has emerged over the past decade, as deep sequencing across human organs has shown that lesions in cancer driver genes are surprisingly common and well tolerated in morphologically normal tissues, including healthy oral mucosa^83^.

To resolve what distinguishes a field from the normal tissue that does not transform, we comprehensively profiled matched tumor, normal-adjacent field (NAF), contralateral normal (CN), and blood samples from HPV-negative HNSCC patients using whole genome sequencing, targeted cancer gene sequencing, single-nucleus multiomics and spatial transcriptomics. Our genomic DNA analyses revealed that while NAF tissues harbor expanded mutant clones, these mutations rarely overlap with those driving the adjacent tumors, arguing against a model of shared ancestry. Single-nuclear transcriptomic and chromatin profiling uncovered a deeper layer of alteration: NAF epithelium exhibits disrupted differentiation hierarchy, with aberrant retention of basal programs in differentiating cells, loss of terminal differentiation, expansion of proliferative capacity into normally post-mitotic compartments, and the emergence of a novel cell state that transcriptionally bridges normal epithelium and tumor. These findings, validated at the protein level, with spatial transcriptomics and through functional proliferation assays, demonstrate that field cancerization involves coordinated transcriptional and epigenetic reprogramming of epithelial organization, independent of the specific mutations present.

Previous studies of field cancerization in HNSCC have identified genetic changes, particularly TP53 mutations and chromosomal alterations, in tumor-adjacent tissues and proposed that these accumulate in a stepwise fashion toward malignancy^10,11^. Our four-tissue sampling strategy per patient, combined with both WGS and targeted sequencing enhanced by UMI-based error correction and laser capture microdissection, provided the sensitivity and specificity to rigorously test this model. WGS revealed no consistent genomic differences between NAF and CN. Both harbored scattered mutations at low allele frequencies, lacked chromosomal instability, and were dominated by endogenous clock-like mutational signatures consistent with normal somatic mosaicism^17,18^. Targeted sequencing did identify significantly higher variant allele frequencies in NAF compared to CN, confirming clonal expansion in the tumor-adjacent field, yet these expanded clones were largely distinct from those driving the adjacent tumors. These results align with recent studies demonstrating that driver mutations are present at low frequency throughout morphologically normal oral and skin epithelium^83,84^, and that overlapping mutations between normal margins and tumors are rare, with detectable genetic alterations emerging primarily in tissues that have already acquired focal dysplasia ^13^. Collectively, our genomic data indicate that while clonal expansion occurs in the cancerized field, it is not driven by a shared mutational program with the adjacent tumor, pointing instead to non-genetic features of the tumor-adjacent epithelial fields and the microenvironment that create permissive conditions for expansion.

Consistent with this idea, studies of other cancer types have shown that transcriptional and epigenetic reprogramming can precede genetic alterations in early tumorigenesis^85,86^. To capture these changes at cellular resolution, we performed single-nucleus RNA+ATAC-seq on matched tumor, NAF, and CN tissues from eight patients. Previous single-cell studies of HNSCC have largely focused on the tumor microenvironment, with most datasets capturing fewer than 10% epithelial cells despite these being the cells of origin in this cancer ^20–28^, and even fewer studies include normal tissues adjacent to or distant from the tumor^22,23,26^. By optimizing tissue dissociation and nuclear isolation, we achieved approximately 40% epithelial recovery, yielding over 60,000 high-quality epithelial nuclei and enabling detection of rare cell populations that conventional approaches would likely miss.

To interpret changes in the cancerized field, we first established a molecular map of normal oral epithelial differentiation using CN tissues. Epithelial cells formed a continuous hierarchy from quiescent basal stem cells through a transient, proliferative parabasal compartment to terminally differentiated suprabasal cells, consistent with the hierarchical model of epithelial organization^32,33^. We identified NOTCH1 as the molecular marker of the parabasal compartment, and defined the transcription factor cascades, chromatin dynamics, and microenvironmental signals that coordinate progression through this hierarchy. To connect these intrinsic programs to microenvironmental cues, we developed ContextNet, a cell-cell communication framework that leverages paired RNA and ATAC measurements to construct context-specific gene regulatory networks, yielding more tissue-appropriate signaling predictions than literature-derived approaches.

Against this baseline, NAF epithelium showed clear disruption. Basal stem cell markers were aberrantly retained in differentiating cells, terminal differentiation programs failed to fully activate, and proliferating cells appeared in suprabasal layers normally restricted to post-mitotic cells. This was functionally confirmed by EdU pulse-labeling experiments. The proliferative zone in NAF was significantly expanded relative to CN, with a spatial pattern similar to overtly dysplastic tissue. At the chromatin level, the transcription factor programs defining each compartment of the normal hierarchy, including TP63/TP73 in basal cells, AP-1 factors driving suprabasal differentiation, and parabasal-associated YBX1 and NFYB, were coordinately reduced in NAF, replaced by aberrant accessibility at RUNX family and WNT effector motifs implicated in epithelial-to-mesenchymal transition and invasion^66–68^. These changes were reinforced by rewiring of signaling in the microenvironment. Differentiation-maintaining signals were diminished, while convergent EGFR-directed and WNT signaling emerged across compartments. Rather than selectively disrupting a single layer of the epithelium, field cancerization is thus a result of changes in the regulatory architecture governing the entire differentiation trajectory.

Beyond broad disruption of the differentiation hierarchy, we identified a distinct epithelial state enriched in NAF that we termed the emergent state. These cells expressed tumor-associated markers including *KRT17*, *VIM*, *ZEB2*, and *ETS1*, and occupied an intermediate position in transcriptional space between normal epithelium and tumor, suggesting a novel, molecularly distinct tumor-advancing state arising within genomically normal tissue. At the chromatin level, emergent state cells had lost the master regulators of stratified squamous epithelial identity, TP63 and TP73, and gained coordinated accessibility at ETS, RUNX, and CREB/ATF family motifs associated with invasion, stress response, and cell fate transitions. ContextNet analysis revealed that emergent state cells received coordinated inputs from multiple stromal and immune cell types. This alignment of intrinsic regulatory reprogramming and extrinsic microenvironmental reinforcement suggests that the emergent state is not a stochastic byproduct of transcriptional noise, but a coordinated cellular program sustained by the cancerized field. Finally, we demonstrated that the emergent state cells are localized and distributed throughout the parabasal layer in the tumor adjacent fields using spatial transcriptomics.

These findings have implications for how we understand and detect early neoplastic changes in HNSCC. Current clinical assessment of tissue adjacent to tumors relies on histological evaluation, which by definition cannot detect the molecular alterations we describe here. The specific features we identified, including disrupted epithelial hierarchy, expanded proliferative zones, and emergent state cells expressing tumor-associated markers, represent candidate molecular biomarkers that could identify at-risk tissue that appears normal by conventional pathology. KRT17, VIM, and ZEB2 in particular merit investigation as markers assessable by immunohistochemistry. An important unresolved question is the directionality of these changes: whether transcriptional reprogramming in NAF arises cell-autonomously and creates a permissive environment for clonal expansion and transformation, or whether paracrine signals from the adjacent tumor induce these changes in otherwise normal epithelium. Our study, like all cross-sectional designs, cannot distinguish between these possibilities, though this distinction may not be critical, as both scenarios implicate the same molecular features as early indicators of field cancerization. Additionally, our cohort was restricted to HPV-negative HNSCC, which is characterized by distinct mutational landscapes and clinical behavior compared to HPV-positive disease. Whether similar non-genetic mechanisms of field cancerization operate in HPV-positive tumors, where viral oncoproteins provide an alternative route to epithelial disruption, remains to be determined.

This study has several limitations. Our cohort, while deeply profiled across multiple modalities, represents a limited number of patients, and validation in larger cohorts will be necessary to confirm the generalizability of these findings. Our analyses are restricted to a single anatomic site within the oral cavity, and whether the specific hierarchy disruptions we observe extend to other mucosal squamous epithelia where field cancerization has been described remains unknown. The computational predictions from ContextNet, while benchmarked against independent datasets with measured signaling activity, are correlative and will require experimental perturbation to establish causal regulatory relationships. Longitudinal studies tracking NAF tissues over time, particularly in patients undergoing surveillance, would help establish whether the emergent state cells identified here represent true precursor populations or stable altered states that do not progress. Finally, whether the molecular features of field cancerization we describe are reversible is an open question with therapeutic implications: if the transcriptional and epigenetic reprogramming that defines the cancerized field can be reversed, this could open avenues for developing therapies for patients at high risk of developing HNSCC^87,88^

## Methods

### Tissue collection and processing

Tissues were collected at the University of Washington through the Otolaryngology Surgical Department and NWBioSpecimen from 21 patients (IRB 10571). All patient characteristics are outlined in **Suppl. Table S1**. Assays performed per patient and tissue type are detailed in **Suppl. Table S2**. No statistical methods were used to determine sample size since this study was exploratory. NWBioSpecimen obtained written informed consents from all patients and all data provided to researchers was de-identified. Tumor samples were collected from the Pathology service and provided to NWBioSpecimen. Normal-adjacent field (in two cases were determined to be dysplasia on pathological review) and contralateral samples were taken by the surgeon and given to NWBioSpecimen. A peripheral blood sample was collected by NWBioSpecimen in collaboration with the anesthesia team. Tissues were stored in HypoThermosol at 4°C and transported by the research team to Fred Hutchinson Cancer Center on ice.

HPV status was determined by P16 immunohistochemistry (P16 clone 16P04,JC2, BioSB cat.#BSB5829) with Head and Neck Pathologist review. Computational validation was performed by aligning unmapped reads from single-cell sequencing and WGS reads to HPV reference genomes (HPV16, 18, 31, 33, 35) using Bowtie2; no viral reads were detected in any sample. All samples included in this study were HPV-negative.

Samples for WGS and targeted-seq were embedded in OCT without fixation to avoid fixation-induced mutagenesis and then flash-frozen and stored at −80°C. Samples for single cell multiomics were processed fresh within 24 hours of collection. One patient (Pt44) demonstrated markedly reduced genomic complexity across all molecular assays (mean 15 SVs vs 227 in other patients, minimal CNV and SNV burden) despite HPV-negative status and histopathologic confirmation as HNSCC. Given the atypical genomic profile with discordant pathology review, this patient was excluded from genomic analyses but included in transcriptomic analyses where tumor cells clustered appropriately with other HNSCC samples.

Tissue sections used for spatial transcriptomics contained either contralateral normal epithelium or both tumor and adjacent epithelium on a single section, which was reviewed by a Head and Neck pathologist. The adjacent epithelium demonstrated reactive atypia without dysplasia.

### Whole Genome Sequencing

<u>Laser capture microdissection (LCM):</u> Flash-frozen tissue samples were sectioned, stained with H&E, and reviewed and annotated by a University of Washington head and neck pathologist. Annotations were made for the epithelial cell regions of normal-adjacent and contralateral tissues and cancerous regions of tumor tissues. These annotations served as a guide for which areas to isolate via LCM. On the day of microdissection, 10-12 sections of 10 µm thick tissues were sectioned and mounted onto PEM permeable membranes. Sections were rapidly fixed with 100% ethanol and stained with H&E to guide dissection. 50 µL of Qiagen buffer ATL was added to a 1.5mL tube cap and loaded as the collection vessel. Epithelial regions were isolated with the Leica LMD6 LCM microscope at 10X magnification in the Experimental Histopathology Shared Resource of the Fred Hutchinson Cancer Center.

<u>DNA isolation, library preparation, and sequencing:</u> After epithelial cell isolation and centrifugation, DNA was extracted using the QIAamp DNA Blood Mini (Qiagen #51104) kit. DNA samples of 20ng-200ng (quantified using Qubit Fluorometer) were used in library preparation and sequencing. Library preparation was completed using the Illumina DNA Prep (#20060060) for Pt32, Pt34 and IDT xGen cfDNA & FFPE DNA Library Preparation Kit (#10010206) for all other samples. All tissue samples (T, NAF, CN) were sequenced to at least 60X coverage of the genome and blood samples were sequenced to 40X coverage. Sequencing was conducted using Illumina NovaSeq^TM^ 6000 flow cells using 2×150 base-pair paired-end reads (Illumina library) or 2×158 (IDT).

<u>Data analysis:</u> Initial WGS preprocessing and variant calling were completed using the nf-core/sarek^89,90^ (v3.2.3) pipeline with Nextflow^91^ (v23.10.1) on Cirro^92^. Sequencing data were aligned to the human reference genome hg38 using BWA-MEM^93^ (v0.7.17-r1188). PCR duplicates were marked using GATK^94^ MarkDuplicates (v4.4.0.0). Base quality score recalibration was performed using GATK^94^ BaseRecalibrator and ApplyBQSR (v4.4.0.0). Matched peripheral blood samples were used as normal controls to call somatic single nucleotide variant (SNV) mutation and insertion and deletion (indel) candidates. The tools used include: MuTect2 following GATK^94^ best practices (v4.4.0.0), Strelka^95^ (v2.9.10) with default settings, and VarScan^96^ (v2.4.4) with default parameters and strand filter disabled per developer recommendations. Variants were normalized using BCFtools^97^ norm (v1.19) to left-align indels and ensure consistent variant representation across callers.

Consensus somatic mutations were defined as variants called by MuTect2^98^ and at least one additional variant caller (Strelka^95^ or VarScan^96^). Variants were filtered to retain those with consensus variant allele frequency (VAF) ≥5%. Reported VAFs are from MuTect2 output. All mutations were annotated with Ensembl VEP^99^ (v106.1). Copy number variants were called using ASCAT^100^ (v3.1.1). Missense variants were annotated with AlphaMissense^16^ pathogenicity predictions using the AlphaMissense_hg38 database (accessed December 2025). Mutational signatures were extracted de novo using SigProfilerExtractor^101^ (v1.1.24) with SigProfilerMatrixGenerator (v1.2.28) in Python 3.9.19. Nonnegative matrix factorization was performed testing 1-10 signatures with 100 NMF replicates and reference genome hg38. Extracted signatures were decomposed into known COSMIC signatures^14^. All downstream analyses were performed in Python 3.12.5 by importing VCF files to pandas dataframes using scikit-allel (v1.3.8).

Mutation burden was calculated as somatic mutations per megabase (Mb) of callable genome. Rather than using a fixed genome size, we calculated sample-specific callable bases to account for possible coverage variability introduced by different library preparation kits (Illumina DNA Prep for patients 32 and 34; IDT xGen for remaining patients). Callable regions were defined as positions with sequencing depth ≥10× using mosdepth^102^ (v0.3.8), a threshold commonly applied for reliable variant detection^103^. Mutation burden was then calculated as:

*Mutations/Mb = (number of consensus somatic mutations) / (sample-specific callable Mb)*

Statistical analyses were performed in Python 3.12.5 using SciPy (v1.11.4). For comparisons of VAF distributions across tissue types, the Kruskal–Wallis H-test was used as an omnibus test followed by pairwise two-sided Mann–Whitney U tests. P-values were corrected for multiple comparisons using Bonferroni correction (α = 0.05/3 = 0.017 for three pairwise comparisons). Sample sizes for whole genome sequencing AlphaMissense analysis: tumor n = 230 variants, NAF n = 148 variants, CN n = 100 variants. Sample sizes for targeted sequencing: tumor n = 77 variants, NAF n = 27 variants, CN n = 38 variants.

### Targeted DNA Sequencing

<u>Library preparation and sequencing:</u> Libraries generated from the IDT xGen cfDNA & FFPE DNA Library Preparation Kit (#10010206) protocol were used as input to the IDT xGen Pan-Cancer Hybridization Panel (#1056205). WGS samples previously sequenced with the Illumina DNA Prep kit were re-prepared using the IDT xGen cfDNA & FFPE kit then used as input for panel enrichment. The xGen Pan-Cancer Panel targets 127 significantly mutated genes spanning 800 kb of the human genome, based on genes identified by The Cancer Genome Atlas (TCGA) network across 12 tumor types, including genes commonly mutated in HNSCC such as TP53, PIK3CA, CDKN2A, and NOTCH1^15^. Libraries were prepared with dual 8-base unique molecular identifiers (UMIs) for PCR duplicate removal and error correction. All tissue samples (T, NAF, CN) and matched peripheral blood samples were sequenced to an average coverage of 5,000X on an Illumina NovaSeq^TM^ X using 2×151 base-pair paired-end reads.

<u>Data analysis:</u> UMI processing and consensus sequence generation were performed using fgbio (v2.0.2). Raw reads were extracted from uBAM files and UMIs were annotated from read names (read configuration: 8B+T 8B+T, dual 8-base UMIs on both read 1 and read 2). Reads were grouped by UMI and mapped position using GroupReadsByUmi, and consensus reads were called from UMI families using CallDuplexConsensusReads to generate duplex consensus sequences. Consensus reads were aligned to the human reference genome hg38 using BWA-MEM (v0.7.17-r1188). Matched peripheral blood samples were used as normal controls to call somatic single nucleotide variant (SNV) and insertion and deletion (indel) candidates using three independent callers: MuTect2 from GATK best practices (v4.4.0.0) including LearnReadOrientationModel, Strelka (v2.9.10), and VarDict (v1.8.3). Variants were normalized using BCFtools norm (v1.19) to left-align indels and ensure consistent variant representation across callers. Consensus somatic mutations were defined as variants called by MuTect2 and at least one additional variant caller. Variants were filtered to retain only PASS variants and those with variant allele frequency (VAF) ≥2% and at least 5 alt reads. Reported VAFs are from MuTect2 output. All mutations were annotated with Ensembl VEP (v110.1). Similar to WGS analysis, missense variants were annotated with AlphaMissense pathogenicity predictions using the AlphaMissense_hg38 database. All downstream analyses were performed in Python 3.12.5 by importing VCF files to pandas dataframes using scikit-allel (v1.3.8).

### Single cell multiomic RNA+ATAC data generation

<u>Tissue processing:</u> An overview is outlined in **Suppl. Fig. 4A**. Tissues were washed 3x with cold HBSS (Gibco #14175095) to remove excess blood and finely minced on ice using 2 scalpels on a glass coverslip inside a 6 well plate. They were then transferred to a new 6 well plate (~50mg per well) with HBSS, 0.5% Collagenase 1 (Gibco #170100-017), 1.26 mM CaCl_2_ (Fisher #10128-558), and 200 µg/mL DNAse I (Roche #10104159001). They were incubated for 1.5 hours at 37°C, 75 rpm and carefully pipetted every 15 min. Samples were strained with a 70µm strainer (Miltenyi #130-098-462). Residual tissue on the strainer was collected and digested for 10 minutes in Trypsin-EDTA (0.25%) at 37°C, 75 rpm (Gibco #25200114) then inactivated with 10% FBS (Genesee #25-550) and added to the tube containing the original digested cells. Samples were centrifuged and assessed for viability. Samples with lower viability were processed using a Dead Cell Removal Kit (Miltenyi #130-090-101). Samples were washed and centrifuged twice with PBS (Gibco #14190144)+0.05% BSA (Miltenyi #130-091-376).

<u>Nuclei isolation:</u> Next, cells were lysed for 2.33 min on ice using a fresh lysis buffer containing 10mM Tris-HCl (pH 7.4 Sigma #T2194), 10mM NaCl (Sigma #59222C), 3mM MgCl_2_ (Sigma #M1028), 0.1% IGEPAL (Sigma #i8896), 0.1% Tween-20 (Bio-Rad #1662404), 1mM DTT (Sigma #646563), 1 U/µL RNase inhibitor (Sigma #3335402001), 0.01% Digitonin (Fisher #BN2006), 1% BSA, Nuclease-free Water (Fisher #10977015). Nuclei were then carefully washed 2-3 times using a wash buffer containing 10mM Tris-HCl, 10mM NaCl, 3mM MgCl_2_, 0.1% Tween-20, 1 mM DTT, 1 U/µL RNase inhibitor, 1% BSA, Nuclease-free Water. Nuclei were counted on a hemocytometer with trypan blue staining. If significant debris or nuclei clumping observed, nuclei were strained (Flowmi # BAH136800040-50EA) and recounted.

Note, all centrifugations were performed using swing bucket rotor at 4°C for 10 min. Cell centrifugations were performed at 300 g. Nuclei centrifugations were performed at 500 g. All low volume steps were performed using lobind tubes (1.5 mL Eppendorf #13698791; 0.2 mL USA Scientific #1402-4700). Lobind tips were used throughout the protocol (Rainin #30389166, 30389188, 30389190, 30389173). Nuclei were pipetted slowly throughout isolation and processing to preserve nuclear integrity.

<u>Library preparation and sequencing:</u> Nuclei were processed using the Chromium Next GEM Single Cell Multiome ATAC + Gene Expression Reagent Kit (10X Genomics, #1000285, CG000338 Rev C) targeting ~10,000 nuclei per sample following the manufacturer’s protocol. Briefly, nuclei were transposed, barcoded in Gel Bead-in-Emulsion (GEMs) on a Chromium Controller, and pre-amplified. The pre-amplified product was split for parallel ATAC and Gene Expression library construction. ATAC libraries were indexed using Single Index Kit N Set A and Gene Expression libraries were indexed using Dual Index Kit TT Set A. Libraries were assessed using Agilent TapeStation and quantified using Qubit Fluorometer. Libraries were sequenced on Illumina NextSeq 2000 targeting 25,000 read pairs per cell (ATAC) and 20,000 read pairs per cell (Gene Expression).

### Single-cell multiomics data analysis

Preprocessing: FASTQ files were processed using Cell Ranger ARC (v2.0.0-2.0.2) for demultiplexing, alignment to the human reference genome hg38, and count matrix generation.

### RNA

<u>RNA quality control and filtering:</u> Genes detected in fewer than 50 cells were excluded. Cells with greater than 40% mitochondrial reads were removed. Doublets were identified and removed using Scrublet^104^ (v0.2.3). We captured 145,769 nuclei after quality control, with a mean of 6,626 cells per sample. Following filtering, gene expression data was normalized per cell using Scanpy^105^ (v1.10.2) so that all cells would have the same total count over all genes. A pseudocount of 0.1 was added and log transformed with base 2.

<u>Dimensionality reduction and clustering:</u> The top 1,500 highly variable genes (HVGs) were identified using the Cell Ranger flavor. Principal component analysis (PCA) was performed using 50 components on the HVGs. For multi-patient analyses, batch correction was performed using Harmony^106^ (v0.0.9) using Scanpy’s harmony_integrate function to correct for patient-specific effects. A k-nearest neighbor (kNN) graph was constructed with k=15. Cells were clustered using the Leiden algorithm at default resolution.

<u>Cell type annotation:</u> Cell types were manually annotated based on expression of canonical marker genes and Leiden cluster assignments ^20,21,25,28,107^. We identified clusters of epithelial, endothelial, fibroblast, myofibroblast, B, T, myeloid, melanocyte, myocyte, and neuronal cells. Marker genes used are included in **Suppl. Table S6**. Cell types are shown in **Suppl. Table S7**.

<u>Copy number variation inference:</u> Copy number variations were inferred from single cell gene expression profiles using infercnvpy^29,30^ (v0.4.3) with immune cells as the reference population, a sliding window size of 250 genes, and excluding sex chromosomes (chrX, chrY) and mitochondrial chromosome (chrMT). CNV scores were calculated for each cell to distinguish malignant from non-malignant epithelial cells.

Cell cycle scoring: Cell cycle phase was assigned using Scanpy’s score_genes_cell_cycle function with standard S phase and G2/M phase gene lists, generating S scores, G2/M scores, and phase assignments for each cell.

<u>Differential gene expression:</u> Differential expression analysis across cell types was performed using Scanpy’s rank_genes_groups function with the Wilcoxon rank-sum test, comparing each cell group against all other cells. A pseudocount of 1 was added to the expression matrix prior to testing. P-values were adjusted for multiple testing using the Benjamini–Hochberg procedure. Genes were ranked by log fold change, and those with a minimum log fold change of 1.25 were considered differentially expressed.

<u>Diffusion maps and MAGIC imputation:</u> Diffusion maps were computed using Palantir^47^ (v1.4.1) to capture the underlying manifold structure of the data. The resulting diffusion operator was used to perform MAGIC^108^ imputation via Palantir, which leverages the diffusion kernel to denoise and impute gene expression values by sharing information across similar cells. Imputed expression values were used for visualization of gene expression patterns.

<u>Pseudotime:</u> Pseudotime trajectories were inferred using Palantir. Diffusion maps were computed on the PCA representation using 10 diffusion components. The starting cell was selected as the cell with the most basal-like transcriptional state based on diffusion component extrema, and the terminal cell was selected as the most differentiated cell using the same approach. Palantir was then run with these specified start and terminal states to compute pseudotime values and fate probabilities for each cell. Gene expression dynamics along pseudotime were characterized using Palantir’s gene trend computation. Gene trends were z-score normalized for visualization.

<u>Gene correlation analysis:</u> To identify genes co-expressed with marker genes of interest (KRT15, KRT13, NOTCH1), Pearson correlation coefficients were computed across all gene pairs using the normalized expression matrix. For each marker gene, positively correlated genes were identified, and their expression dynamics were visualized along pseudotime.

<u>Cell-state Density:</u> Cell-state density in the differentiation manifold was estimated using Mellon^45^ (v1.6.1). The density estimator was applied to diffusion map eigenvectors using the fractal dimension method (d_method=“fractal”). Log-density values were computed for each cell, with visualization values clipped at the 5th percentile to reduce the influence of outliers in sparse regions of the manifold.

<u>Sample diversity:</u> To quantify local sample heterogeneity, we computed neighborhood entropy as a measure of sample diversity. For each cell, the 15 nearest neighbors were identified in Harmony-corrected PCA space using scikit-learn’s NearestNeighbors (v1.1.3). Shannon entropy was then calculated over the sample label distribution within each cell’s neighborhood. Higher entropy values indicate neighborhoods with cells from multiple samples (high sample diversity), while lower values indicate neighborhoods dominated by cells from a single sample.

<u>Cell type label transfer:</u> Epithelial cell type labels were transferred from contralateral cells to normal adjacent cells using k-nearest neighbors classification. A k-NN model (k=30) was fit on the Harmony-corrected PCA coordinates of contralateral cells with known cell type annotations. For each normal adjacent cell, the 30 nearest neighbors among contralateral cells were identified, and the most frequent cell type label among neighbors was assigned as the predicted label.

<u>Kompot Differential Abundance, Differential Expression, and Gene Set Enrichment Analysis:</u> Diffusion maps were computed with Palantir using 10 diffusion components derived from the Harmony-corrected PCA representation, and this diffusion map embedding served as the latent cell-state representation for all Kompot^63^ (v0.6.2) analyses. Differential abundance was computed across conditions (NAF-CN) with Kompot using default parameters (ls_factor = 10, n_landmarks = 5000) for pairwise comparison between Normal-Adjacent Field vs. Contralateral. Differential expression (DE) was performed on the same comparisons with Kompot defaults (ls_factor = 10, n_landmarks = 5000, sigma = 1). For DE on the RNA dataset, we used the Scanpy-normalized expression matrix described above, applying a log_2_ transformation to normalized counts after adding a 0.1 pseudocount, and included all 16,524 genes in the analysis. Statistical significance was determined using a null distribution of 2,000 permuted genes to estimate false discovery rates (FDR). Gene set enrichment analysis was carried out using the Kompot StringDBReport function, which interfaces with the STRING^72^ database through its API. Gene names were mapped to STRING protein identifiers using the default mapping, unmapped genes were skipped, and enrichment was computed using STRING’s default annotation categories for any gene set derived from the Kompot DE results.

<u>Tumor Similarity Analysis:</u> To quantify the transcriptional distance of each cell from healthy epithelium, we computed a tumor similarity score using iterative pseudotime analysis. Palantir pseudotime measures biological distance through shortest paths in a nearest-neighbor graph built on diffusion map cell-state representations; cells requiring a longer path from a given start are more transcriptionally distant from it. We computed pseudotime 100 times on the co-embedded epithelial data containing CN, NAF, and tumor cells, each time selecting a different randomly sampled cell from the CN compartment as the starting cell (Palantir parameters: knn = 10, use_early_cell_as_start = True). For each cell, we retained the minimum pseudotime value across all 100 iterations. Because pseudotime increases monotonically with biological distance from the start, the minimum over many healthy-epithelium starts represents the closest any healthy cell comes to a given cell in transcriptional space, and therefore a robust lower bound on that cell’s biological distance from the healthy compartment. Cells consistently far from all healthy starting points, i.e., tumor-like cells, thus receive high minimum pseudotime values, yielding a score where higher values reflect greater transcriptional similarity to tumor. Because the score is a graph distance from the healthy compartment rather than a directed distance toward tumor, cells that diverge from healthy epithelium in any direction receive high values; intermediate values are therefore obtained only by cells positioned between the healthy and tumor compartments on the manifold.

### ATAC

<u>ATAC processing:</u> ATAC data were processed using ArchR^109^ (Setty Lab fork, https://github.com/settylab/ArchR) aligning to the hg38 genome. Arrow files were generated from fragment files with filtering parameters: minimum TSS enrichment score of 1, minimum fragment count of 500, and exclusion of mitochondrial chromosome. ATAC analysis was restricted to cells that passed RNA quality control filters. Dimensionality reduction was performed using iterative Latent Semantic Indexing (LSI) on the tile matrix, followed by clustering and UMAP embedding. Peaks were called using ArchR’s reproducible peak set method and a peak matrix was generated for downstream analyses. Gene activity scores were computed using ArchR’s gene scoring model, which infers regulatory potential based on chromatin accessibility near gene bodies weighted by distance to the transcription start site. Transcription factor motif accessibility was assessed using chromVAR^48^ with the CIS-BP motif database^110^.

For analyses of non-tumor cells, batch correction was performed using Harmony via Scanpy’s harmony_integrate function on the LSI coordinates to correct for patient-specific effects. Tumor cells were excluded from batch correction due to substantial copy number alterations that cannot be appropriately corrected. Diffusion maps were computed using Palantir on either the uncorrected or Harmony-corrected LSI coordinates as appropriate, and diffusion-based imputation was applied to impute sparse chromatin accessibility signals for visualization.

<u>ChromVAR pseudotime heatmap:</u> To identify and visualize chromVAR scores which changed over pseudotime in the contralateral tissue, a Mellon function estimator (sigma = 1, ls = .2) was trained to predict chromVAR scores using RNA pseudotime. To enrich for transcription factors whose motif accessibility varied along pseudotime, we retained factors for which the mean squared error of the fit was at least 0.2 log-fold lower than the overall variance of their chromVAR scores, ensuring that pseudotime explained a meaningful fraction of the signal. We further required that each transcription factor be expressed in more than 500 cells. For visualization, transcription factors were ordered by the pseudotime position at which their imputed chromVAR score reached its maximum.

<u>Kompot differential ChromVAR:</u> Diffusion maps were computed with Palantir using the top 4 diffusion components derived from the Harmony-corrected LSI representation of non-tumor cells. This diffusion map embedding served as the latent cell-state representation for all Kompot^63^ (v0.6.2) analyses performed on the ATAC data. Differential motif accessibility was computed across conditions (NAF vs. CN) using Kompot with default parameters and chromVAR scores as the input layer. A log fold change cutoff of 0.1 was applied to highlight motifs with consistently higher or lower accessibility between conditions.

<u>Differential ChromVAR analysis:</u> To identify chromVAR scores which were enriched in specific cell types (**Fig. 5H**) we used sc.tl.rank_genes_groups (method=“Wilcoxon”, layer=“chromVAR”). The top 30 markers for each group were selected, and the mean of their chromVAR score within a cell type were plotted.

### ContextNet: Context-specific cell-cell communication inference

Cell-cell communication inference aims to identify which ligands from sender cells drive transcriptional changes in receiver cells. NicheNet^31^ addresses this by modeling signaling as information flow through three integrated network layers: (i) ligand-receptor interactions that initiate signaling, (ii) intracellular signaling pathways that propagate the signal, and (iii) a gene regulatory network (GRN) that links signaling intermediates to target genes. Using Personalized PageRank, NicheNet computes a ligand-target regulatory potential matrix and scores each ligand’s ability to predict an observed gene set of interest (e.g., differentially expressed genes in receiver cells) using AUPRC.

NicheNet’s gene regulatory network derives from aggregated prior knowledge across diverse biological contexts. While this provides broad coverage, gene regulatory relationships are inherently context-specific. The same transcription factor can activate different target genes in different tissues due to cell-type-specific chromatin accessibility and enhancer usage. Single-cell multiomics data, jointly profiling gene expression (snRNA-seq) and chromatin accessibility (snATAC-seq), enables direct inference of context-specific regulatory relationships from experimental data, presenting an opportunity to improve ligand-target predictions.

We developed ContextNet to leverage this opportunity by integrating multiomics-derived GRN with NicheNet’s signaling inference framework (**Suppl. Fig. 5B**). ContextNet replaces NicheNet’s generic GRN with a context-specific GRN inferred from paired snRNA-seq and snATAC-seq data, while preserving NicheNet’s curated ligand-receptor interactions and signaling pathways (**Suppl. Fig. 5C**). This grounds ligand-target predictions in experimentally observed regulatory relationships specific to the biological system under study.

The context-specific cell-cell communication inference comprises of three components: (1) GRN inference that constructs data-derived TF-target relationships from multiomics data, (2) ContextNet, an R package that integrates the data-derived GRN with NicheNet or MultiNicheNet to performs ligand activity prediction, and (3) ccflowR for visualizing signaling flow using Sankey diagrams.

### Context-specific gene regulatory network construction

We constructed context-specific GRNs using a multiomics workflow (**Suppl Fig. 5C**) that links TFs to target genes through cis-regulatory elements^111^. The approach integrates three lines of evidence: (i) correlation between gene expression and nearby chromatin accessibility, (ii) presence of TF binding motifs in accessible regions, and (iii) correlation between TF expression and chromatin accessibility at those motifs. The output is a gene × TF matrix where each entry represents the regulatory weight linking a TF to a target gene.

*<u>Input data</u>*: The input is matched snRNA-seq and snATAC-seq profiles from the same cells. Chromatin accessibility peaks (regions of open chromatin) were identified from snATAC-seq data using ArchR^112^. To reduce technical sparsity inherent in single-cell data while preserving biological heterogeneity, we aggregated cells into metacells using SEACells^113^. SEACells groups transcriptionally or epigenomically similar cells into representative cellular states, averaging out technical noise while retaining meaningful variation across cell types and conditions. All subsequent correlations were computed across metacells rather than individual cells.

*<u>Linking peaks to target genes</u>*: We assigned candidate cis-regulatory peaks to potential target genes based on genomic proximity. For each gene *g*, *P*_g_ comprised peaks overlapping the genomic interval spanning its longest annotated transcript and extending 100 kb upstream and downstream, consistent with previous enhancer– promoter interaction studies^111,112^. For each candidate gene–peak pair, we calculated the Spearman correlation r(*g*, *p*) between gene expression and peak accessibility across metacells. Each observed correlation was evaluated against a background distribution generated using 100 peaks matched for GC content and total accessibility with the built-in SEACells procedure. We defined the binary gene–peak association *B*_g,*p*_ as

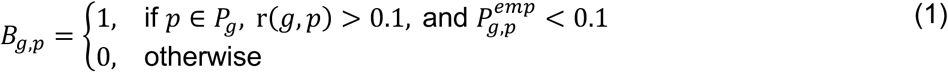

where *P^emp^*is the unadjusted empirical one-sided value obtained from the matched background distribution. To reduce potentially spurious associations driven by a single peak, subsequent analyses were restricted to genes linked to at least two retained peaks:

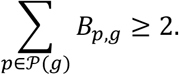

*<u>Prioritize TF-motif links using TF expression</u>*: To identify potential TF-binding sites within gene associated peaks, we extracted a 150-bp sequence centered on the midpoint of each peak interval and scanned each region against position weight matrices from the CIS-BP database^110^ using FIMO^114^. For each match, FIMO assigns a p-value representing the probability of obtaining an equal or higher motif score by chance under the background sequence model. Motif matches with p < 1 × 10⁻⁴ were retained.

To distinguish functional binding from spurious motif matches, we utilized the in-silico ChIP framework ^111^. in-silico ChIP integrates motif scanning results with correlation between TF expression and accessibility at the predicted binding site to compute a score that reflects the likelihood of functional TF binding at each peak. Positive correlation suggests the TF likely contributes to chromatin accessibility at that site, consistent with functional binding. For TF *t* and peak *p*, the scores are computed by in-silico ChIP as follows

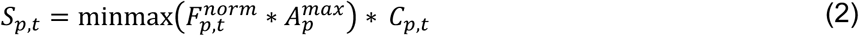

where,

- *F_p_*_,*t*_*^norm^* is the normalized FIMO score such that *F*_∗,*t*_*^norm^* ∈ {0, 1} for each TF *t*
- *A_p_^max^* is the maximum accessibility of peak *p* across metacells
- minmax normalizes values to {0, 1}
- *C_p_*_,*t*_ is the correlation between accessibility of peak *p* and TF *t*. *C_p_*_,*t*_ is set to 0 for negative values

Peaks with *S_p_*_,*t*_ > 0 have both predicting binding sites for TF *t* and accessibility that correlates with expression of TF *t*.

*<u>Assembling the gene regulatory network</u>*: To construct candidate TF–target relationships, we combined the peak-gene associations from Eq. 1 and peak-TF in-silico ChIP scores from Eq. 2. For TF *t* and gene *g*, the regulatory weight was calculated by summing the in-silico ChIP scores across peaks linked with gene *g*:

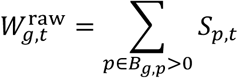

Summing across peaks integrates regulatory support from multiple candidate cis-regulatory elements linking the same TF to the same gene. Regulatory weights were subsequently normalized within each TF:

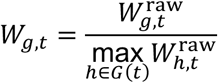

where G is the set of candidate targets associated with each TF t.

This normalization scales the target-gene weights for each TF to [0,1], providing a common range for network integration while preserving the relative ranking of targets within each TF.

### Integration with NicheNet

ContextNet integrates the context-specific GRN with NicheNet’s prior knowledge networks (**Suppl Fig 5B-C**). NicheNet organizes its networks into two structures: **lr_network**, containing ligand-receptor interactions, and **weighted_networks**, which combines an intracellular signaling network (**sig**) linking receptors to downstream signaling molecules and a gene regulatory network (**gr**) linking signaling molecules and transcription factors to target genes. During integration, we replaced NicheNet’s generic gene regulatory network (**gr**) with the context-specific GRN constructed above. NicheNet’s ligand-receptor interactions (**lr_network**) and intracellular signaling network (**sig**) were retained without changes. This design leverages curated databases for ligand-receptor biology and signaling pathways while using context-specific inference for gene regulatory relationships where tissue-specific enhancer usage dominates. NicheNet’s ligand-activity framework is used to evaluated how well each ligand’s predicted target ranking recovered the receiver-cell gene set relative to the background gene set. Ligands are then ranked by their background-corrected area under the precision–recall curve (AUPRC) using the same approach in NicheNet.

### ContextNet Benchmarking

#### Dataset with positive controls

To evaluate the performance of using context-specific GRN on ligand prioritization, we benchmarked ContextNet against NicheNet in human PBMC DOGMA-seq dataset^115^ with a defined anti-CD3/CD28 perturbation (**Suppl Fig. 6**).

Since the dataset contained no defined sender population, ligand availability could not be attributed to a cellular source. We therefore removed direct ligand–receptor edges from NicheNet’s weighted signaling network and initiated propagation at the receptors, allowing receptor-to-target predictions to be evaluated independently of the ligand layer. Using NicheNet’s built-in propagation function and the fitted cutoff and damping parameters, each receptor was propagated through the intracellular signaling network to TFs. The resulting receptor-to-TF matrix was shared between the two methods and multiplied by either the stimulated-PBMC ContextNet GRN or NicheNet’s literature-based GRN.

Receptors detected in at least 10% of stimulated cells were eligible for ranking. The stimulation-response program comprised genes detected in at least 10% of cells, increased after stimulation with log2FC > 0.5 and associated with a Bonferroni-adjusted P ≤ 0.05. Both methods ranked the same 170 receptors, with lower ranks indicating stronger recovery of the stimulation-response program.

CD28 and CD3D were prespecified as perturbation-related controls, whereas CD40LG and CD69 were prespecified as early activation-associated controls (**Suppl. Fig. 6B**).

#### Lineage specificity of predicted transcription factor target programs

To assess whether the HNSCC-derived regulatory network captured context-specific transcription-factor (TF) programs, we computed target-program activity of lineage specific TFs across all 145,769 cells in our dataset. For each TF, we compared target programs derived from ContextNet with those derived from the literature-based regulatory prior (NicheNet GRN). Each network was first restricted to target genes represented in our dataset. The remaining targets were ranked by regulatory weight, and maximum of 150 genes were retained to compute TF activity scores (**Fig. 2G**, **Suppl. Fig. 5A**).

#### Matched comparison of ligand prioritization in basal epithelial cells

We compared ContextNet and NicheNet on their ability to recapitulate known signaling pathways in basal epithelial cells (**Fig. 2H**). Genes with log2FC >= 0.25 and associated with a Bonferroni-adjusted P < 0.05 in basal cells compared to other epithelial cells were defined as the receiver program. Cell types with at least 500 cells were used as sender populations. Genes detected in at least 20% of cells defined the expressed ligand, receptor and background gene sets. The top 30 ligands from each method were used for evaluation. Note that the only difference between ContextNet and NicheNet in this evaluation is the GRN used.

Biological relevance was assessed using a prespecified, non-exhaustive reference set of 12 ligands. This set comprised six basement-membrane or pericellular-matrix ligands (COL4A2, COL6A2, COL12A1, COL15A1, COL18A1 and LAMA4), three Wnt ligands or regulators (WNT4, WNT9A and SFRP1), and three additional basal-state paracrine signals (DLL1, CXCL12 and MIF). These ligands reflected three processes relevant to basal epithelial maintenance: matrix interactions that support basal-cell anchorage and basement-membrane organization, Wnt signaling that regulates differentiation, and Notch or stromal–epithelial signaling. ContextNet ranked 11 of the 12 reference ligands in the top 30, whereas NicheNet ranked only two reference ligands (**Fig. 2H**).

### Visualization of signaling across receiver populations with ccflowR

To compare signaling across receiver populations in a single view, we developed **ccflowR**, an R framework that displays selected cell-cell communication routes as a four-layer Sankey diagram connecting sender cell types, ligand, receptor and receiver cell types. Joint visualization therefore reveals which signals are shared across receiver cell types, which are specific to individual receivers, and which sender populations are the most likely sources.

ccflowR first computes an edge weight for each sender-ligand-receptor-receiver combination. This width represents the width of the flow if only one interaction was visualized. The individual weights are combined across all interactions for joint visualization.

#### Calculation of signaling flow weights

*Single-condition:* Weights for each sender-ligand-receptor-receiver combination are computed by combining ligand expression in the sender cell type, predicted receptor activity in the receiver cell type and prior ligand-receptor weights.

Consider a sender cell type *s*, ligand *l*, receptor *r*, and receiver cell type *c*. Ligand expression is z-scored across cells and converted to probabilities per cell type and is computed as:

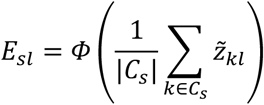

where C*_s_* is the set of cells in sender cell type *s*, *z_kl_* is the clipped, z-scored expression of ligand *l* in cell *k* and Φ is the standard normal cumulative distribution function. *E_sl_* ranges between [0, 1].

Receiver-specific ligand activity was quantified using ContextNet AUPRC. Let *Q_Pc_* denote the AUPRC of ligand *P* for receiver cell type *c*. Normalized ligand activity was calculated as:

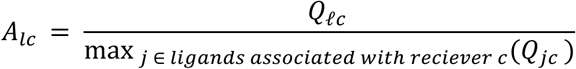

The activity score for each sender *s*, ligand *l*, and receiver *c* is then computed as

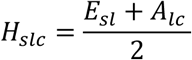

Higher activity scores represent ligands that are both preferentially expressed in the sender cell type and strongly predictive of the target gene set in the receiver cell type.

Since multiple receptors can contribute to an interaction between sender and receiver cell types through the same ligand, the prioritization score for a sender-ligand-receptor-receiver combination is computed as follows. If *P_Pr_* denotes the NicheNet prior weight of ligand–receptor pair (*l*, *r*), the normalized weight is computed as

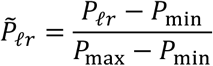

The weight for each sender-ligand-receptor-receiver combination is then computed as

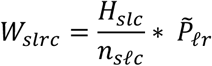

Where *n_sPc_* is the number of receptors associated with ligand *l* that led to a predicted interaction between sender *s* and receiver *c*.

*<u>Multi-condition:</u>* MultiNicheNet assigns each sender–ligand–receptor–receiver–condition combination a composite prioritization score integrating ligand and receptor differential expression, their statistical support and pseudo bulk expression, ligand activity, and the fractions of cells expressing the ligand and receptor. This composite prioritization score is used in place of the activity score (*H_slc_*) defined above for the single-condition analysis. The weight for each sender-ligand-receptor-receiver combination is then computed as

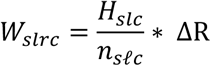

Where ΔR is the scaled receptor log fold change toward the condition.

#### Sankey diagram visualization

The ccflowR Sankey diagram comprises of four node columns (sender cell types, ligands, receptors, receiver cell types) and connected by three edges (sender-to-ligand, ligand-to-receptor, receptor-to-receiver). Together, these three edges formed a complete sender–ligand–receptor–receiver route. The thickness of each route is determined by *W_slrc_*. Node height represents the total incoming or outgoing route. Routes are colored by receiver cell type in single-condition analyses or by the favored condition in multi-condition analyses.

### Application of ContextNet to infer cell-cell communication

To characterize the cell–cell communication landscape associated with HNSCC field cancerization, we applied ContextNet using context-specific epithelial gene regulatory networks (GRNs). We analyzed contralateral and Normal adjacent tissues separately to identify communication patterns within each tissue and find interactions that differed between the two conditions.

<u>Context-specific gene regulatory networks</u>: We constructed two epithelial GRNs from the matched multiome data: (i) CN-specific GRN was used for the CN single-condition analysis and (ii) NAF GRN, constructed from epithelial cells in both CN and NAF tissues, was used for the NAF single-condition and CN-versus-NAF analyses. Both gene × TF matrices contained 16,524 genes and 748 TFs.

<u>ContextNet application to CN tissues:</u> CN-specific GRN was used for analysis of contralateral normal tissues (**Fig. 3**). Cell types represented by more than 500 cells were eligible senders and included endothelial, fibroblast, myeloid, myofibroblast, T cell, basal, dividing, parabasal and suprabasal populations. Basal, parabasal, dividing and suprabasal epithelial cells were analyzed as receivers. Ligand activity was calculated separately for each receiver cell type. Genes with log₂ fold-change threshold of 0.25, expression in at least 5% of cells and Bonferroni-adjusted *P* < 0.05 were used as the receiver-specific gene set for ligand-activity prediction. Genes expressed in at least 20% of cells were included in the expressed ligand, receptor and background-gene sets. The top 30 ligands ranked by AUPRC were retained for downstream analysis for each receiver cell type. These predictions were combined with the two sender cell types with the highest ligand expression for each ligand-receptor pair for visualization with ccflowR (**Suppl. Fig. 9A**). Differentially expressed genes in receiver cell types in the top 500 predicted targets in the ligand-target matrix were nominated as candidate genes influenced by external signaling (**Suppl. Fig. 9B**) if they were TFs with motifs enriched in CN (**Fig. 3J**) or epithelial cell state makers (**Fig. 3H**),

<u>ContextNet application to NAF tissues</u>: We used the same approach used for CN tissue for analysis of NAF tissue (**Fig. 5**) running ContextNet with the NAF GRN. The same sender and receiver populations used for CN analysis were used with the addition of emergent cells a receiver population. Given the focus on emergent cells which are fewer in number, we increase the differential gene expression fold change to 0.5 for receiver-specific gene set and decreased the threshold to 15% of cells for background gene sets. Similar to CN analysis, top 30 ligands ranked by AUPRC were retained for downstream analysis for each receiver were combined with the two sender cell types with the highest ligand expression for visualization with ccflowR (**Suppl. Fig. 13A-B**). Differentially expressed genes in emergent cell type in the top 500 predicted targets in the ligand-target matrix were nominated as candidate genes influenced by external signaling (**Suppl. Fig. 13C**) if they were TFs with motifs enriched in NAF (**Fig. 4I**) or genes upregulated in NAF (**Fig. 4E**).

<u>Differential signaling between CN and NAF:</u> Differential communication between CN and normal adjacent field (NAF) tissues was determined using MultiNicheNet^69^ and the context-specific NAF GRN. Basal, parabasal, suprabasal, and dividing cells were used as receivers. Emergent cells were not included in this analysis. Cell types with at least 10 cells in at least two separate samples in each condition were used as sender cell types. The 10 eligible sender populations were endothelial, fibroblast, melanocyte, myeloid, myofibroblast, T, basal, dividing, parabasal, and suprabasal cells. CN was specified as the condition of interest and NAF as the reference condition. We used a log₂ fold-change threshold of 0.5, a P-value threshold of 0.05, a minimum expression fraction of 5%, and the top 250 predicted targets per ligand. The top highest-ranked routes were retained for visualization (**Fig. 4J**). Differentially expressed genes between CN and NAF (**Fig. 4E**) in the top 500 predicted targets in the ligand-target matrix were nominated as candidate genes influenced by external signaling (**Suppl. Fig. 11A-B**) if they were TFs with differential motif accessibility (**Fig. 4I**) or epithelial cell state markers.

### Tissue slice culture and EdU proliferation assay

<u>EdU Pulse Labeling:</u> Specimens were collected in HypoThermosol FRS (Stem Cell #7935) medium and transported on ice. Once sterilized, specimens were handled in biosafety hood at all times. Specimens were sterilized by submerging in 1% povidone-iodide/HBSS solution and washed 3 times with gentle rotation in room temperature HBSS. Specimens were fitted on a sterilized brain matrix, epithelial side up and stromal side down, and were cut to 1 mm thick slices using a sterilized razor blade. Specimen slices were pulsed in 250 µM EdU (Invitrogen #C10338) in KSFM media (Gibco #10744019) solution at 37°C for 2 hours. After the 2-hour incubation, all specimens were fixed (4% PFA/PBS, overnight, 4°C), cryoprotected (30% sucrose/PBS solution, overnight, 4°C) and frozen in OCT blocks.

<u>Immunofluorescence (IF):</u> All washing steps were performed in 0.1% Triton/PBS at room temperature, 5 min with gentle rotation unless noted otherwise. Refer to **Supplemental Table S13** for antibody details and dilutions. Tissues frozen in OCT blocks were sectioned to 8µm thickness and collected on Superfrost^+^ (Fisher Scientific #1255015) slides. Sections were washed twice in PBS and permeabilized with 0.3% Triton/PBS (room temperature, 20 min with gentle rotation). Permeabilized sections were washed once and rimmed with a PAP pen (Vector Labs #H-4000) and blocked in Blocking Solution at RT for 1 hour (1% BSA, 1% Gelatin, 2% Goat Serum, 0.1% Triton in PBS). Sections were incubated in primary antibody (diluted in Blocking Solution) overnight at 4°C in a humidified chamber. The next day, sections were washed twice with 0.1% Triton/PBS, once with PBS, and stained with Click-iT EdU Cell proliferation kit (Invitrogen #C10338) (room temperature, 30 min in the dark). Sections were washed 3 times before incubation with secondary antibody and DAPI (1:1000) diluted in Blocking Solution (room temperature, 30 min in the dark). Sections were washed twice with 0.1% Triton/PBS, once with PBS and mounted with ProLong Gold (Invitrogen #P36930) or Fluoromount-G (Southern Biotech #0100-35). IF stained sections were imaged at 20X or 40X on Zeiss LSM 700 or Leica Stellaris 8-STED confocal microscopes.

<u>Quantification of EdU uptake from basement membrane:</u> Images of tissue sections were analyzed using IMARIS software. Using the internalized shape recognition tool (“spheres”), EdU positive nuclei were individually selected and their closest distance to basement membrane (as defined by KRT14 signal) was calculated. Statistical significance was assessed using Welch’s ANOVA, followed by Games-Howell post hoc pairwise comparisons to account for multiple testing (α = 0.05).

<u>Spindle orientation:</u> Z-stacks of 20 µm thick sections stained with Aurora B, Ki67, KRT15 and DAPI were imaged on Andor Dragonfly spinning disk confocal microscope at 40X magnification. Daughter nuclei pairs in telophase were identified by the presence of an Aurora B positive midbody positioned between two Ki67 positive nuclei. Cells in anaphase were identified by alignment of Aurora B signal along the central spindle and positive Ki67 signal. Division angle was defined as the angle between the plane of the basement membrane and the line connecting the centers of two daughter nuclei. Angles were measured using the angle tool and quantified with the Measure function in Fiji. Data were analyzed and statistical tests were performed using ordinary one-way ANOVA in Prism (GraphPad).

### Single Cell Spatial Transcriptomics Data Generation

We processed HPT-32 and HPT-34 on a single slide that included three small skin tissues for a separate project, using a prerelease version of the Illumina Spatial Transcriptome Assay (StrataMap^TM^). First, sections of HPT-34 were used for a permeabilization optimization assay. Sections were fixed in methanol, H&E stained, and brightfield imaged using the Leica DMi8. Tissue permeabilization was performed at 5 to 25 minutes across six wells on a thermocycler set to 37 degrees. First strand synthesis was performed on the substrate followed by tissue digestion and RNA removal before a linear amplification step in the presence of fluorescently labeled nucleotide within the 6 wells to generate a detectable signal. Fluorescent images of each tissue (time point) were analyzed using the ROI method in Leica’s LASX software v.3.7.5 and the highest mean fluorescent intensity was used to select a 10-minute permeabilization time.

<u>Permeabilization optimization</u>: Pt32M and Pt34M (mixed tumor/NAF tissue blocks) were processed on a single slide using <u>a</u> prerelease version of the Illumina Spatial Transcriptome Assay (StrataMap^TM^). Initial sections of Pt34 were used for permeabilization optimization. Sections were fixed in methanol, H&E stained, and brightfield imaged on a Leica DMi8. Tissue permeabilization was performed across six wells on a thermocycler at 37°C for 5-25 minutes. First strand synthesis was performed on the substrate followed by tissue digestion and RNA removal before <u>a linear</u> amplification step in the presence of fluorescently labeled nucleotide within the 6 wells to generate a detectable signal. Fluorescent images of each tissue (time point) were analyzed using the ROI method in Leica’s LASX software v.3.7.5 and the highest mean fluorescent intensity was used to select a 10-minute permeabilization time. In a second experiment, Pt32C and Pt34C were processed with (200073552 Prerelease Draft B - 12/15/2025), which removed permeabilization testing steps, and used a universal 10 minutes for all tissues.

<u>Library preparation and sequencing:</u> The transcriptome assay substrate was similarly fixed, H&E stained and imaged at 20X. Each tissue was imaged separately and assessed using the Illumina Spatial Imaging Tool (ISIT) software which evaluated fiducial quality to provide an accurate spatial map of the tissue. We proceeded with tissue permeabilization and synthesized first stand DNA, complementing the spatially barcoded mRNA transcripts on the surface of the substrate. Next, the tissue was digested and RNA was removed by buffer washes containing NaOH. Amplification of the second strand was performed on the substrate. All the eluted cDNA was used to generate a library of spatially barcoded DNA from all tissues on the slide. We identified the appropriate cycle number based on the qPCR signal of the eluted cDNA. The library was cleaned using Illumina Purification Beads and sized with Agilent’s Tape Station 4150. The library was quantified by qPCR before sequencing on an Illumina NovaSeq^TM^ X Plus using all eight lanes of the 25B reagent kit. The <u>BCL files</u> and images were uploaded processed with the Illumina BioInsight Platform Core cloud-based bioinformatic pipeline to map transcripts back to each sample.

### Single Cell Spatial Transcriptomics Data Analysis

<u>Cell segmentation and preprocessing:</u> Raw sequencing and image data were uploaded to the Illumina Connected Analytics platform where initial cell segmentation was performed. Data were then downloaded, converted to SpatialData format^116^, and re-segmented using Proseg^117^ (v3.1.0), which leverages transcript data to improve segmentation accuracy.

Transcript diffusion is a major source of technical noise in interpretation of spatial transcriptomics data^118^. Diffusion of transcripts between neighboring cells during the data generation process leads to mis-assignment of transcripts to unrelated cell types with the result being equivalent to doublets in single-cell data that contain a mix of transcripts from two or more distinct cell types. We therefore used SPLIT to identify and correct such errors in our spatial data. SPLIT uses RCTD for identification of cells that require correction^118,119^. RCTD uses cell-type annotated reference single-cell data to simulate synthetic doublets. Using this reference, each spatial cell is labeled either as a singlet with primary cell type or a doublet that is a mixture of a primary and secondary cell types. SPLIT purifies the doublets by removing the transcripts that belong to the secondary cell type. The inferred primary cell type is the one used for cell type annotation of all spatial cells. SPLIT purified spatial expression matrices along with cell type annotation is used for all downstream analysis (**Suppl. Fig. 15E-H**).

RCTD was run using genes shared between spatial data and snRNA-seq reference. The following snRNA-seq datasets were used as references: Pt32 mixed-tissue snRNA-seq for Pt32 mixed-tissue spatial sample and Pt32 contralateral snRNA-seq for Pt32 contralateral sample. Since Pt34 was not part of the patient cohort profiled using single-cell multiomics, Pt32 snRNA-seq was used as reference for the Pt34 spatial samples as well. RCTD model was run with UMI_min = 10, counts_MIN = 10, UMI_min_sigma = 100, and CELL_MIN_INSTANCE = 25. RCTD outputs were subsequently used as input to SPLIT to for cell-type-aware purification to generate purified spatial expression matrices that were used for all downstream analyses.

<u>Integration of spatial and snRNA-seq data:</u> Spatial epithelial profiles from Pt32 and Pt34 contralateral (CN) and normal-adjacent field (NAF) regions were integrated with matched CN and NAF epithelial cells from snRNA-seq using Harmony^106^. This included 37,779 snRNA-seq cells, 97,052 spatial cells, and 13,079 shared genes (**Suppl. Fig. 16A-D**).

snRNA raw counts and SPLIT purified raw counts were log-normalized in Seurat (v5.4.0) using LogNormalize with a scale factor of 10,000. Highly variable genes were selected from snRNA-seq reference, rather than recomputed. This gene set was intersected with genes shared across both modalities, yielding 1,103 highly variable genes.

Since snRNA-seq and spatial measurements have different dynamic ranges and noise profiles, log-normalized expression was z-scored separately within each modality and clipped to −10 to 10 to limit extreme values. PCA was run on the scaled matrix, and the first 50 PCs were used as input to Harmony^106^ (v1.2.4), with modality and sample identity as covariates. The nearest-neighbor graph and UMAPs were computed from the Harmony-corrected PC space.

Spatial epithelial subtype labels were transferred by k-nearest neighbor classification in the Harmony-corrected embedding. Each cell was assigned an epithelial subtype using the majority vote of its 30 nearest snRNA-seq neighbors. Label transfer was evaluated by snRNA-seq subtype-neighborhood preservation, which measured whether the annotation in the reference remained locally coherent after the classification.

Harmony-corrected PCs were also used as input to Mellon for gene expression imputation^45^. Marker gene expression was used only for validation and visualization and not for cell type annotations.

<u>Spatial proximity analysis</u>: To assess the spatial relationship between emergent cells and other epithelial populations, we calculated the minimum Euclidean distance from each emergent cell to the nearest basal, parabasal, and suprabasal cell using a KD-tree based on spatial coordinates. Distances were converted from pixels to micrometers using a scale factor of 0.311 µm/pixel. Only emergent cells with a defined nearest neighbor for all three target types were retained. Overall differences across target cell types were assessed using a Friedman test, followed by pairwise two-sided Wilcoxon signed-rank tests with Benjamini–Hochberg correction for multiple testing.

## Data availability

Raw whole genome and targeted panel DNA sequencing data for this study was deposited in dbGaP under accession number phs004595.v1. Raw single cell RNA and ATAC sequencing data as well as count matrices were deposited in GEO under accession number GSE339273 (https://www.ncbi.nlm.nih.gov/geo/query/acc.cgi?acc=GSE339273). Single-cell spatial transcriptomics data including cellxgene and cellxlocation matrices were deposited in GEO under the same accession number. All processed data objects including all cell metadata and analyses outputs are also available in GEO.

## Code availability

ContextNet (v0.1.0) is available as an R package (R ≥ 4.1.0) at https://github.com/settylab/ContextNet, leveraging nichenetr (v2.0.4)^1^ for network integration and PPR computation. ccflowR provides Sankey visualization with support for NicheNet, MultiNicheNet, and other cell-cell communication frameworks and is available at https://github.com/settylab/ccflowR.

## Acknowledgements

We acknowledge the generous patient donors for contributing their samples to this study. We thank the University of Washington NWBioSpecimen service staff including Sarah Bowell, Aldous Subang, Piper Driskell, and Lauren Wahl, who were instrumental in sample collection. We thank the Fred Hutch Innovation Lab for piloting the Illumina StrataMap^TM^ Spatial Transcriptome Assay and performing all spatial transcriptomics data generation. Illumina graciously provided reagents for spatial transcriptomics assays and supported associated sequencing costs. We also would like to thank Dr. Jon Cooper for providing various antibodies for our immunofluorescence experiments. We would like to thank Pritha Chanana and Matthew Fitzgibbon for assistance in data deposition. We would also like to thank the nf-core community for developing the nf-core infrastructure and resources for Nextflow pipelines.

This project was supported by the 2021 Fred Hutchinson Cancer Center Human Biology Pilot Award, 2023 Fred Hutchinson Cancer Center Translational Data Science Integrated Research Center New Collaboration Award, Dick & Loraine Burger Pilot Funding, 2026 Andy Hill CARE Fund, NIH/NCI Cancer Center Support Grant P30 CA015704 of the Fred Hutch/University of Washington/Seattle Children’s Cancer Consortium which includes the Cellular Imaging Shared Resources, RRID:SCR_022609, NIH grants R01-AR070780 (S.B), R35-GM147125 (M.S), ORIP S10OD028685 to support high-performance computing at Fred Hutch and the RSG-19-074-01-TBE award (S.B) from the American Cancer Society. C.M. was funded by the University of Washington Oral Health Sciences T90 Comprehensive Training in Inter-Disciplinary Oral Health Research Training Grant (5T90DE021984-14). S.H. was supported by the Fred Hutchinson Cancer Center Interdisciplinary Training in Cancer Research Program. C.F. was supported by NIH F31 (1HD118770-01). D.O. was supported by NIH K99 (GM159044-01). R.K.L was supported by American Cancer Society Cancer Stem Cell Consortium Postdoctoral fellowship (CSCC-PF-23-994844).

## Author Contributions

Conceptualization: C.M., M.S., S.B.; Methodology: C.M., S.H., C.M.L., C.F., D.O., S.M., R.R., P.I., S.S., G.H.; Validation: C.M., C.M.L., R.K.L., M.S., S.B.; Formal analysis: C M., S.H., C.F., D.O., C.M.L.; Investigation: C M., C.M.L., H.W-F., L.S., E.N.; Resources: A.B., B.B., M.S., S.B.; Data curation: C.M., S.M., C.M.L.; Writing – original draft: C.M., M.S., S.B.; Writing - review & editing: C.M., R.R., M.S., S.B.; Visualization: C.M., C.M.L., S.H., C.F., D.O.; Supervision: M.S., S.B.; Project administration: M.S., S.B.; Funding: C.M., M.S., S.B.

## References

1. Slaughter, D. P., Southwick, H. W. & Smejkal, W. “Field cancerization” in oral stratified squamous epithelium. Clinical implications of multicentric origin. Cancer 6, 963–968 (1953).

2. Mohan, M. & Jagannathan, N. Oral field cancerization: An update on current concepts. Oncol. Rev. 8, 13–19 (2014).

3. Curtius, K., Wright, N. A. & Graham, T. A. An evolutionary perspective on field cancerization. Nat. Rev. Cancer 18, 19–32 (2017).

4. Weckx, A. et al. Time to recurrence and patient survival in recurrent oral squamous cell carcinoma. Oral Oncol. 94, 8–13 (2019).

5. Johnson, D. E., et al. Head and neck squamous cell carcinoma. Nat. Rev. Dis. Primers 6, (2020).

6. Shetty, K. S. R. et al. Salvage Surgery in Recurrent Oral Squamous Cell Carcinoma. Frontiers in Oral Health 2, (2022).

7. Athanassios, A., Karamouzis, M. V, Raben, D. & Ferris, R. L. Head and neck cancer. The Lancet 371, 1695–1709 (2008).

8. Leemans, C. R., Tiwari, R., Nauta, J. J. P., Waal, I. Van Der & Snow, G. B. Recurrence at the primary site in head and neck cancer and the significance of neck lymph node metastases as a prognostic factor. Cancer 73, 187–190 (1994).

9. Brennan, J. et al. Molecular Assessment of Histopathological Staging In Squamous-Cell Carcinoma of the Head and Neck. New England Journal of Medicine 332, 429–435 (1995).

10. Tabor, M. P. et al. Persistence of genetically altered fields in head and neck cancer patients: Biological and clinical implications. Clinical Cancer Research 7, 1523–1532 (2001).

11. Califano, J. et al. Genetic progression model for head and neck cancer: Implications for field cancerization. Cancer Res. 56, 2488–2492 (1996).

12. Braakhuis, B. J. M., Tabor, M. P., Kummer, J. A., Leemans, C. R. & Brakenhoff, R. H. A genetic explanation of slaughter’s concept of field cancerization: Evidence and clinical implications. Cancer Res. 63, 1727–1730 (2003).

13. Wu, P. et al. The genomic architectures of tumour-adjacent tissues, plasma and saliva reveal evolutionary underpinnings of relapse in head and neck squamous cell carcinoma. Br. J. Cancer 125, 854–864 (2021).

14. Alexandrov, L. B. et al. The repertoire of mutational signatures in human cancer. Nature 578, 94–101 (2020).

15. Lawrence, M. S. et al. Comprehensive genomic characterization of head and neck squamous cell carcinomas. Nature 517, 576–582 (2015).

16. Cheng, J. et al. Accurate proteome-wide missense variant effect prediction with AlphaMissense. Science (1979). 381, (2023).

17. Yokoyama, A. et al. Somatic mosaicism in the buccal mucosa reflects lifestyle and germline risk factors for esophageal squamous cell carcinoma. Sci. Transl. Med. 17, eadq6740 (2025).

18. Martincorena, I. et al. Somatic mutant clones colonize the human esophagus with age. Science (1979). 362, 911–917 (2018).

19. Slyper, M. et al. A single-cell and single-nucleus RNA-Seq toolbox for fresh and frozen human tumors. Nat. Med. 26, 792–802 (2020).

20. Puram, S. V. et al. Single-Cell Transcriptomic Analysis of Primary and Metastatic Tumor Ecosystems in Head and Neck Cancer. Cell 171, 1611–1624.e24 (2017).

21. Kürten, C. H. L. et al. Investigating immune and non-immune cell interactions in head and neck tumors by single-cell RNA sequencing. Nat. Commun. 12, (2021).

22. Choi, J. H. et al. Single-cell transcriptome profiling of the stepwise progression of head and neck cancer. Nat. Commun. 14, (2023).

23. Xiong, M. et al. Single-cell sequencing of head and neck carcinoma: Transcriptional landscape and prognostic model based on malignant epithelial cell features. FASEB Journal 38, (2024).

24. Bill, R. et al. CXCL9:SPP1 macrophage polarity identifies a network of cellular programs that control human cancers. Science (1979). 381, 515–524 (2023).

25. Puram, S. V. et al. Cellular states are coupled to genomic and viral heterogeneity in HPV-related oropharyngeal carcinoma. Nat. Genet. 55, 640–650 (2023).

26. Meng, X. et al. Single-Cell Analyses Reveal the Metabolic Heterogeneity and Plasticity of the Tumor Microenvironment during Head and Neck Squamous Cell Carcinoma Progression. Cancer Res. 84, 2468–2483 (2024).

27. Quah, H. S. et al. Single cell analysis in head and neck cancer reveals potential immune evasion mechanisms during early metastasis. Nat. Commun. 14, (2023).

28. Mair, F. et al. Extricating human tumour immune alterations from tissue inflammation. Nature 605, 728–735 (2022).

29. Tickle, T. T. I. G. C. B. M. H. B. inferCNV of the Trinity CTAT Project. Preprint at https://github.com/broadinstitute/inferCNV (2019).

30. Sturm, G. infercnvpy: Infer copy number variation (CNV) from scRNA-seq data. Preprint at https://github.com/icbi-lab/infercnvpy (2021).

31. Browaeys, R., Saelens, W. & Saeys, Y. NicheNet: modeling intercellular communication by linking ligands to target genes. Nat. Methods 17, 159–162 (2020).

32. Potten, C. S. THE EPIDERMAL PROLIFERATIVE UNIT: THE POSSIBLE ROLE OF THE CENTRAL BASAL CELL. Cell Prolif. 7, 77–88 (1974).

33. Blanpain, C. & Fuchs, E. Epidermal homeostasis: a balancing act of stem cells in the skin. Nat. Rev. Mol. Cell Biol. 10, 207–217 (2009).

34. Tumbar, T. et al. Defining the Epithelial Stem Cell Niche in Skin. Science (1979). 303, 359–363 (2004).

35. Potten, C. S. & Morris, R. J. Epithelial stem cells in vivo. J. Cell Sci. 1988, 45–62 (1988).

36. Clayton, E. et al. A single type of progenitor cell maintains normal epidermis. Nature 446, 185–189 (2007).

37. Doupé, D. P., Klein, A. M., Simons, B. D. & Jones, P. H. The Ordered Architecture of Murine Ear Epidermis Is Maintained by Progenitor Cells with Random Fate. Dev. Cell 18, 317–323 (2010).

38. Mascré, G. et al. Distinct contribution of stem and progenitor cells to epidermal maintenance. Nature 489, 257–262 (2012).

39. Takeda, T. et al. Immunohistological evaluation of Ki-67, p63, CK19 and p53 expression in oral epithelial dysplasias. Journal of Oral Pathology & Medicine 35, 369–375 (2006).

40. Dunaway, S. et al. Divide and conquer: two stem cell populations in squamous epithelia, reserves and the active duty forces. Int. J. Oral Sci. 11, 26 (2019).

41. Andl, C. D. et al. Association of TGFβ signaling with the maintenance of a quiescent stem cell niche in human oral mucosa. Histochem. Cell Biol. 146, 539–555 (2016).

42. Khanom, R. et al. Expression of basal cell keratin 15 and keratin 19 in oral squamous neoplasms represents diverse pathophysiologies. Histol. Histopathol. 27, 949–59 (2012).

43. Harris, A., et al. Human-Specific Organization of Proliferation and Stemness in Squamous Epithelia: A Comparative Study to Elucidate Differences in Stem Cell Organization. Int. J. Mol. Sci. 26, (2025).

44. Sakamoto, K. et al. Down-regulation of keratin 4 and keratin 13 expression in oral squamous cell carcinoma and epithelial dysplasia: a clue for histopathogenesis. Histopathology 58, 531–542 (2011).

45. Otto, D. J., Jordan, C., Dury, B., Dien, C. & Setty, M. Quantifying cell-state densities in single-cell phenotypic landscapes using Mellon. Nat. Methods 21, 1185–1195 (2024).

46. Rangarajan, A. et al. Notch signaling is a direct determinant of keratinocyte growth arrest and entry into differentiation. EMBO J. 20, 3427–36 (2001).

47. Setty, M. et al. Characterization of cell fate probabilities in single-cell data with Palantir. Nat. Biotechnol. 37, 451–460 (2019).

48. Schep, A. N., Wu, B., Buenrostro, J. D. & Greenleaf, W. J. chromVAR: inferring transcription-factor-associated accessibility from single-cell epigenomic data. Nat. Methods 14, 975–978 (2017).

49. Elbediwy, A., Vincent-Mistiaen, Z. I. & Thompson, B. J. YAP and TAZ in epithelial stem cells: A sensor for cell polarity, mechanical forces and tissue damage. BioEssays 38, 644–653 (2016).

50. Piccolo, S., Dupont, S. & Cordenonsi, M. The Biology of YAP/TAZ: Hippo Signaling and Beyond. Physiol. Rev. 94, 1287–1312 (2014).

51. Mahoney, J. E., Mori, M., Szymaniak, A. D., Varelas, X. & Cardoso, W. V. The Hippo Pathway Effector Yap Controls Patterning and Differentiation of Airway Epithelial Progenitors. Dev. Cell 30, 137–150 (2014).

52. Li, J. et al. TEAD1 and TEAD3 Play Redundant Roles in the Regulation of Human Epidermal Proliferation. J. Invest. Dermatol. 140, 2081–2084.e4 (2020).

53. Romano, R.-A. et al. Δ *Np63* knockout mice reveal its indispensable role as a master regulator of epithelial development and differentiation. Development 139, 772–782 (2012).

54. Soares, E. & Zhou, H. Master regulatory role of p63 in epidermal development and disease. Cellular and Molecular Life Sciences 75, 1179–1190 (2018).

55. Di Girolamo, D., Di Iorio, E. & Missero, C. Molecular and Cellular Function of p63 in Skin Development and Genetic Diseases. Journal of Investigative Dermatology 145, 766–779 (2025).

56. Schmuth, M., Moosbrugger-Martinz, V., Blunder, S. & Dubrac, S. Role of PPAR, LXR, and PXR in epidermal homeostasis and inflammation. Biochimica et Biophysica Acta (BBA) - Molecular and Cell Biology of Lipids 1841, 463–473 (2014).

57. Eckert, R. L. et al. AP1 Transcription Factors in Epidermal Differentiation and Skin Cancer. J. Skin Cancer 2013, 1–9 (2013).

58. Welter, J. F. & Eckert, R. L. Differential expression of the fos and jun family members c-fos, fosB, Fra-1, Fra-2, c-jun, junB and junD during human epidermal keratinocyte differentiation. Oncogene 11, 2681–7 (1995).

59. Pontarollo, G. et al. Commensal bacteria weaken the intestinal barrier by suppressing epithelial neuropilin-1 and Hedgehog signaling. Nat. Metab. 5, 1174–1187 (2023).

60. Lefort, K. et al. Notch1 is a p53 target gene involved in human keratinocyte tumor suppression through negative regulation of ROCK1/2 and MRCKalpha kinases. Genes Dev. 21, 562–77 (2007).

61. Nguyen, B.-C. et al. Cross-regulation between Notch and p63 in keratinocyte commitment to differentiation. Genes Dev. 20, 1028–42 (2006).

62. WHO Classification of Tumours Editorial Board. Head and Neck Tumours. vol. 9 (International Agency for Research on Cancer, Lyon, 2024).

63. Otto, D. J. et al. Comparing phenotypic manifolds with Kompot: Detecting differential abundance and gene expression at single-cell resolution. Preprint at 10.1101/2025.06.03.657769 (2025).

64. Zhao, L. et al. Screening and clinical significance of tumor markers in head and neck squamous cell carcinoma through bioinformatics analysis. Mol. Med. Rep. 19, 143–154 (2019).

65. Chan, J. M. et al. Signatures of plasticity, metastasis, and immunosuppression in an atlas of human small cell lung cancer. Cancer Cell 39, 1479–1496.e18 (2021).

66. Chang, W.-M. et al. Dysregulation of RUNX2/Activin-A Axis upon miR-376c Downregulation Promotes Lymph Node Metastasis in Head and Neck Squamous Cell Carcinoma. Cancer Res. 76, 7140–7150 (2016).

67. Tsunematsu, T. et al. RUNX3 has an oncogenic role in head and neck cancer. PLoS One 4, e5892 (2009).

68. AL-Hamad, K. A. Role of RUNX2 in Oral Squamous Cell Carcinoma (OSCC): A Systematic Scoping Review. Eur. J. Cancer Care (Engl). 2024, 1–9 (2024).

69. Browaeys, R. et al. MultiNicheNet: a flexible framework for differential cell-cell communication analysis from multi-sample multi-condition single-cell transcriptomics data. Preprint at 10.1101/2023.06.13.544751 (2023).

70. Kania, A. & Klein, R. Mechanisms of ephrin–Eph signalling in development, physiology and disease. Nat. Rev. Mol. Cell Biol. 17, 240–256 (2016).

71. Byrd, K. M. et al. Heterogeneity within Stratified Epithelial Stem Cell Populations Maintains the Oral Mucosa in Response to Physiological Stress. Cell Stem Cell 25, 814–829.e6 (2019).

72. Szklarczyk, D. et al. STRING v11: protein–protein association networks with increased coverage, supporting functional discovery in genome-wide experimental datasets. Nucleic Acids Res. 47, D607–D613 (2019).

73. Baraks, G. et al. Dissecting the Oncogenic Roles of Keratin 17 in the Hallmarks of Cancer. Cancer Res. 82, 1159–1166 (2022).

74. Mendez, M. G., Kojima, S.-I. & Goldman, R. D. Vimentin induces changes in cell shape, motility, and adhesion during the epithelial to mesenchymal transition. FASEB J. 24, 1838–51 (2010).

75. Fardi, M., Alivand, M., Baradaran, B., Farshdousti Hagh, M. & Solali, S. The crucial role of ZEB2: From development to epithelial-to-mesenchymal transition and cancer complexity. J. Cell. Physiol. 234, 14783–14799 (2019).

76. Adams, J. M. & Cory, S. The Bcl-2 apoptotic switch in cancer development and therapy. Oncogene 26, 1324–37 (2007).

77. Gluck, C. et al. Molecular dissection of the oncogenic role of ETS1 in the mesenchymal subtypes of head and neck squamous cell carcinoma. PLoS Genet. 15, e1008250 (2019).

78. Dittmer, J. The biology of the Ets1 proto-oncogene. Mol. Cancer 2, 29 (2003).

79. Nagarajan, P. et al. Ets1 blocks terminal differentiation of keratinocytes and induces expression of matrix metalloproteases and innate immune mediators. J. Cell Sci. 123, 3566–3575 (2010).

80. Hai, T. & Hartman, M. G. The molecular biology and nomenclature of the activating transcription factor/cAMP responsive element binding family of transcription factors: activating transcription factor proteins and homeostasis. Gene 273, 1–11 (2001).

81. Wang, H. et al. POSTN+ cancer-associated fibroblasts determine the efficacy of immunotherapy in hepatocellular carcinoma. J. Immunother. Cancer 12, e008721 (2024).

82. Dessaux, C., Ganier, L., Guiraud, L. & Borg, J.-P. Recent insights into the therapeutic strategies targeting the pseudokinase PTK7 in cancer. Oncogene 43, 1973–1984 (2024).

83. Lawson, A. R. J. et al. Somatic mutation and selection at population scale. Nature 10.1038/s41586-025-09584-w (2025) doi:10.1038/s41586-025-09584-w.

84. Martincorena, I. et al. High burden and pervasive positive selection of somatic mutations in normal human skin. Science (1979). 348, 880–886 (2015).

85. Aran, D. et al. Comprehensive analysis of normal adjacent to tumor transcriptomes. Nat. Commun. 8, (2017).

86. Teschendorff, A. E. et al. Epigenetic variability in cells of normal cytology is associated with the risk of future morphological transformation. Genome Med. 4, (2012).

87. Bourhis, J. et al. Nivolumab added to cisplatin and radiotherapy versus cisplatin and radiotherapy alone after surgery for people with squamous cell carcinoma of the head and neck at a high risk of relapse (GORTEC 2018-01 NIVOPOST-OP): a randomised, open-label, phase 3 trial. The Lancet 407, 363–374 (2026).

88. Uppaluri, R. et al. Neoadjuvant and Adjuvant Pembrolizumab in Locally Advanced Head and Neck Cancer. New England Journal of Medicine 393, 37–50 (2025).

89. Ewels, P. A. et al. The nf-core framework for community-curated bioinformatics pipelines. Nat. Biotechnol. 38, 276–278 (2020).

90. Garcia, M. et al. Sarek: A portable workflow for whole-genome sequencing analysis of germline and somatic variants. F1000Res. 9, 63 (2020).

91. Di Tommaso, P. et al. Nextflow enables reproducible computational workflows. Nat. Biotechnol. 35, 316–319 (2017).

92. Cirro. Preprint at https://cirro.bio/.

93. Li, H. & Durbin, R. Fast and accurate short read alignment with Burrows–Wheeler transform. Bioinformatics 25, 1754–1760 (2009).

94. Auwera, G. A. V. der. & O’Connor, B. D. Genomics in the Cloud : Using Docker, GATK, and WDL in Terra. (O’Reilly Media, Incorporated, 2020).

95. Kim, S. et al. Strelka2: fast and accurate calling of germline and somatic variants. Nat. Methods 15, 591–594 (2018).

96. Koboldt, D. C. et al. VarScan 2: Somatic mutation and copy number alteration discovery in cancer by exome sequencing. Genome Res. 22, 568–576 (2012).

97. Danecek, P. et al. Twelve years of SAMtools and BCFtools. Gigascience 10, (2021).

98. Benjamin, D. et al. Calling Somatic SNVs and Indels with Mutect2. Preprint at 10.1101/861054 (2019).

99. McLaren, W. et al. The Ensembl Variant Effect Predictor. Genome Biol. 17, 122 (2016).

100. Van Loo, P. et al. Allele-specific copy number analysis of tumors. Proceedings of the National Academy of Sciences 107, 16910–16915 (2010).

101. Islam, S. M. A. et al. Uncovering novel mutational signatures by de novo extraction with SigProfilerExtractor. Cell Genomics 2, 100179 (2022).

102. Pedersen, B. S. & Quinlan, A. R. Mosdepth: quick coverage calculation for genomes and exomes. Bioinformatics 34, 867–868 (2018).

103. Franco, I. et al. Whole genome DNA sequencing provides an atlas of somatic mutagenesis in healthy human cells and identifies a tumor-prone cell type. Genome Biol. 20, 285 (2019).

104. Wolock, S. L., Lopez, R. & Klein, A. M. Scrublet: Computational Identification of Cell Doublets in Single-Cell Transcriptomic Data. Cell Syst. 8, 281–291.e9 (2019).

105. Wolf, F. A., Angerer, P. & Theis, F. J. SCANPY: large-scale single-cell gene expression data analysis. Genome Biol. 19, 15 (2018).

106. Korsunsky, I. et al. Fast, sensitive and accurate integration of single-cell data with Harmony. Nat. Methods 16, 1289–1296 (2019).

107. Williams, D. W. et al. Human oral mucosa cell atlas reveals a stromal-neutrophil axis regulating tissue immunity. Cell 184, 4090–4104.e15 (2021).

108. van Dijk, D. et al. Recovering Gene Interactions from Single-Cell Data Using Data Diffusion. Cell 174, 716–729.e27 (2018).

109. Granja, J. M. et al. ArchR is a scalable software package for integrative single-cell chromatin accessibility analysis. Nat. Genet. 53, 403–411 (2021).

110. Weirauch, M. T. et al. Determination and Inference of Eukaryotic Transcription Factor Sequence Specificity. Cell 158, 1431–1443 (2014).

111. Argelaguet, R. et al. Decoding gene regulation in the mouse embryo using single-cell multi-omics. Preprint at 10.1101/2022.06.15.496239 (2022).

112. Feng, J., Liu, T., Qin, B., Zhang, Y. & Liu, X. S. Identifying ChIP-seq enrichment using MACS. Nat. Protoc. 7, 1728–1740 (2012).

113. Persad, S. et al. SEACells infers transcriptional and epigenomic cellular states from single-cell genomics data. Nat. Biotechnol. 10.1038/s41587-023-01716-9 (2023) doi:10.1038/s41587-023-01716-9.

114. Bailey, T. L., Johnson, J., Grant, C. E. & Noble, W. S. The MEME Suite. Nucleic Acids Res. 43, W39–W49 (2015).

115. Mimitou, E. P. et al. Scalable, multimodal profiling of chromatin accessibility, gene expression and protein levels in single cells. Nat. Biotechnol. 39, 1246–1258 (2021).

116. Marconato, L. et al. SpatialData: an open and universal data framework for spatial omics. Nat. Methods 22, 58–62 (2025).

117. Jones, D. C. et al. Cell simulation as cell segmentation. Nat. Methods 22, 1331–1342 (2025).

118. Bilous, M. et al. Resolving sensitivity, specificity and signal contamination in Xenium spatial transcriptomics. Nat. Methods 23, 1152–1162 (2026).

119. Cable, D. M. et al. Robust decomposition of cell type mixtures in spatial transcriptomics. Nat. Biotechnol. 40, 517–526 (2022).

