## Supplementary Figures and Supplementary Table Legends for "Dysregulation of epithelial differentiation architecture underlies field cancerization in head and neck cancer"

### Supplemental Figures

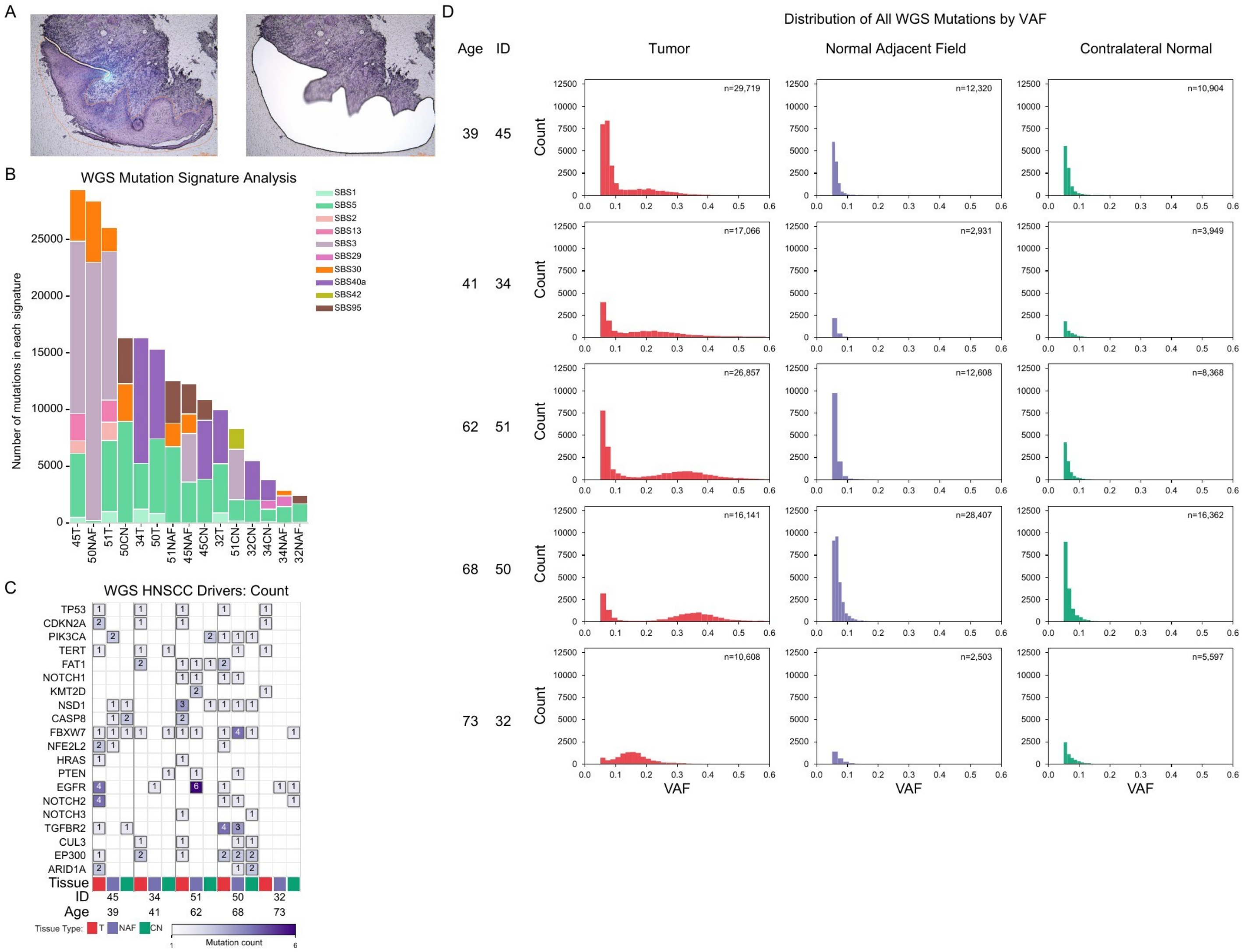

**Supplemental Figure 1. Whole genome sequencing quality metrics and mutational signatures.**

(**A**) Representative images of laser capture microdissection showing isolation of epithelial cells from tissue sections, guided by annotation from a head and neck pathologist.

(**B**) Mutational signature composition across all samples. Signatures were extracted *de novo* using SigProfilerExtractor[1] and decomposed into COSMIC reference signatures[2]. Bar height indicates number of mutations attributed to each signature per sample.

(**C**) Mutation counts in canonical HNSCC driver genes detected by whole genome sequencing across tissue types.

(**D**) Distribution of somatic mutations by VAF for each patient and tissue type. Total mutation count (n) indicated for each sample.

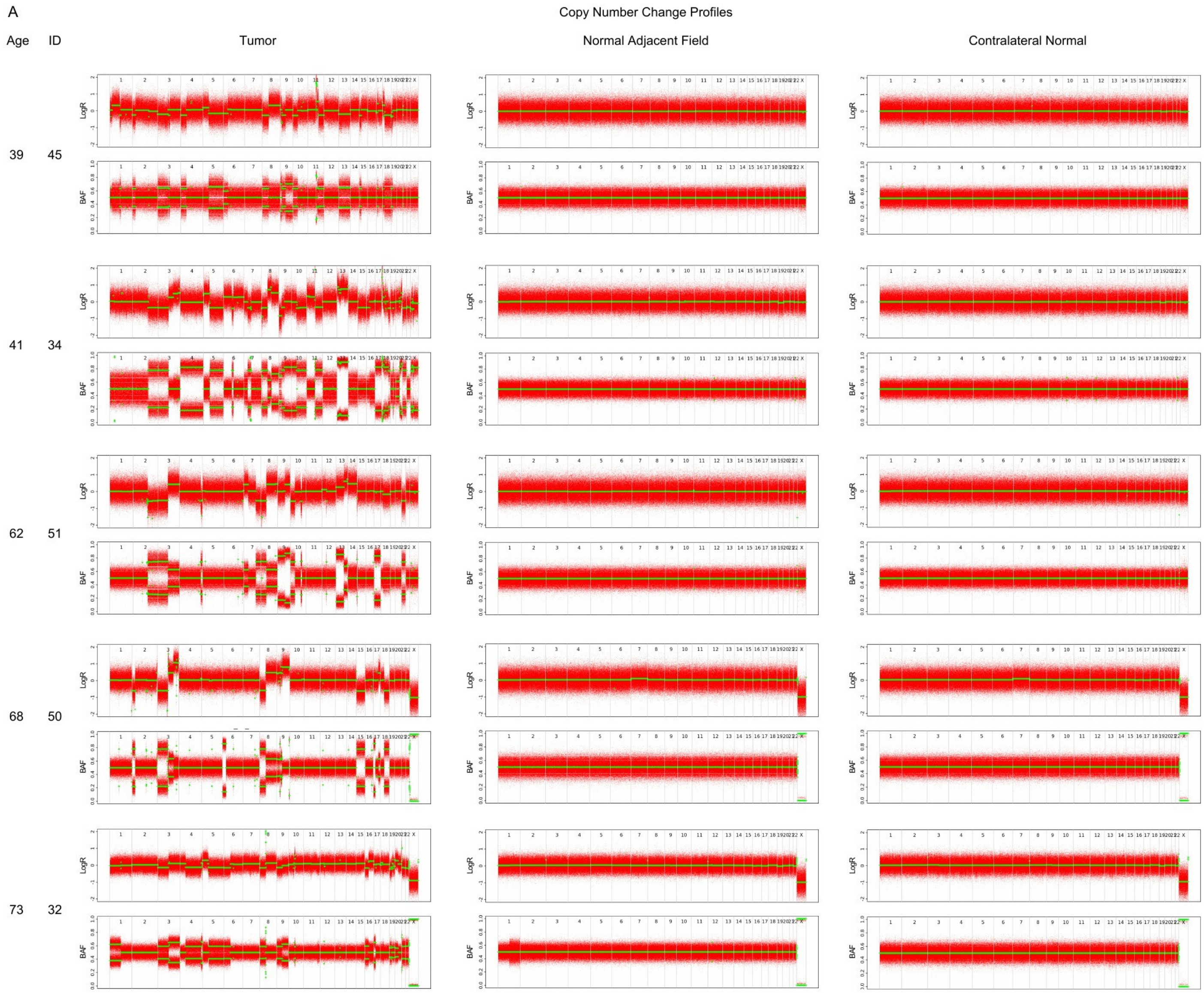

**Supplemental Figure 2. Copy number profiles across all samples.**

(**A**) ASCAT[3] copy number profiles for all samples showing $\log_2$ ratio (LogR) and B-allele frequency (BAF) across chromosomes. LogR values near zero indicate diploid copy number; deviations indicate gains (positive) or losses (negative). BAF values near 0.5 indicate balanced heterozygosity; values near 0 or 1 indicate loss of heterozygosity or allelic imbalance. NAF and CN tissues maintain diploid genomes across all patients, while tumors display variable chromosomal alterations.

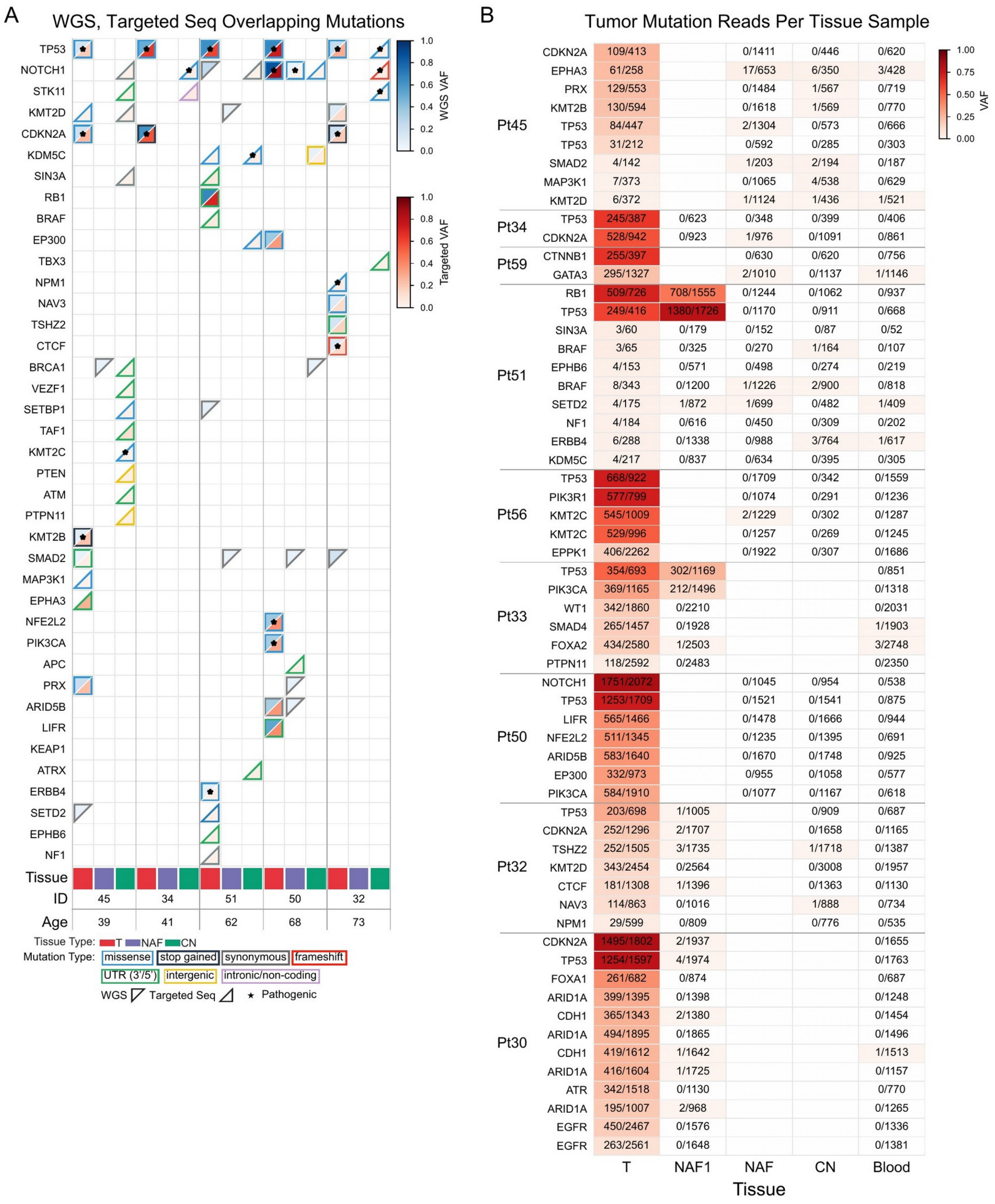

**Supplemental Figure 3. Integration of whole genome and targeted sequencing data.**

(**A**) VAF of driver gene mutations detected by whole genome sequencing (top left, blue scale) and targeted sequencing (bottom right, red scale) across matched tissue samples from five patients.

(**B**) Detection of tumor-associated mutations in matched tissues using targeted sequencing. To evaluate whether NAF clones derive from adjacent tumors, we identified reads supporting each tumor-identified mutation in matched NAF1 (samples with NAF close to tumor, dissected with LCM under guidance of pathologist), NAF, CN, and blood samples. Numbers indicate mutant reads/total reads at each locus; color intensity reflects VAF.

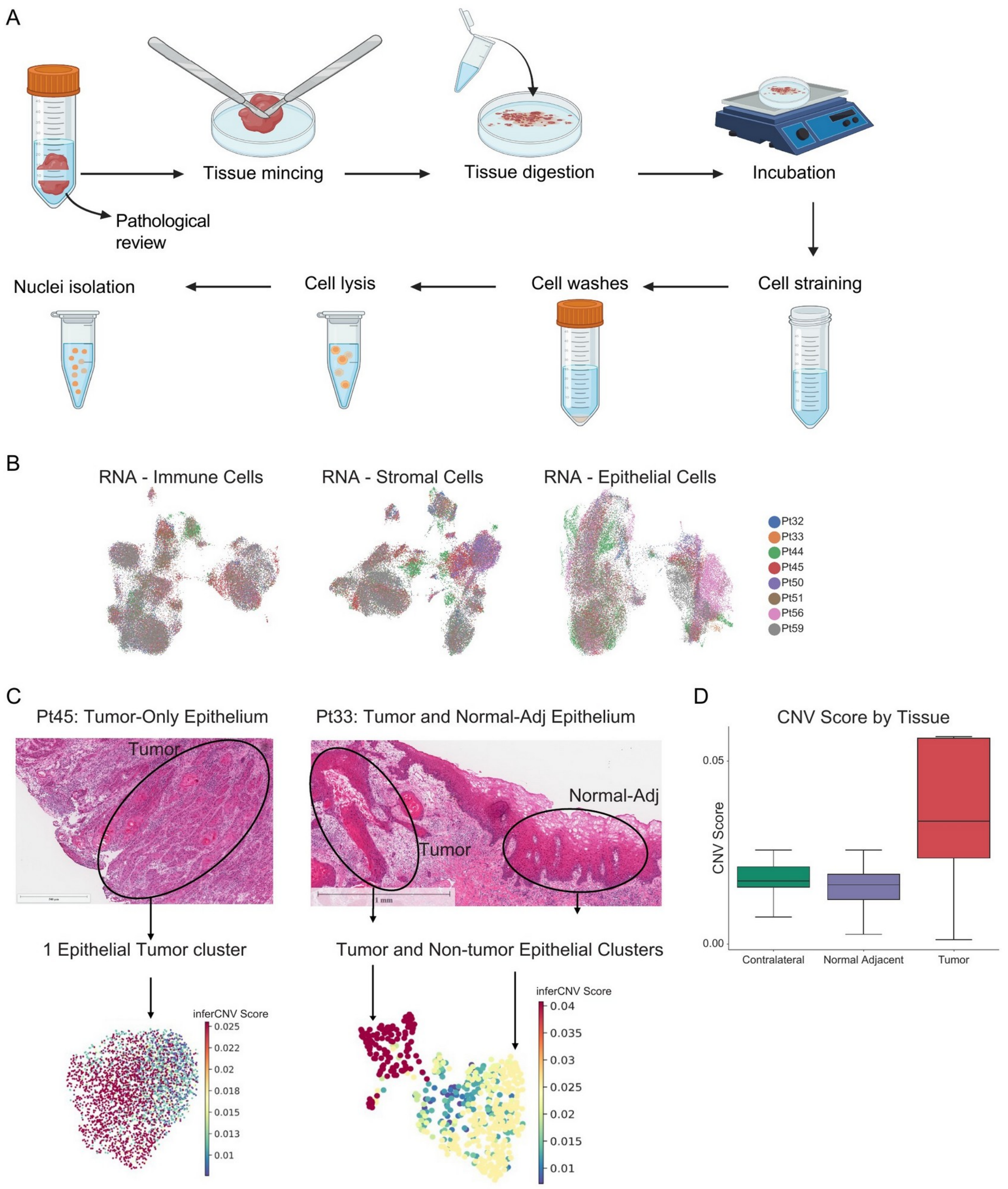

**Supplemental Figure 4. Single-nucleus multiome data generation and tumor cell annotation.**

(**A**) Overview of optimized protocol to generate single-nucleus multiomic data from patient tissue samples (see **Methods**).

(**B**) RNA UMAPs of immune, stromal, epithelial cells colored by patient.

(**C**) InferCNV analysis distinguishes tumor from non-tumor epithelial cells. (Left) Patient 45, whose tissue sample contained only tumor epithelium per pathological review, shows a single epithelial cluster with high inferCNV scores. (Right) Patient 33, whose tissue sample contained both tumor and normal-adjacent epithelium on pathological review, shows two distinct epithelial clusters: tumor cells with high inferCNV scores and non-tumor cells with low scores.

(**D**) Box plot of inferCNV scores across epithelial cell tissue categories (contralateral, normal-adjacent, and tumor). Tumor-derived epithelial cells exhibit markedly higher CNV scores compared to normal-adjacent and contralateral epithelial cells, confirming the ability of inferCNV to distinguish malignant from non-malignant epithelium.

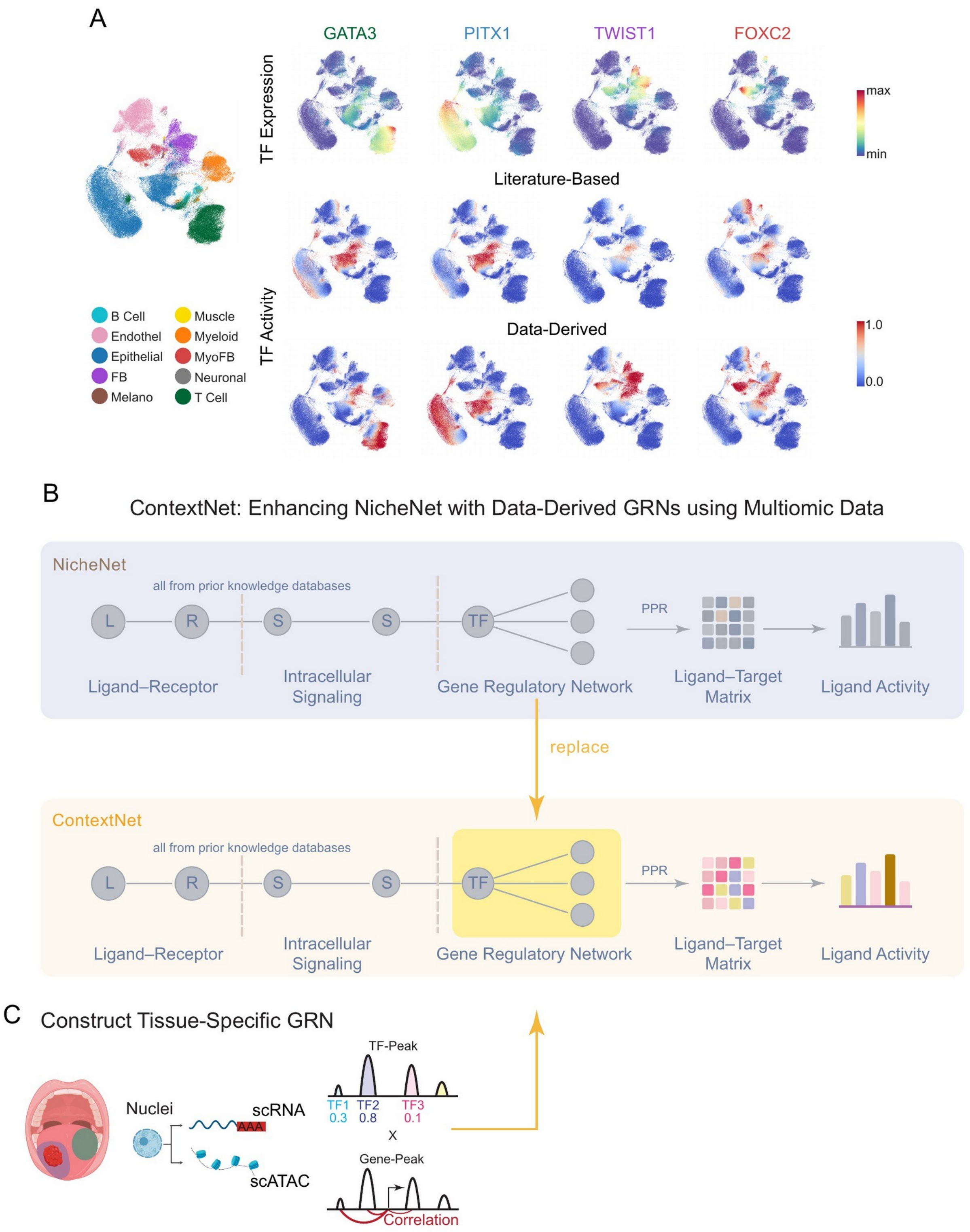

**Supplemental Figure 5. ContextNet approach and data-derived TF-gene regulatory network.**

(**A**) Data-derived GRN specificity across additional transcription factors. For each TF (GATA3, PITX1, TWIST1, FOXC2), gene expression is shown at the top, TF activity from the literature-based networks in the middle and TF activity inferred from the data-derived gene regulatory networks at the bottom.

**(B, C)** Schematic comparing NicheNet and ContextNet approaches.

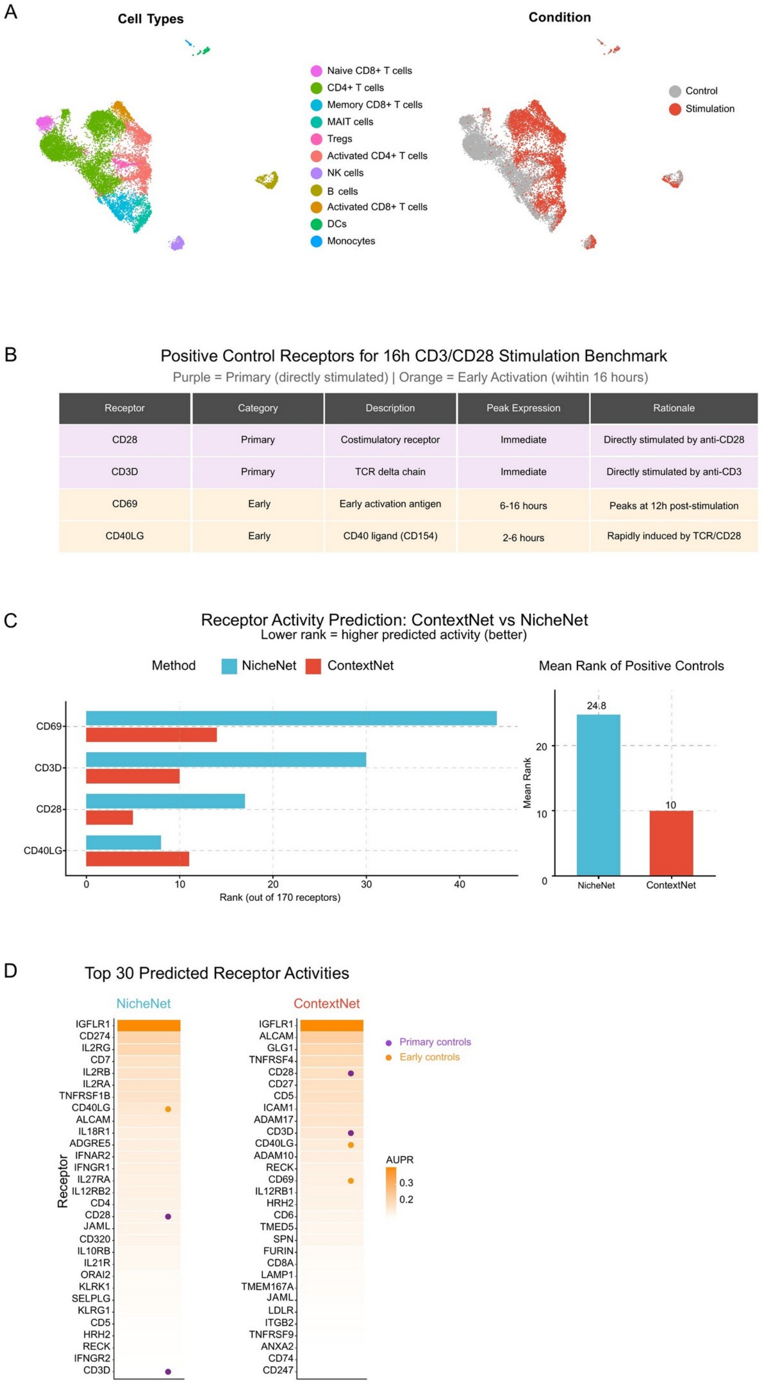

**Supplemental Figure 6. ContextNet improves prediction of T cell activation markers compared to literature-based NicheNet.**

(**A**) UMAP of DOGMA-seq data from stimulated and unstimulated human PBMCs, colored by cell type (left) and condition, stimulated with anti-CD3/CD28 versus unstimulated control[4] (right).

(**B**) Positive control genes used in benchmarking: known early T cell activation markers whose protein expression increases upon TCR stimulation.

(**C**) Predicted receptor activity ranks for positive controls; lower rank indicates higher predicted activity. ContextNet achieves lower mean rank across all positive controls.

(**D**) Top 30 predicted receptors for each method. ContextNet ranks more positive controls in top positions.

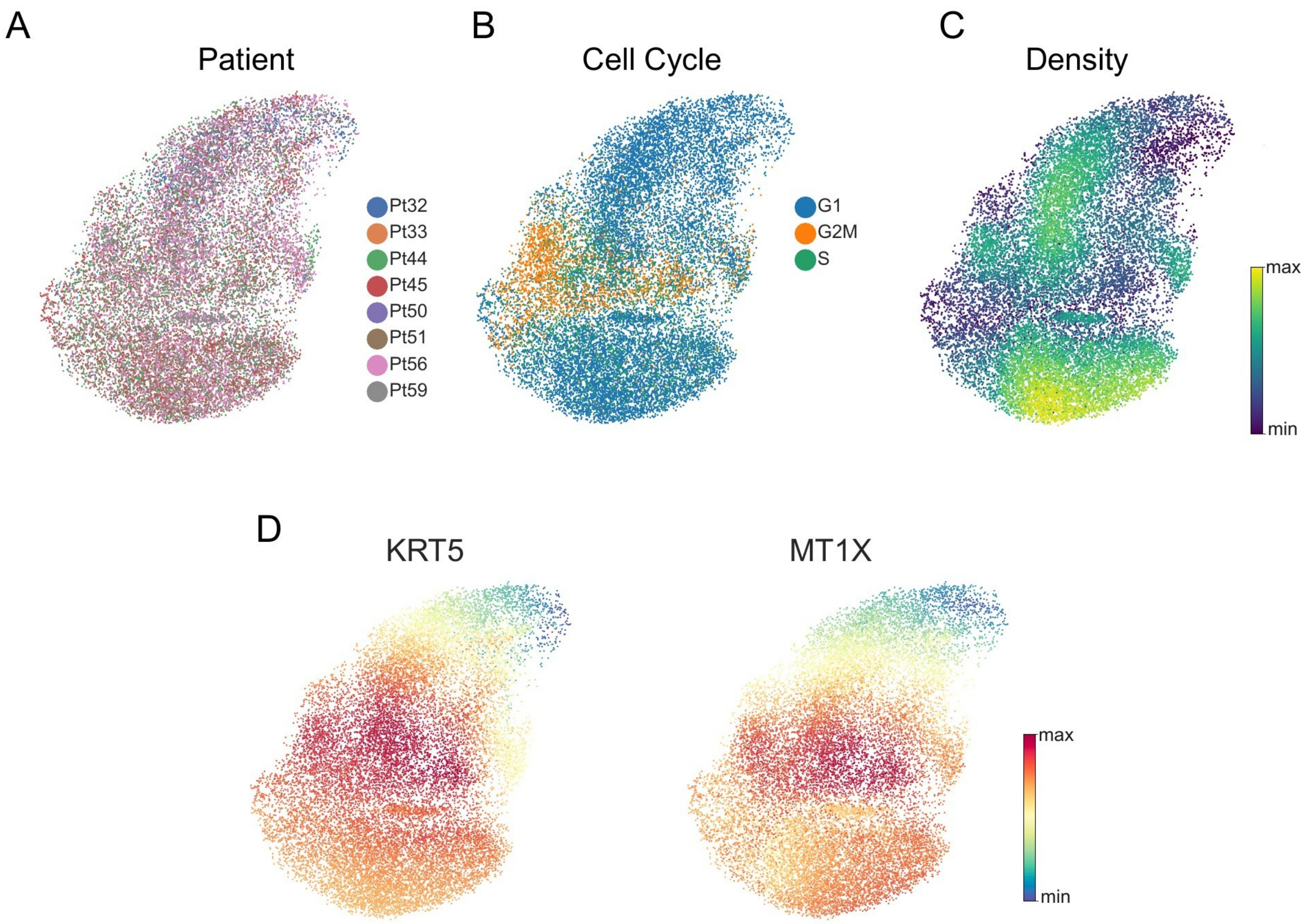

**Supplemental Figure 7. Characterization of contralateral normal epithelium.**

(**A**) UMAP of contralateral epithelial cells colored by patient.

(**B**) UMAP colored by cell cycle phase.

(**C**) UMAP colored by Mellon density.

(**D**) UMAP visualization of parabasal marker expression: *KRT5* and *MT1X*.

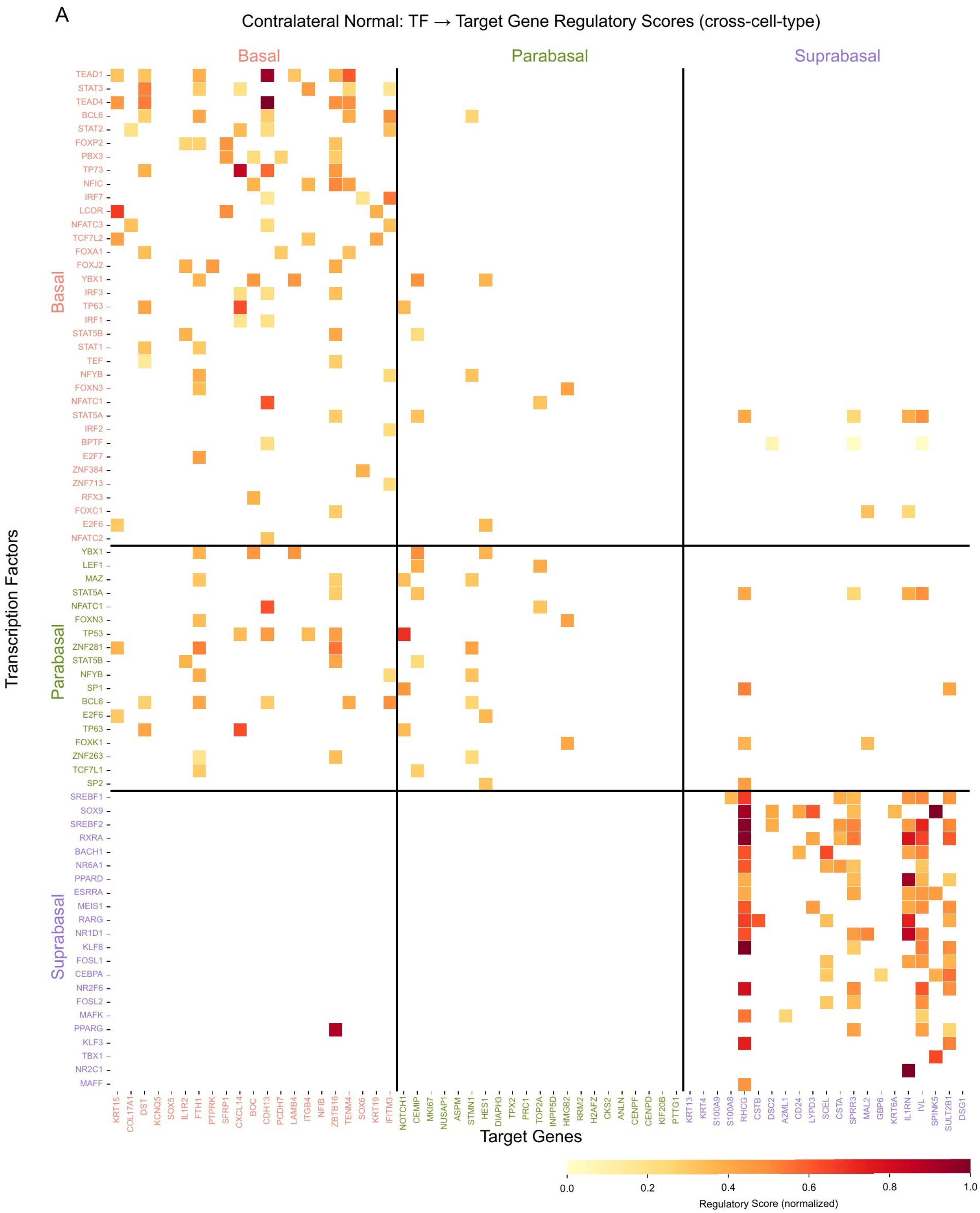

**Supplemental Figure 8. Context-specific transcription factor-target gene regulatory scores in contralateral normal epithelium.**

(**A**) Heatmap of regulatory scores linking transcription factors (rows) to target genes (columns), partitioned by receiver cell compartment (basal, parabasal, suprabasal). Regulatory scores were derived from the context-specific gene regulatory network constructed from paired RNA and ATAC data (see **Methods**). Transcription factors are grouped by the compartment in which they show highest motif accessibility. Color intensity reflects normalized regulatory weight.

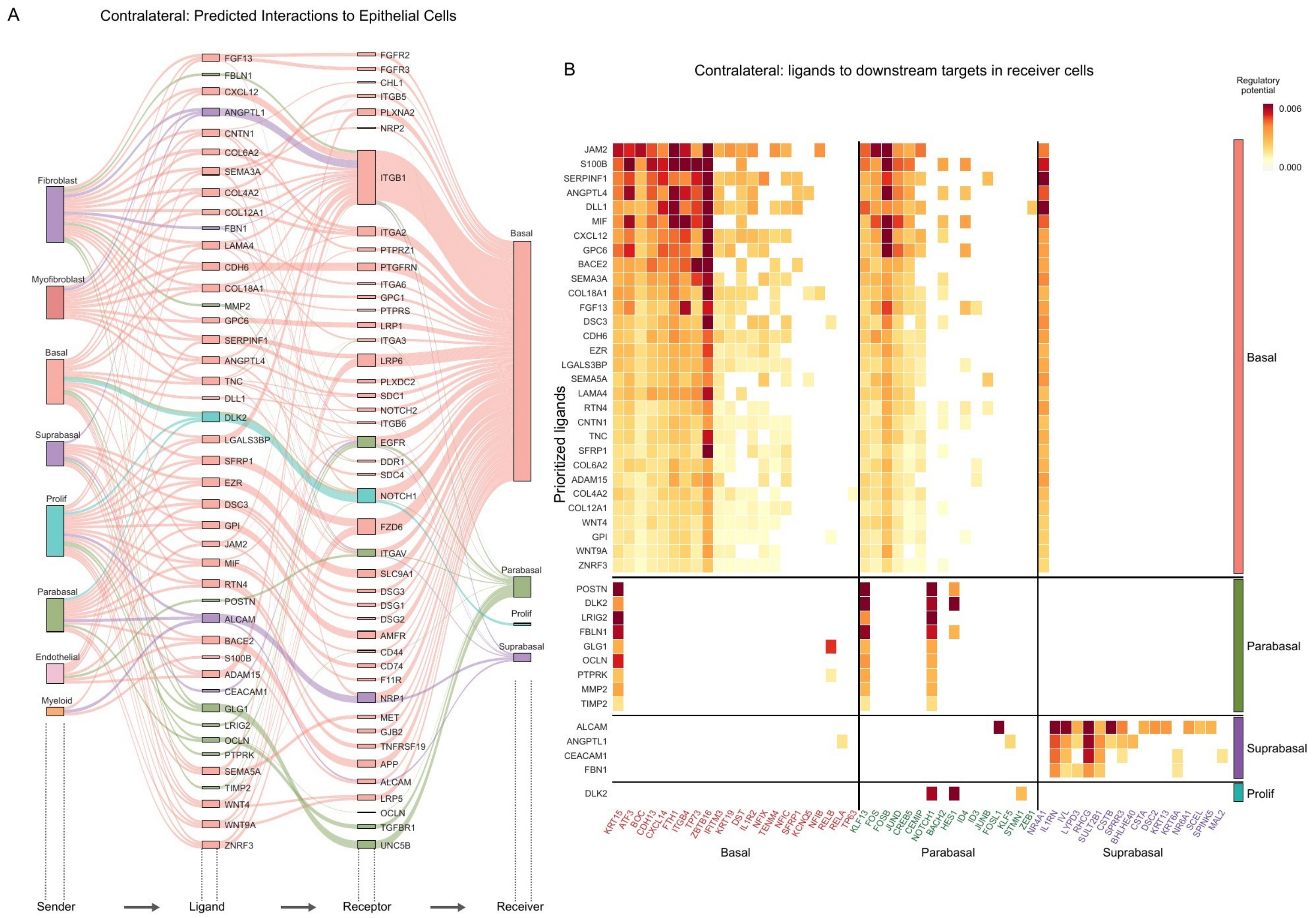

**Supplemental Figure 9. Ligand-target gene regulatory potential in contralateral normal epithelial compartments.**

**(A)** Cell-cell communication analysis using ContextNet. Sankey diagram depicting ligand-receptor signaling flow from sender cell types (left) through ligands and receptors to receiver epithelial compartments (right). Edge width reflects a composite score integrating ligand expression in receiver cells and predicted regulatory activity on receiver cells; edge color indicates receiver cell type.

(**B**) Heatmaps showing predicted regulatory potential of prioritized ligands (rows) on target genes (columns) for the different contralateral epithelial compartments, inferred by ContextNet. Color intensity reflects regulatory potential score. Target genes are both ordered by compartments and then total regulatory potential across ligands.

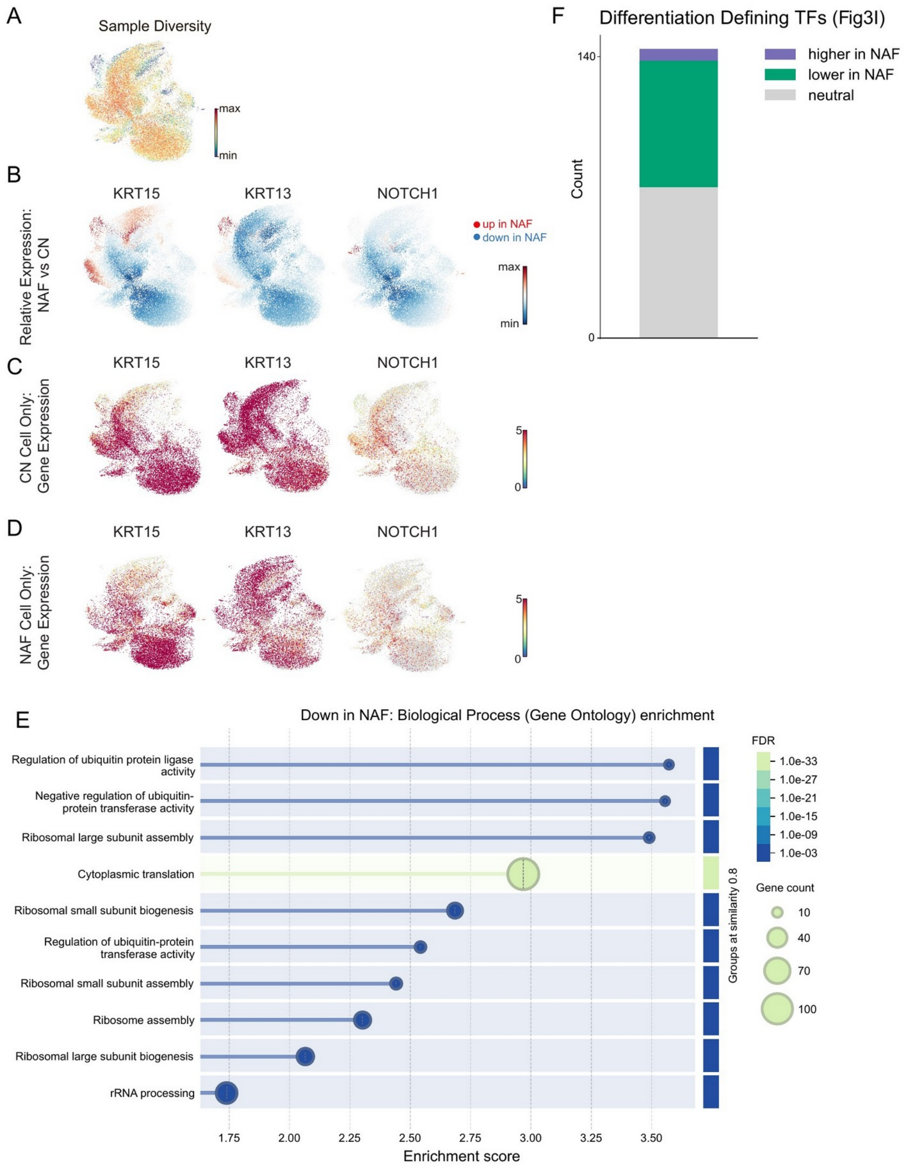

**Supplemental Figure 10. Extended characterization of epithelial transcriptional changes in NAF versus CN.**

(**A**) RNA UMAP colored by sample diversity calculated by neighborhood entropy across co-embedded NAF (normal adjacent field) and CN (contralateral normal) epithelial cells. High scores indicate many different patient samples represented in that region. Low scores indicate fewer samples represented in the region.

(**B**) UMAP colored by expression changes of *KRT15*, *KRT13* and *NOTCH1* between NAF and CN; Red indicates higher expression in NAF, blue indicates lower expression in NAF.

(**C**) Gene expression of *KRT15*, *KRT13* and *NOTCH1* in CN epithelial cells only.

(**D**) Gene expression of *KRT15*, *KRT13* and *NOTCH1* in NAF epithelial cells only.

(**E**) STRINGdb[5] Gene Ontology (Biological Process) enrichment analysis of genes significantly downregulated in NAF. Dot size indicates gene count; color indicates FDR.

(**F**) Bar chart showing the distribution of differentiation-defining transcription factors (identified in **Fig. 3J**) by differential chromVAR accessibility between NAF and CN: higher in NAF (purple) and lower in NAF (teal).

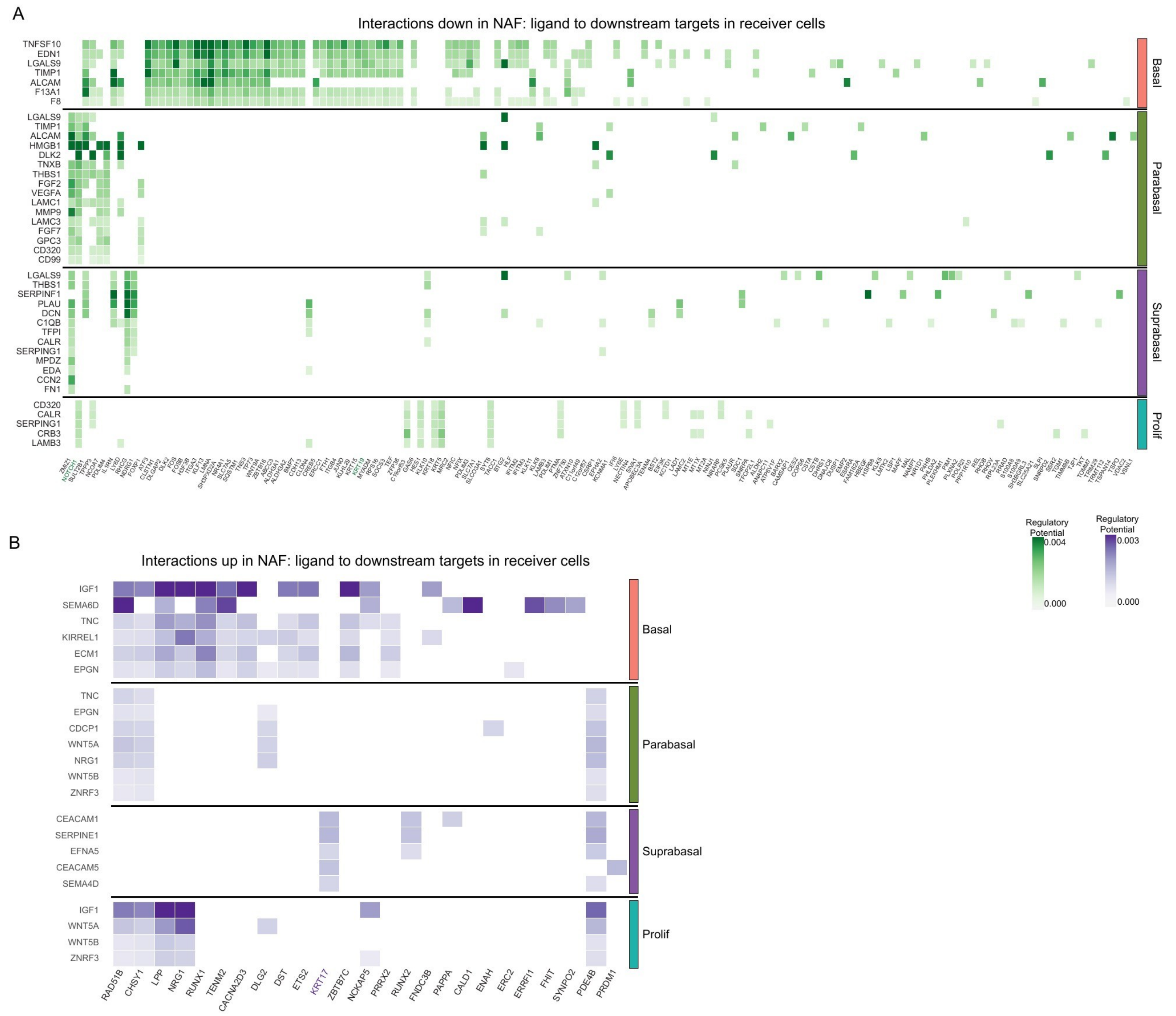

**Supplemental Figure 11. ContextNet ligand-target analysis by epithelial compartment.**

(**A**) Heatmap of predicted regulatory potential of prioritized ligands (rows) on predicted target genes (columns) for interactions downregulated in NAF (normal-adjacent field) relative to contralateral normal (CN) in all epithelial compartments.

(**B**) Heatmap of predicted regulatory potential of prioritized ligands (rows) on predicted target genes (columns) for interactions upregulated in NAF (normal-adjacent field) relative to contralateral normal in all epithelial compartments.

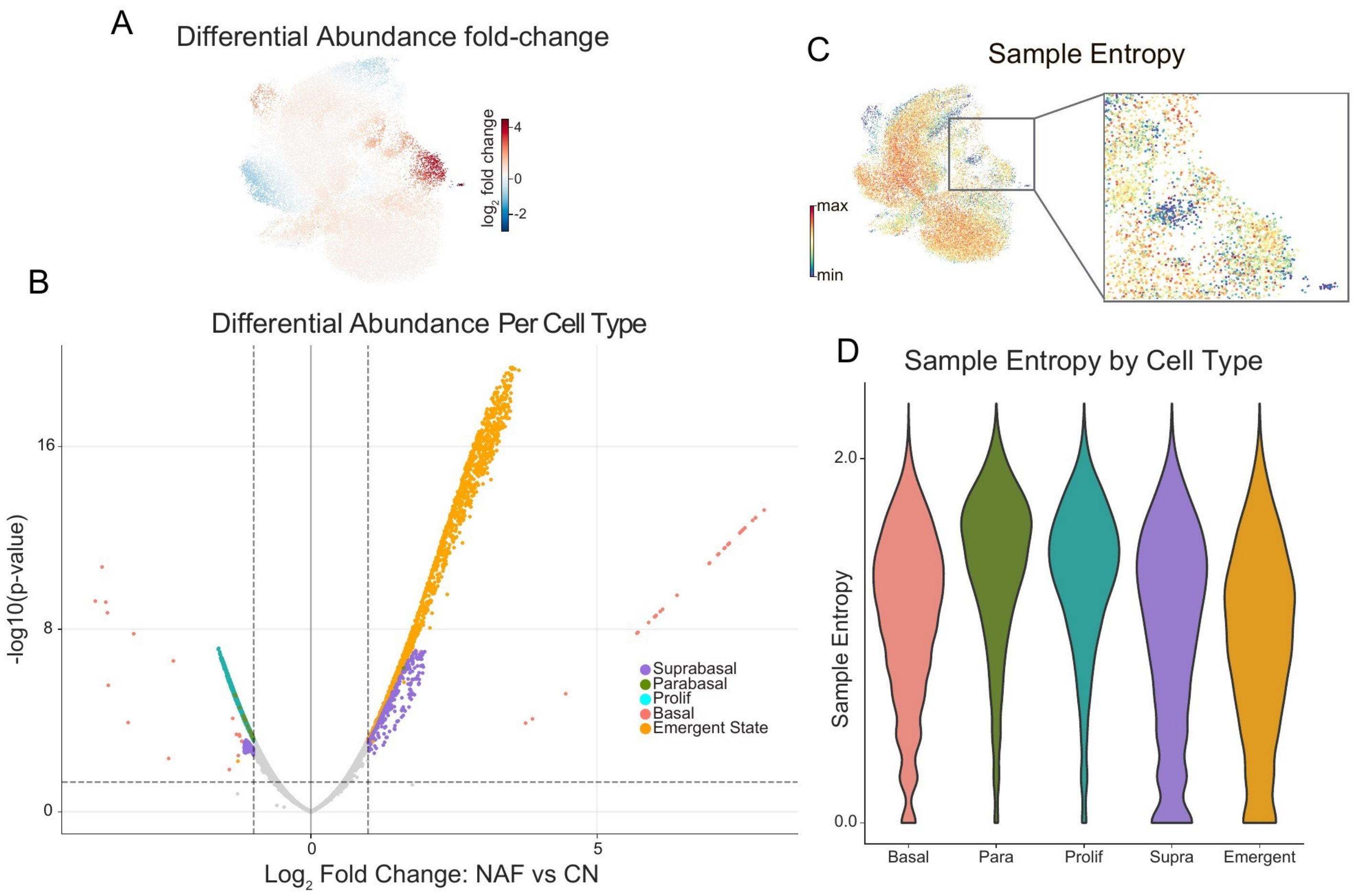

**Supplemental Figure 12. Differential abundance analysis of NAF versus CN epithelial cells.**

(**A**) RNA UMAP colored by Kompot differential abundance significance; color indicates $log_2$ fold change between NAF and CN.

(**B**) Volcano plot of differential abundance per cell type, showing $log_2$ fold change (CN to NAF) versus $-log_{10}$ (p-value). Points are colored by cell type.

(**C**) UMAP showing sample diversity calculated by neighborhood entropy across co-embedded NAF and CN epithelial cells with zoomed in view of emergent cell region with high differential abundance.

**(D)** Violin plot of neighborhood entropy across epithelial subtypes (basal, parabasal, proliferating, suprabasal, and emergent state) in co-embedded NAF and CN cells.

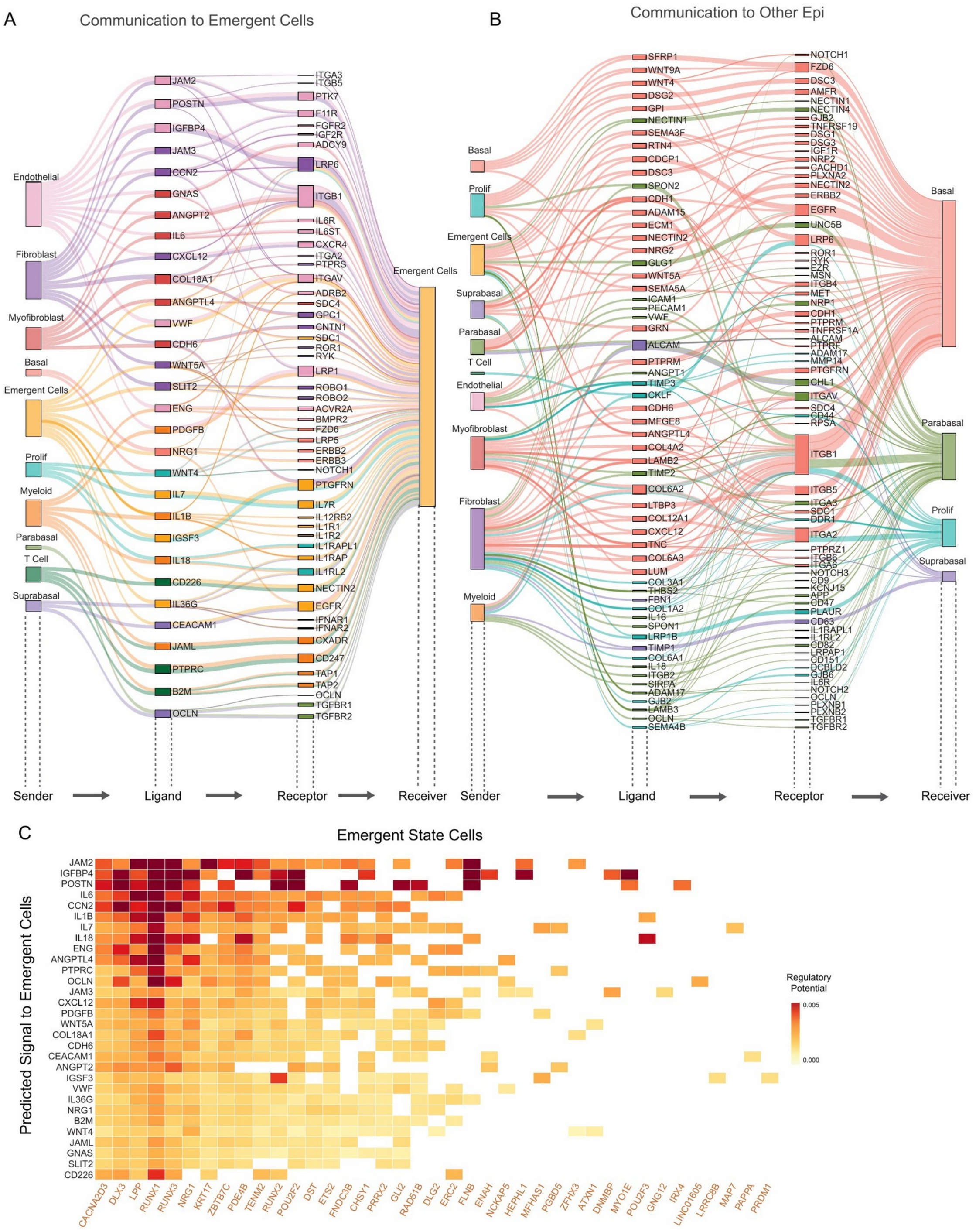

**Supplemental Figure 13. ContextNet ligand-target regulatory potential for emergent state cells.**

(**A**) ContextNet cell-cell communication analysis showing ligand-receptor signaling flow directed toward emergent state compartments (yellow). Edge width reflects composite regulatory score; edge color indicates receiver cell type.

(**B**) ContextNet results for other epithelial cell types in NAF.

(**C**) Heatmap of predicted regulatory potential of prioritized ligands (rows) on predicted target genes (columns) for the emergent state compartment. Color intensity reflects regulatory potential score.

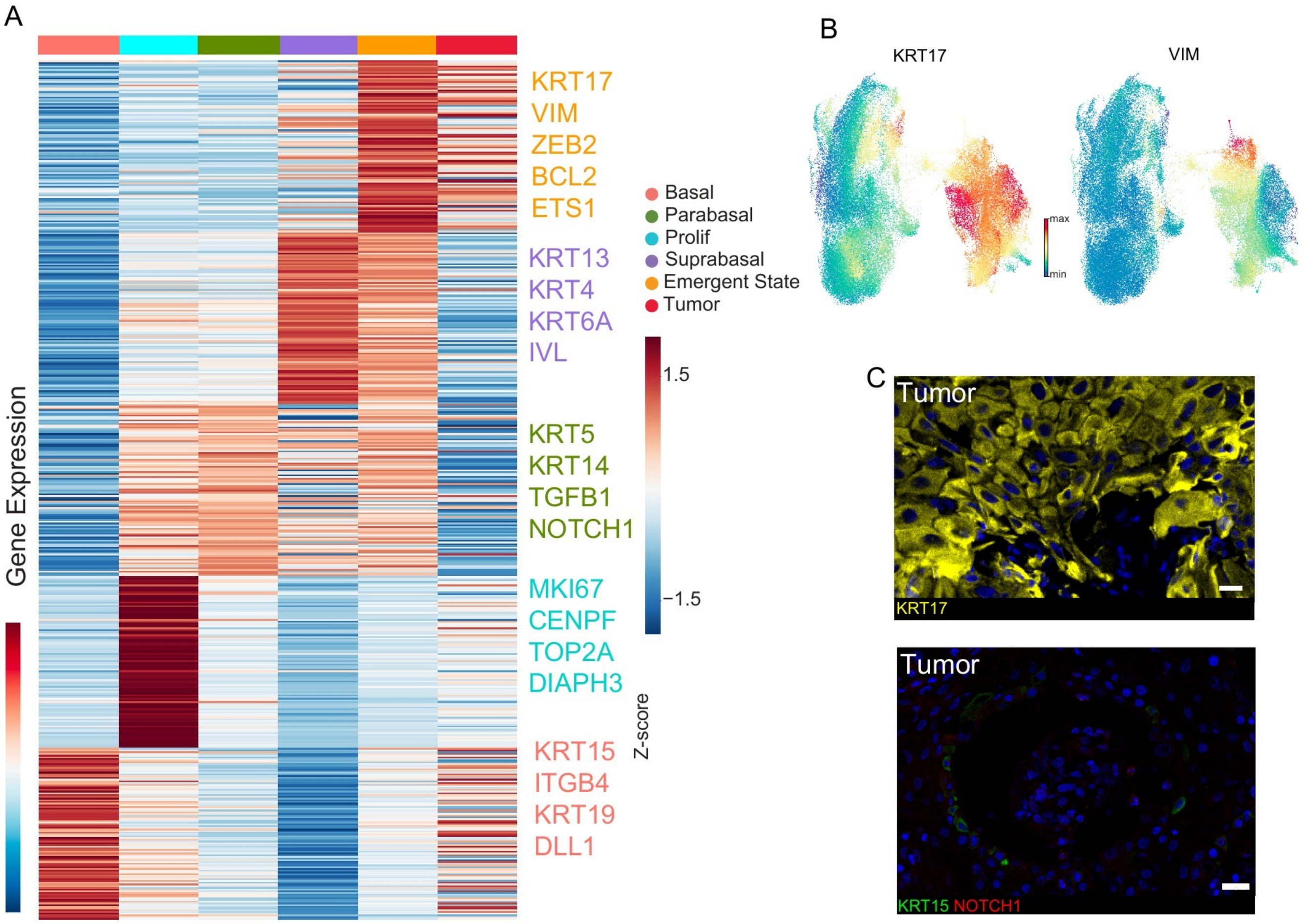

**Supplemental Figure 14. Emergent state markers in the context of tumor.**

(**A**) Heatmap of differentially expressed genes in NAF tissues, as **Fig. 5E**, and tumor cells.

(**B**) RNA UMAP of all epithelial cells including tumor colored by *KRT17* (left) and *VIM* (right) expression.

(**C**) Immunofluorescence staining of tumor tissue for KRT17 (yellow, top) and KRT15 (green)/NOTCH1 (red) (bottom). Nuclei stained with DAPI (blue). KRT15 and NOTCH1 image was taken using acquisition parameters as in **Fig. 4H**. Scale bars indicate 20 µm.

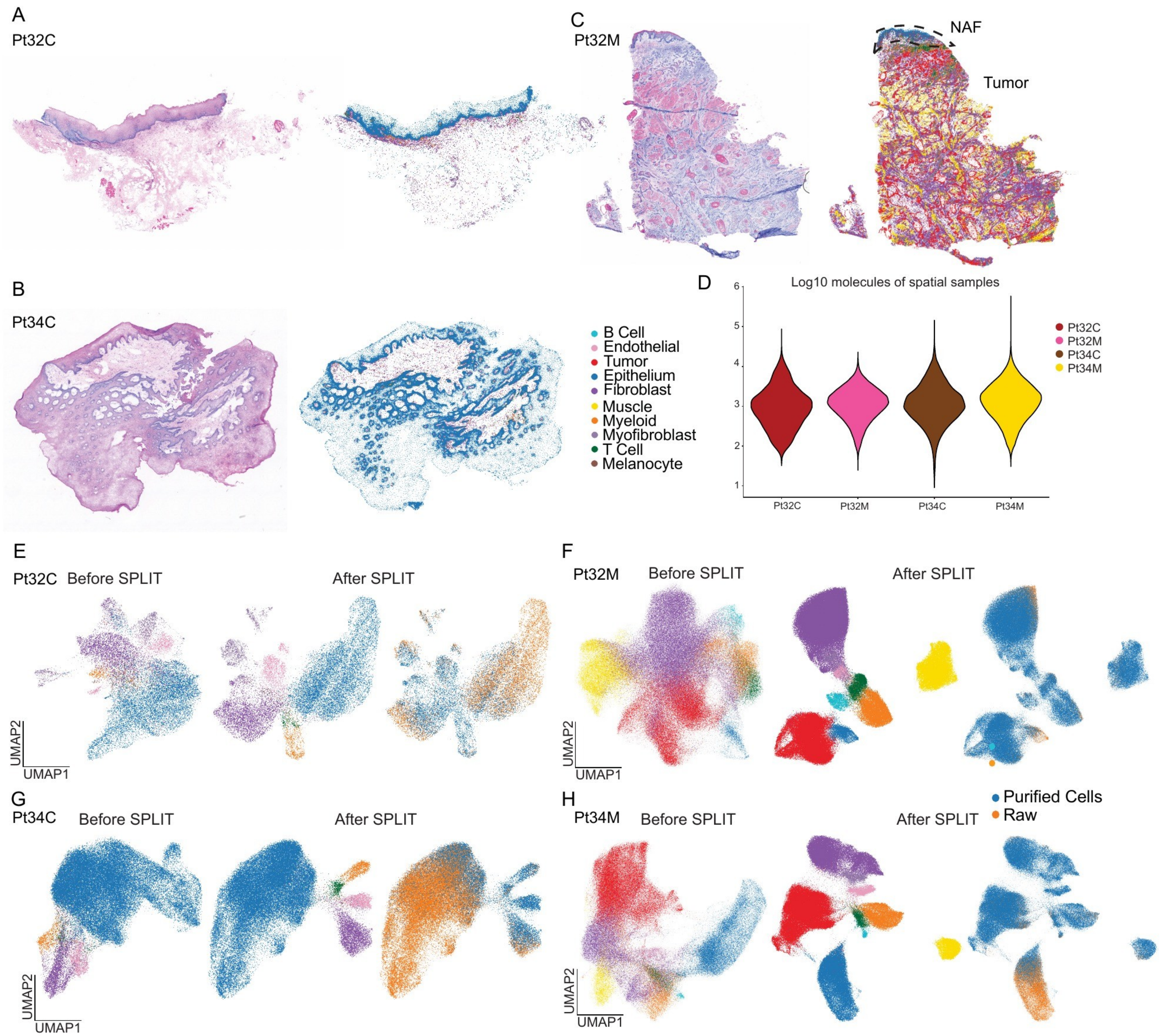

**Supplemental Figure 15. Spatial sample overview and preprocessing.**

(**A-C**) Spatial sample overview with H&E staining on the left and spatial cell-type annotation on the right: (**A**) Pt32C, (**B**) Pt34C, and (**C**) Pt34M, with the NAF region outlined by a dotted boundary. C: Contralateral, M: Mixed NAF/tumor.

(**D**) Violin plot of log10 molecular counts across the four spatial samples.

(**E-H**) SPLIT[6] purification results for each spatial sample: (**E**) Pt32C, (**F**) Pt32M, (**G**) Pt34C, and (**H**) Pt34M. Left: UMAP before SPLIT purification colored by cell types. Middle: UMAP after SPLIT purification colored by cell types. Right: UMAP after SPLIT purification highlight the cells purified by SPLIT.

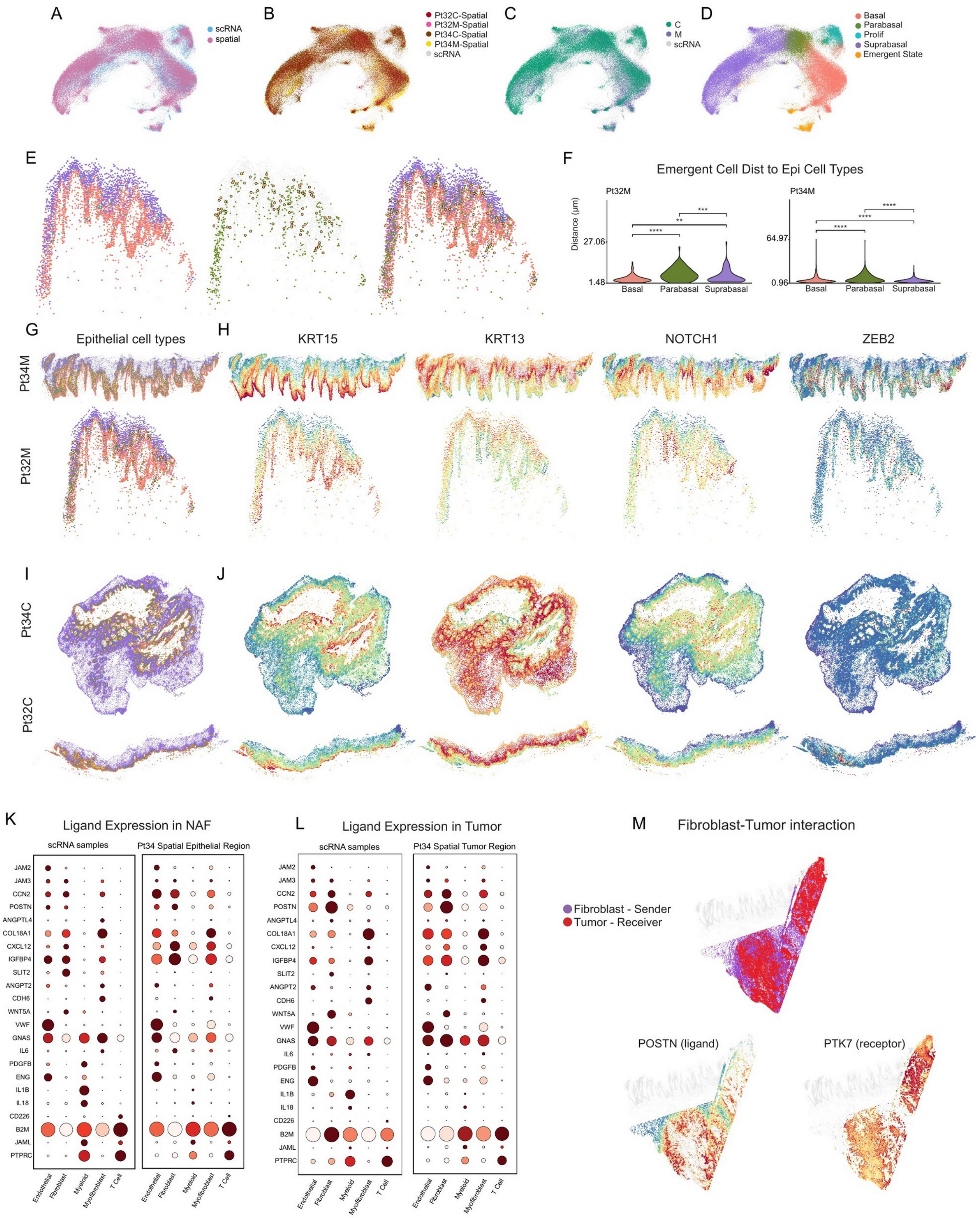

**Supplemental Figure 16. snRNA-seq/spatial integration, epithelial subtype mapping, and ContextNet ligand signals.**

(**A**) UMAP of epithelial cells generated by integration of snRNA-seq and spatial data, colored by data source. Contralateral (C) and normal-adjacent field (NAF) epithelial cells were used.

(**B**) UMAP in (A), colored by spatial sample.

(**C**) UMAP in (A), colored by tissue site.

(**D**) UMAP in (A), colored by cell types. Cell types were transferred from snRNA-seq to spatial data.

(**E**) Pt32 NAF spatial map showing basal, suprabasal, and dividing cells on the left, parabasal and emergent cells in the middle, and all epithelial subtypes on the right.

(**F**) Distance from emergent cells to other epithelial subtypes in the mixed samples.

(**G**) Spatial maps of epithelial subtypes in the following order from to bottom: NAF region from the Pt34 mixed sample, NAF region from the Pt32 mixed sample.

(**H**) Spatial maps from (G) colored by expression of KRT15, KRT13, NOTCH1 and ZEB2. (I) Spatial maps of epithelial subtypes in the following order from to bottom: NAF region from the Pt34 contralateral section, Pt32 contralateral section.. (**J**) Spatial maps from (I) colored by expression of KRT15, KRT13, NOTCH1 and ZEB2. (**K-L**) Dot plots of ContextNet-predicted ligands signaling to emergent cells. Left, ligand expression in snRNA-seq data; right, ligand expression in Pt34 spatial data. Dot size indicates the percentage of cells expressing the ligand, and color indicates mean scaled expression. (**K**) Ligand expression in NAF. (**J**) Ligand expression Tumor. (**M**) Spatial maps of POSTN ligand and PTK7 receptor expression in tumor, showing that the ContextNet-predicted POSTN-PTK7 signal to transitioning cells is also present in tumors.

Supplemental Tables

**Supplemental Table S1. Clinical and demographic data for 23 patients with HPV-negative oral squamous cell carcinoma (OSCC).**

Columns include patient ID, age at surgery, sex, anatomical site of the primary tumor, TNM staging, HPV status, neoadjuvant and adjuvant therapy history, tobacco and substance use history, alcohol use, and recurrence status.

**Supplemental Table S2. Summary of tissue samples and assays performed for each patient.**

Columns include patient ID, tissue type (B, blood; T, tumor; CN, contralateral normal mucosa; NAF, normal-appearing tissue adjacent to the tumor field; M, margin), assays conducted (whole-genome sequencing, WGS; targeted sequencing panel; single-cell multiome, scMultiome; spatial transcriptomics), and distance of the sampled tissue from the tumor margin where available.

**Supplemental Table S3. Whole-genome sequencing mutation summary for 15 tissue samples across 5 patients.**

For each sample, the table reports total variant counts (SNVs and INDELs), mutational burden (mutations per megabase), variant impact categories (HIGH, MODERATE, LOW, MODIFIER), mutation types (missense, synonymous, stop gained, frameshift, splice), genomic context (intergenic, intronic, protein-coding), variant allele frequency (VAF) statistics (mean, median, standard deviation), clonality classification (clonal VAF > 0.3, subclonal 0.1–0.3, low VAF < 0.1), caller concordance, number of genes affected, transition/transversion ratio, single-nucleotide substitution spectrum (C>A, C>G, C>T, T>A, T>C, T>G), AlphaMissense pathogenicity predictions (likely pathogenic, ambiguous, likely benign), and sequencing depth.

**Supplemental Table S4. Copy number variation (CNV) segments derived from whole-genome sequencing for 15 samples across 5 patients.**

Each row represents a genomic segment with columns for sample ID, chromosome, start and end coordinates (hg38), major allele copy number, minor allele copy number, and total copy number.

**Supplemental Table S5. List of 127 genes included in the targeted sequencing panel used for cancer-associated mutational profiling of tissues.**

**Supplemental Table S6. Marker genes used for cell type annotation of single-cell multiome data.**

The table is divided into two sections: (1) general cell type markers for epithelial, fibroblast, myofibroblast, muscle, B cell, myeloid, T cell, endothelial, neuronal, and melanocyte populations; and (2) epithelial subtype markers distinguishing basal, proliferating (dividing), and suprabasal states.

**Supplemental Table S7. Cell-level metadata for 145,769 cells from single-cell multiome (joint RNA and ATAC) profiling across 8 patients.**

Columns include cell barcode, tissue category (Contralateral, Tumor, or Normal Adjacent Field), patient ID, and assigned cell type annotation.

**Supplemental Table S8. Differentially expressed genes in contralateral normal (CN) epithelial cells across epithelial subtypes.**

Columns include gene name, $log_2$ fold change, Benjamini-Hochberg adjusted P value, and epithelial cell type (suprabasal, dividing, parabasal, or basal). A total of 761 differentially expressed genes are reported.

**Supplemental Table S9. Results of Kompot differential gene expression analysis comparing epithelial cells from contralateral normal (CN) versus normal adjacent field (NAF) tissues at the RNA level.**

For each of 16,524 genes, the table reports the mean $log_2$ fold change, Mahalanobis distance, local false discovery rate, differential expression status, and direction of change (upregulated in normal adjacent, downregulated in normal adjacent, or not significant). A total of 1,711 genes were identified as differentially expressed.

**Supplemental Table S10. Results of Kompot differential chromatin accessibility analysis comparing epithelial cells from contralateral normal (CN) versus normal adjacent field (NAF) tissues at the ATAC level.**

For each of 421 transcription factor motifs, the table reports the mean chromVAR expression difference, Mahalanobis local false discovery rate, differential accessibility status, and direction of change (higher in CN, lower in NAF, or neutral). A total of 156 transcription factor motifs showed differential accessibility.

**Supplemental Table S11. Differentially expressed genes between contralateral normal (CN) and normal adjacent field (NAF) epithelial cells, stratified by epithelial subtype.**

Columns include gene name, $log_2$ fold change, Benjamini-Hochberg adjusted P value, and cell type (emerging state, suprabasal, parabasal, dividing, or basal). A total of 500 differentially expressed genes are reported.

**Supplemental Table S12. Transcription factor activity scores for 150 transcription factors across epithelial subtypes, with enrichment in the emerging state population.**

Columns show each transcription factor and its mean activity score in basal, dividing, parabasal, suprabasal, and emerging state cells, enabling identification of transcription factors specifically activated in the emerging epithelial state found in normal adjacent field tissue.

**Supplemental Table S13. Primary and secondary antibodies used for immunofluorescence validation experiments.**

For each antibody, the table lists the target antigen, manufacturer, catalogue number, and working dilution.

References

1. Islam, S. M. A. *et al.* Uncovering novel mutational signatures by de novo extraction with SigProfilerExtractor. *Cell Genomics* **2**, 100179 (2022).
2. Alexandrov, L. B. *et al.* The repertoire of mutational signatures in human cancer. *Nature* **578**, 94–101 (2020).
3. Van Loo, P. *et al.* Allele-specific copy number analysis of tumors. *Proceedings of the National Academy of Sciences* **107**, 16910–16915 (2010).
4. Mimitou, E. P. *et al.* Scalable, multimodal profiling of chromatin accessibility, gene expression and protein levels in single cells. *Nat. Biotechnol.* **39**, 1246–1258 (2021).
5. Szklarczyk, D. *et al.* STRING v11: protein–protein association networks with increased coverage, supporting functional discovery in genome-wide experimental datasets. *Nucleic Acids Res.* **47**, D607–D613 (2019).
6. Bilous, M. *et al.* Resolving sensitivity, specificity and signal contamination in Xenium spatial transcriptomics. *Nat. Methods* **23**, 1152–1162 (2026).
